# Antimalarial mass drug administration in large populations and the evolution of drug resistance

**DOI:** 10.1101/2021.03.08.434496

**Authors:** Tran Dang Nguyen, Thu Nguyen-Anh Tran, Daniel M. Parker, Nicholas J White, Maciej F Boni

**Author notes:** **Correspondence** Maciej F Boni, 252W Millennium Science Complex, Department of Biology, Pennsylvania State University, University Park, PA, 16802, USA. contributed equally (Tran Dang Nguyen, Thu Nguyen-Anh Tran).

## Abstract

Mass drug administration (MDA) with antimalarials has been shown to reduce prevalence and interrupt transmission in small populations, in populations with reliable access to antimalarial drugs, and in populations where sustained improvements in diagnosis and treatment are possible. Effective MDA eliminates drug-resistant parasites which has the long-term benefit of extending the useful therapeutic life of first-line therapies for all populations, not just the focal population where MDA was carried out. However, in order to plan elimination measures effectively, it is necessary to characterize the conditions under which failed MDA could exacerbate resistance. We use an individual-based stochastic model of *Plasmodium falciparum* transmission to evaluate this risk in large populations (>40K) where access to antimalarial treatments may not be uniformly high and where re-importation of drug-resistant parasites may be common. We find that drug-resistance evolution can be accelerated by MDA when all three of the following conditions are met: (1) strong genetic bottlenecking that falls short of elimination, (2) re-importation of resistant genotypes, and (3) continued selection pressure during routine case management post-MDA. Accelerated resistance levels are not immediate but follow the rebound of malaria cases post-MDA, if this is allowed to occur. Crucially, resistance is driven by the selection pressure during routine case management post-MDA and not the selection pressure exerted during the MDA itself. Second, we find that increasing treatment coverage post-MDA increases the probability of local elimination in low-transmission regions (PfPR < 2%) in scenarios with both low and high levels of drug-resistance importation. This emphasizes the importance of preparation and planning to ensure that MDA has a high probability of leading to elimination, and the necessity of supporting public health infrastructure to provide high coverage of diagnosis and treatment post-MDA.

## 1 INTRODUCTION

Global malaria burden dropped substantially between 2000 and 2015, prompting the World Health Organization to once again articulate the case for and evaluate the feasibility of global malaria eradication [1,2]. These efforts have been progressing via national-level malaria elimination plans in countries that are nearing elimination phase, and mass drug administration (MDA) will be one of the key public health tools deployed in regions or transmission pockets where complete elimination has proved otherwise difficult. In small populations and populations with low malaria prevalence, MDA implementations have proved successful at pushing infection numbers to zero [3,4]. In larger populations or certain high-prevalence scenarios, MDA can be followed by a rebound of malaria cases after the intended rounds of MDA have been completed. Rebounds of case numbers and malaria prevalence have been observed in the field months or years after the transmission-reducing effects of the MDA have worn off [5–11], an effect that is predicted by mathematical modeling analyses [12–14] if malaria control efforts are not intensified from the pre-MDA status quo. An important question in MDA application and follow-up is how to keep transmission levels low with the eventual goal of interrupting transmission permanently.

A second concern addressed here is the natural selection of drug-resistant genotypes during MDA [15,16]. Ideally, a therapy chosen for MDA would have high efficacy in patients and no prior history of use in the focal population, in order to minimize the probability of pre-existing drug resistance being selected for during the MDA. These measures may mitigate the risk, but the MDA’s core approach – to dose every resident in a population with antimalarial drugs whether parasitaemic or not – is necessarily accompanied by substantial selection pressure for drug resistance. Quantifying this risk is not straightforward as resistance frequencies can go up while prevalence is dropping, which is why it is critical to combine drug-resistance risks (net negatives) with likelihood of malaria elimination (a net positive) into a single unified analysis. In some scenarios, drug-resistant alleles can appear to fix while the parasite population is on the path to elimination. An evaluation of an MDA’s benefits and risks needs to account for drug-resistant allele frequencies, absolute numbers of individuals carrying resistant genotypes, the probability that elimination will occur before resistant genotypes fix in the target population, and the long-term number of treatment failures experienced in the population.

In small populations and low-prevalence scenarios, the balance of the field and modeling evidence shows that MDA leads to net positive outcomes with high likelihood of local elimination and little drug resistance risk. In this study, we evaluate drug-resistance risks associated with MDA programs deployed in large populations (>40K individuals) where prevalence may still be moderate to high and ACT adoption may be low; these scenarios are motivated by African epidemiological contexts where diagnosis, treatment, and ACT adoption are all lower than in Southeast Asia. Using a previously developed individual-based mathematical transmission model of *Plasmodium falciparum* [17], we explicitly model the small numbers of parasite positive individuals post-MDA and describe the conditions under which drift, selection, and migration can act synergistically to accelerate the fixation of drug-resistant genotypes. Finally, we outline epidemiological scenarios where MDA policies are likely to pose the most and least risk to near-term and long-term drug-resistance trends.

## 2 Methods

### 2.1 Model Description

We modified a previously published individual-based mathematical model of *P. falciparum* transmission and evolution [17]. The upgraded model (https://github.com/bonilab/malariaibm-MDA-2018WHOERG) includes a locus-based resistance framework, with drug-resistance phenotypes parameterized for the K76T locus (*pfcrt* gene), N86Y and Y184F loci (*pfmdr1* gene), C580Y locus (*kelch13* gene), copy number of *pfmdr1*, and copy number of *plasmepsin 2-3* genes. Drug resistance of a *P. falciparum* genotype in the simulation is fully described by these six genetic features. Each genotype-drug combination has assigned to it its own EC50 value and parasite killing rate value (*p*_max_), from which we derive the efficacy of each therapy/drug on each parasite genotype; see Supplementary Appendix 2 for the parametrization of the drug-by-genotype table used in these analyses.

Individuals newly infected by *P. falciparum* may experience symptoms (depending on their immune status), and 55% of symptomatic individuals (60% if <5) will seek and receive antimalarial treatment. At the beginning of the simulation (January 1 2022) it is assumed that among treatment-seeking individuals, 36% will receive artemether-lumefantrine (AL) [18] which is the first-line policy in our modeled scenario, and the remaining 64% will purchase SP (assumed efficacy of 40%), amodiaquine, chloroquine, or AL on the private market; these are typical values in many African settings. Assuming that ACT adoption continues to increase, we set an annual increase of 2% per year in ACT use, and the model’s 36% public-market use in 2022 increases to 76% in 2042 [18].

The new model allows for mass drug administration (MDA) with dihydroartemisinin-piperaquine (DHA-PPQ). Each individual in the population is assigned a probability of participating in a single round of MDA, as some individuals by the nature of their work, travel, or age will be less likely or more likely to be present when MDA team members are present and distributing treatments [19,20]. Each individual’s participation probability is drawn from a beta distribution with mean 0.75 for individuals aged 10-40, and mean 0.85 [21] for individuals aged <10 or >40; standard deviation of this beta distribution is set at 0.3 [22,23] for all ages.

As *P. falciparum* population sizes after the MDA may be quite low, we evaluate the hypothesis of importation rather than mutation presenting the primary drug-resistance risk. Importation is modeled as one new parasite coming into the population every 10 days; other importation scenarios are noted where appropriate. Thus, our standard importation scenario is intended to model a region or site where re-importation of drug resistance would be common and expected, in order to evaluate a situation perceived as having maximum risk of drug-resistance re-importation. The genotype of the imported parasite is chosen at random with equal probabilities for alleles and copy numbers for all loci. This means that during an MDA, an imported parasite has a 25% chance of being a double-resistant to both DHA and PPQ and a 50% chance of being resistant to exactly one of the drugs in the combination. Mutations do arise *de novo* [24] in the simulation, at all loci, only if the mutated phenotype carries an advantage in the present within-host drug environment (by being associated with lower drug efficacy). In a small minority of simulations, 580Y alleles may exist at low frequencies prior to the start of an MDA campaign due to early *de novo* mutation.

### 2.2 Scenarios Modeled

For this modeling exercise, populations of 40,000 and 300,000 are considered [25], and the planned MDA targets all individuals in the population, although only between 75% and 85% will be present for participation for any given round of MDA. Four rounds of MDA are administered five weeks apart; it takes two weeks to complete one round and the campaign is completed in 17 weeks. The start date for the first round is Jan 1 2022, and the subsequent rounds start on February 5, March 12, and April 16. The antimalarial used in each round of MDA is a three-day course of DHA-PPQ. Simulations are run for twenty years to investigate long-term patterns of drug resistance following a single MDA program run in the first year only.

As it has been raised in a number of MDA trials that improved coverage and increased treatment access are key to lowering prevalence and incidence [3,6,20,25,26], we implemented a model feature where improved treatment coverage (ITC) follows the MDA. In this scenario, the program’s infrastructure and improved knowledge on antimalarial use is leveraged to increase treatment coverage from 55%/60% (pre-MDA levels) to 80% of all symptomatic malaria cases treated (unless another percentage is noted); the increase occurs over a six-month period and remains constant for the reminder of the simulation. *P. falciparum* prevalence levels investigated ranged from 1% to 5%; higher prevalence scenarios (10%, 15%) did not lead to elimination and had qualitatively similar dynamics to a 5% setting. Initial genotypes in the population are 50% N86 and 50% 86Y at the N86Y locus of *pfmdr1*. All other loci are *pfcrt*-76T, *pfmdr1*-Y184 single copy, *kelch13*-C580, and single copy of *plasmepsin 2-3*; variation at the N86Y locus only is included due to the multitude of effects this locus has on resistance phenotypes to different drugs (Supplementary Appendix 2).

### 2.3 Outcomes

Allele frequency is computed using a weighting approach to account for individuals with multi- clonal falciparum infections. The frequency of an allele or genotype is computed as the weighted number of individuals carrying a certain allele divided by the total number of parasite-positive individuals. The weight of an individual in this sum is their number of parasite clones carrying the allele divided by the person’s total number of clones circulating in the blood.

The *kelch13* 580Y allele frequency is used as a proxy for artemisinin-resistant phenotypes in our analysis. Double copy of *plasmepsin*-*2* or *plasmepsin*-*3* genes is a proxy for PPQ resistance [27,28]. The *P. falciparum* blood-slide prevalence in 2-10 year-olds (PfPR_2-10_ [29]) is used to evaluate the success of MDA and any subsequent ITC. The time it takes for 580Y allele frequencies to reach 0.25 is used as a measure of how quickly artemisinin resistance reaches an irreversible course of fixation; this is a milestone at which it is too late to contain the spread of resistance. Differences among distributions are presented using Mann-Whitney (MW) or Kruskal-Wallis (KW) tests, treating multiple simulation runs as independent samples from the set of all evolutionary-epidemiological outcomes that are possible given a particular set of fixed assumptions.

## 3 Results

### 3.1 Effects of MDA on drug-resistance evolution

Implementation of high-coverage mass drug administration is followed by a drop in prevalence which results in a genetic bottleneck lasting from months to years following the MDA. When drug-resistance alleles are present or imported, whether at low or moderate frequencies, this bottleneck period is marked by substantial uncertainty and the near-term evolutionary outcome post-bottleneck will have low predictability (Figure 1). This is an unavoidable outcome of the action of random genetic drift in small populations, as successful MDA will reduce parasite population size and genetic diversity by forcing the parasite population through a genetic bottleneck [30–32]. Traditionally, the population genetics literature treats a bottleneck as a genotype-neutral event in which the expected reduction in population size for each allele is the same (i.e. no allele is preferentially selected), and the stochastic nature of the intervention results in randomly altered allele frequencies when comparing allelic distributions pre- and post-bottleneck. Under an MDA program there is one important difference, namely, that the cause of the population crash is non-neutral in its effects on the standing genetic variation in the parasite population as the MDA itself selects for drug-resistant alleles. Thus, an MDA can be viewed as a bottleneck created by negative viability selection. In the following analysis, we present both the population biological effects of MDA (reductions in malaria prevalence) and the population-genetic effects (changes in genotype distribution) resulting from the bottleneck following MDA.

**Figure 1.**
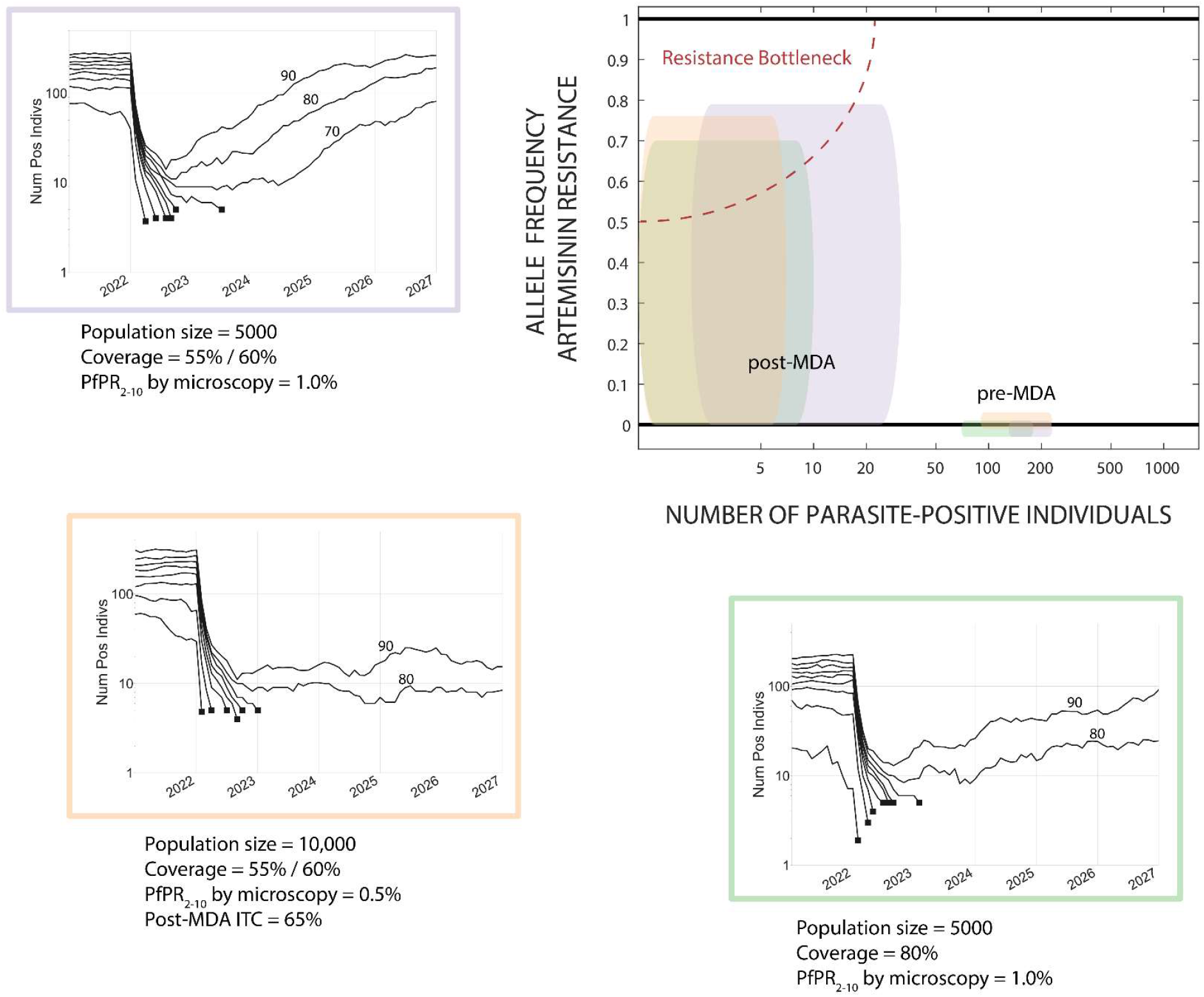
Schematic showing the relationship between prevalence and drug-resistance pre- and post-MDA in three selected small-population examples. The three scenario plots summarize 100 simulations under three rounds of MDA by showing the deciles (10th percentile to 90th percentile) of the simulation outcomes. The *y*-axis shows the number of parasite-positive individuals, including those that are not detectable by microscopy. Simulation trajectories that reach extinction/elimination (<5 parasite-positive individuals) end in a black square; a decile may not be shown if the number of parasite-positive individuals was <5 prior to the MDA. All examples include importation of drug-resistance of about two artemisinin-resistant genotypes per year. Elimination occurs for ∼70% of simulations in the scenarios shown, and for ∼80% of scenarios the number of parasite-positive individuals stays below 20 for at least three years. The scenarios chosen are boundary examples; in all three, probability of elimination increases with an extra round of MDA, lower importation rates of drug-resistance, lower pre-MDA prevalence, higher treatment coverage, or improved treatment coverage (ITC) post-MDA. The top-right panel shows the change in the parasite population profile from pre-to post-MDA. The box widths and heights show the inter-quartile range for the number of parasite-positive individuals (any level of parasitaemia) and the frequency of 580Y alleles, respectively; 580Y allele frequencies are zero pre-MDA. Parasite population sizes drop to the single digits or low double digits, but 580Y allele frequencies are unpredictable and can potentially be high. The upper-left corner of the schematic shows the ‘resistance bottleneck’ region where the number of infected individuals low, 580Y allele frequency is high, and the future path of drug resistance is unpredictable.

When population sizes are low (<10,000 individuals) or when prevalence levels are low (PfPR < 1%), MDA leads either to elimination or low-level persistence with dozens of infected individuals present (Figure 1). The simple reason is that populations starting with several hundred *Plasmodium* carriers will have these numbers reduced to dozens after the MDA, numbers that are low enough that they may lead to local elimination or a stuttering chain of transmission for a prolonged period. Rapid rebounds are rare in these scenarios. In larger populations (>40K individuals), epidemiological rebounds do occur in our model after an MDA (Figure 2A, 2D). In these cases, the basic reproduction ratio of *P. falciparum* will bring prevalence back to equilibrium levels which will be slightly lower than pre-MDA levels. As we assume that ACT adoption is increasing slowly over the twenty years of the simulation, equilibrium prevalence will be lower in 2042 than in 2022. However, ACT adoption is not rapid enough to interrupt malaria transmission permanently in the months following the last round of mass drug administration.

**Figure 2.**
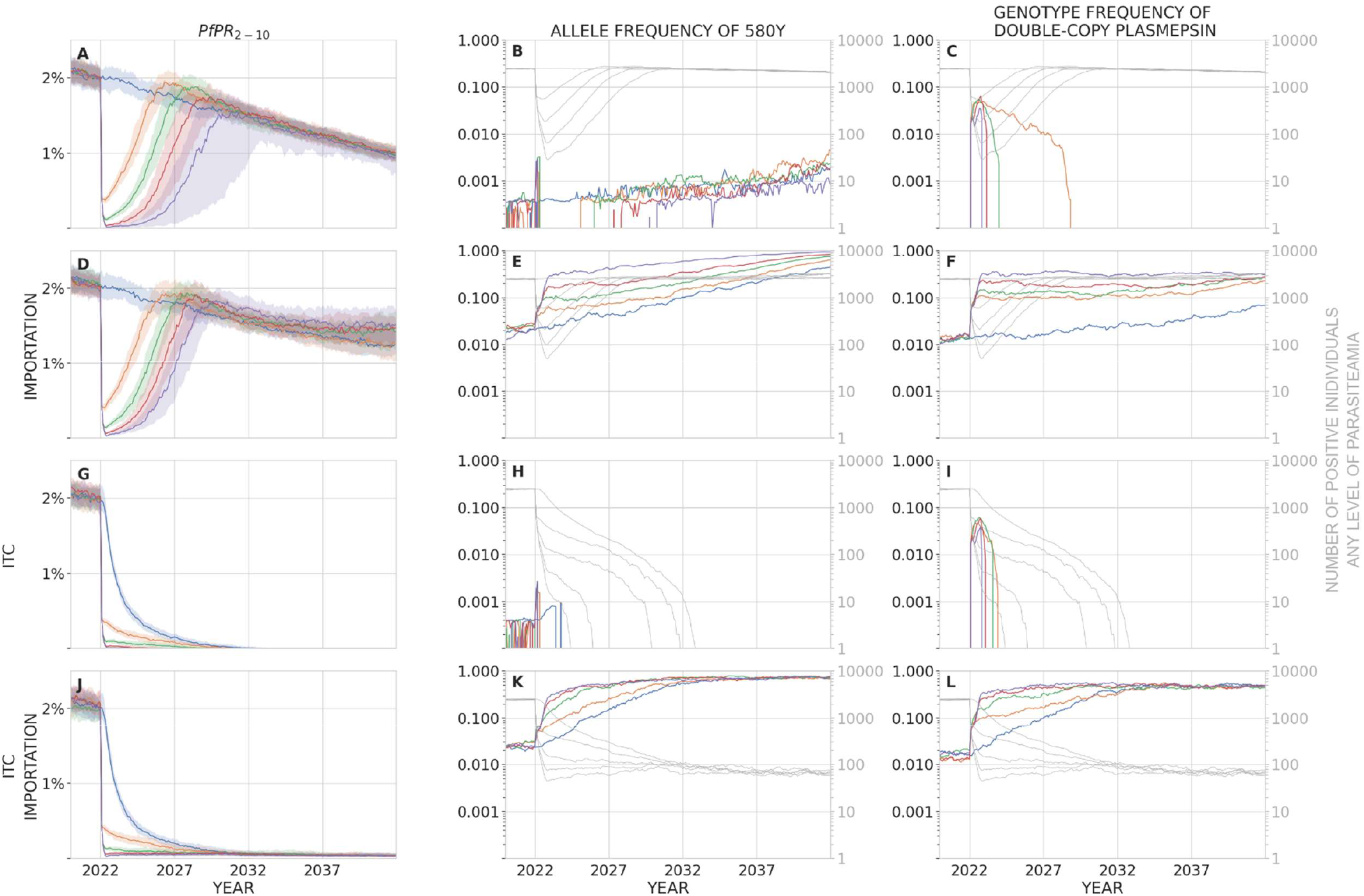
In a population of 40,000 individuals, panels show malaria prevalence (PfPR_2-10_, left), allele frequency of 580Y (middle), and the frequency of parasites with two or more copies of the plasmepsin-2,3 genes (right) over a period of 20 years after a mass drug administration has been carried out. In these scenarios, baseline PfPR_2-10_ = 2%. Median trajectories are shown from 100 simulations, and the shaded areas (left panels) show the interquartile range. Simulations are colored by the number of rounds of MDA carried out: blue (0), orange (1), green (2), red (3), purple (4). The top row (**ABC**) shows a scenario with no importation of drug-resistant genotypes and no improvement in treatment coverage (ITC) after the MDA is carried out. The second row (**DEF**) shows a scenario where new parasite importation occurs every 10 days; the imported parasite has a 50% probability of carrying the 580Y allele and a 50% probability of carrying multiple copies of the plasmepsin-2,3 genes, independently. The third row (**GHI**) shows a scenario with no importation, but where treatment coverage is increased post-MDA to 80% of the symptomatic patient population. The fourth row (**JKL**) shows a scenario with both importation and ITC. In the middle and right columns, the light gray lines show the absolute number of infected individuals (of any parasitaemia level) in the simulation and correspond to the right-hand gray tick marks on each panel. Note for example that in panels **K** and **L**, artemisinin-resistant and piperaquine-resistant genotype frequencies are very high, but in a population of only about 100 infected individuals, most of which are imports that occur during the course of the simulation; this region appears to have eliminated malaria but recently imported parasite-positive cases can still be found. MDA begins on January 1 2020, and the drop in prevalence in early 2020 can be seen in all plots in the left column. The bottleneck period lasts months to years, depending on the number of rounds of MDA carried out. Rapid selection of 580Y can be seen during the bottleneck period when importation is present (panels **E** and **K**). Double-copy plasmepsin variants are maintained in the population through linkage disequilibrium with 580Y genotypes (panels **F** and **L**, see Figure S4.1). The bottleneck period is risky when importation of 580Y alleles is expected; under these conditions, more rounds of MDA result in worse long-term drug-resistance outcomes.

In a baseline scenario with 40,000 individuals, no parasite importation, and PfPR_2-10_ = 2% (Figure 2A), one round of MDA (Jan 2022) brings median prevalence down to 0.37% after four months followed by a rebound in prevalence to 1.96% in 2026; four rounds of MDA result in a median prevalence of 0.013% (a total of 26 infected individuals at its lowest point) with the median simulation rebounding to 1.5% prevalence in 2030. When four rounds of MDA were carried out, 33/100 simulations resulted in malaria elimination; elimination was also observed for three rounds of MDA (8/100), but not for one or two rounds. Note that there is no re-importation of malaria in this baseline scenario, and there is little to no pre-existing artemisinin resistance (no 580Y alleles) at the start of the simulation. During treatment, parasites can mutate from one genotype to another as long as the new genotype provides a fitness advantage in the presence of drug treatment (e.g. C580 to 580Y mutations are allowed during AL and DHA-PPQ treatment); however, both real-world and modeled *de novo* mutation rates are low enough that new mutants are rarely observed in a population of thousands of infected individuals pre-MDA or hundreds of infected individuals post-MDA. For this reason, artemisinin resistance takes a long time to emerge in a population of 40,000 individuals, and 580Y allele frequencies in this scenario remain below 0.01 from 2022 to 2042 irrespective of the number of MDA rounds applied. The size and duration of the post-MDA bottleneck has no long-term effect on genotype distributions (Kruskal-Wallis *p* > 0.53, Figure 2B) as there are nearly no drugresistant alleles present during this period, and the occurrence of *de novo* mutations during this low-prevalence period is unlikely.

*P. falciparum* evolutionary dynamics following mass drug administration are qualitatively different when re-importation of drug-resistant alleles is considered (Figure 2D-2F), as would occur in a scenario where geographical areas receiving MDA are surrounded by untreated areas with drug resistance. When importation is occurring, the post-MDA bottleneck in the parasite population allows for more efficient selection of drug-resistant alleles: the smaller the bottleneck the faster the initial increase in gene frequency when a resistant allele is introduced. As expected, applying more rounds of MDA results in a smaller bottleneck and faster drug-resistance evolution. Figure 2E shows that each additional round of MDA has a detrimental effect on future artemisinin resistance when re-importation is common (KW *p* < 10^−12^). The long-term driver of this effect is the post-MDA continued use of ACT as first-line therapy, which maintains selection pressure on 580Y throughout the entire post-MDA period. With no MDA, 580Y alleles reach a median frequency of 0.038 in 2026; if MDA is implemented, median 580Y frequencies in 2026 are 0.06 (one round), 0.12 (two rounds), 0.18 (three rounds), and 0.32 (four rounds).

At higher malaria prevalence and higher population size, the bottlenecking effect is weaker. In a population of 300,000 individuals with PfPR_2-10_ = 5% (Figure 3), the parasite bottlenecks are larger (i.e. more parasite-positive individuals post-MDA) and of shorter duration, and elimination is not possible with ∼80% MDA coverage (Figures S1 and S2 show the dynamics when only prevalence or only population size are changed, respectively). With no re-importation (Figure 3A-3C), MDA has little effect on long-term artemisinin resistance evolution. In 2042, median 580Y frequencies are between 0.04 and 0.08 depending on the number of MDA rounds applied (KW *p* < 0.04; KW *p* > 0.12 when excluding 580Y frequencies after four rounds of MDA). The effects of mutation in this scenario are small but not completely negligible since the population of infected individuals is large enough for 580Y mutations to have appeared at low frequencies (median 0.0037; IQR: 0.0015-0.0074) prior to the initiation of MDA in 2022. There is some genetic variation for the MDA to act on, which is why a small spike in 580Y frequencies is observed in 2022. However, in the majority of scenarios, these mutants are rare and it is possible for them to not survive the bottlenecking period despite being selected for. When re-importation is allowed (Figures 3D-3F), we observe the same effects as in Figures 2D-2F, namely that more rounds of MDA result in higher artemisinin-resistance frequencies in the long run (KW *p* < 10^−5^). Piperaquine resistance is maintained in the population via positive linkage disequilibrium with artemisinin resistance (Figures S4).

**Figure 3.**
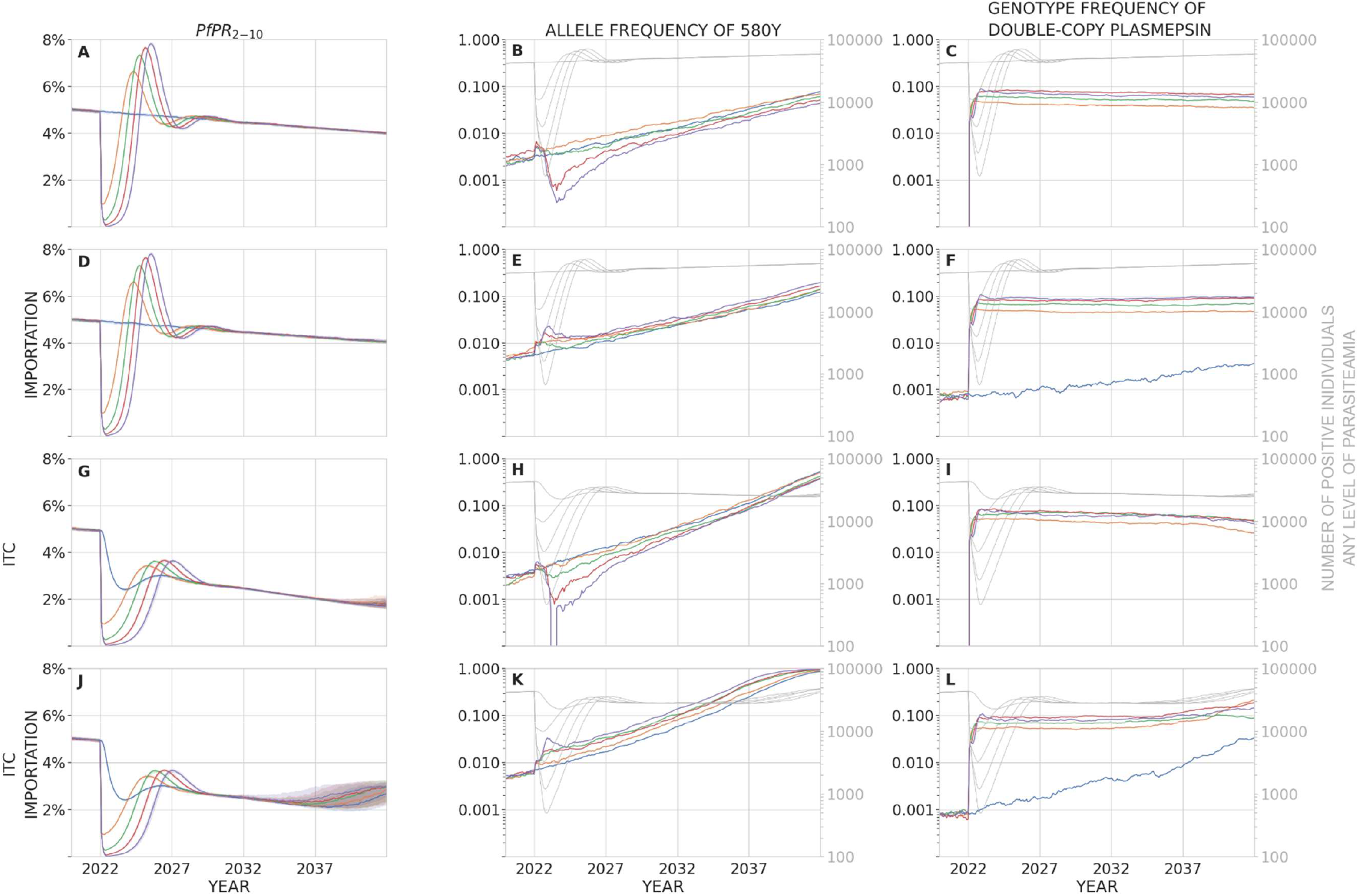
In a population of 300,000 individuals, panels show malaria prevalence (PfPR_2-10_, left), allele frequency of 580Y (middle), and the frequency of parasites with two or more copies of the plasmepsin-2,3 genes (right) over a period of 20 years after a mass drug administration has been carried out. In these scenarios, baseline PfPR_2-10_ = 5%. Median trajectories are shown from 100 simulations, and the shaded areas (left panels) show the interquartile range. Simulations are colored by the number of rounds of MDA carried out: blue (0), orange (1), green (2), red (3), purple (4). The top row (**ABC**) shows a scenario with no importation of drug-resistant genotypes and no improvement in treatment coverage (ITC) after the MDA is carried out. The second row (**DEF**) shows a scenario where new parasite importation occurs every 10 days; the imported parasite has a 50% probability of carrying the 580Y allele and a 50% probability of carrying multiple copies of the plasmepsin-2,3 genes, independently. The third row (**GHI**) shows a scenario with no importation, but where treatment coverage is increased post-MDA to 80% of the symptomatic patient population. The fourth row (**JKL**) shows a scenario with both importation and ITC. In the middle and right columns, the light gray lines show the absolute number of infected individuals (of any parasitaemia level) in the simulation and correspond to the right-hand gray tick marks on each panel. MDA begins on January 1 2020, and the drop in prevalence in early 2020 can be seen in all plots in the left column. The bottleneck period lasts months to years, depending on the number of rounds of MDA carried out. Selection of 580Y can be seen during the bottleneck period when importation is present (panels **E** and **K**), although this bottleneck effect is weaker in a population of 300,000 individuals (this figure) than a population of 40,000 individuals (Figure **2E** and **2K**). Double-copy plasmepsin variants are maintained in the population through linkage disequilibrium with 580Y genotypes (panels **C, F, I, L**, see Figure S4.2). The bottleneck period is risky when importation of 580Y alleles is expected; under these conditions, more rounds of MDA result in worse long-term drug-resistance outcomes.

The major MDA-related risk to safeguard against is exacerbation of drug resistance, wherein MDA generates higher drug-resistance frequencies than if no MDA had taken place. In our modeling analysis, the greatest risk of this ‘perverse effect’ occurs when all three of the following elements are present: (1) a small genetic bottleneck post-MDA without eliminating the parasites, (2) importation or pre-existence of drug-resistant parasites, and (3) persistent selection for drug-resistant alleles post-MDA. When no bottleneck is present, there are no rapid increases of gene frequency caused by small population sizes (compare blue and purple lines in Figures 2E and 3E). Without re-importation, there are no drug-resistant alleles to be selected for during the bottleneck and mutation is not rapid enough to generate them *de novo* when absolute population sizes are small (compare panels in Figures 2B/2E, 3B/3E). When there is no selection for artemisinin resistance or piperaquine resistance after the MDA, these alleles recede from the population (see Figures S3) due to an assumed fitness cost for resistant genotypes. Therefore, if one of these elements is missing, MDA should not exacerbate the population’s drug-resistance levels as compared to a policy of no MDA; in these cases MDA should have a neutral effect on drug-resistant alleles, and high-coverage MDA can be pursued without concern that drug-resistance will undermine the intended public health goals. Note that an MDA implementation followed by a small bottleneck and importation risks (elements 1 and 2 together) will still generate high levels of drug resistance, but this will not lead to treatment failures in patients if first-line therapy is changed after the MDA.

The dangers of drug-resistance importation during MDA can be seen more clearly when we look at a classic ‘waiting time’ measure in population-level drug resistance analyses; this is the time until a critical and potentially dangerous level of drug resistance has been reached in the population. Choosing 25% artemisinin resistance as our milestone, Figure 4 shows that 580Y alleles increase in frequency and reach 25% allele frequency earlier when importation is more frequent. This effect is more pronounced at smaller population size (40,000 individuals) and more rounds of MDA, as it is in these smaller bottlenecks that imported drug-resistance alleles have the largest advantage by starting their fixation process at a higher frequency.

**Figure 4.**
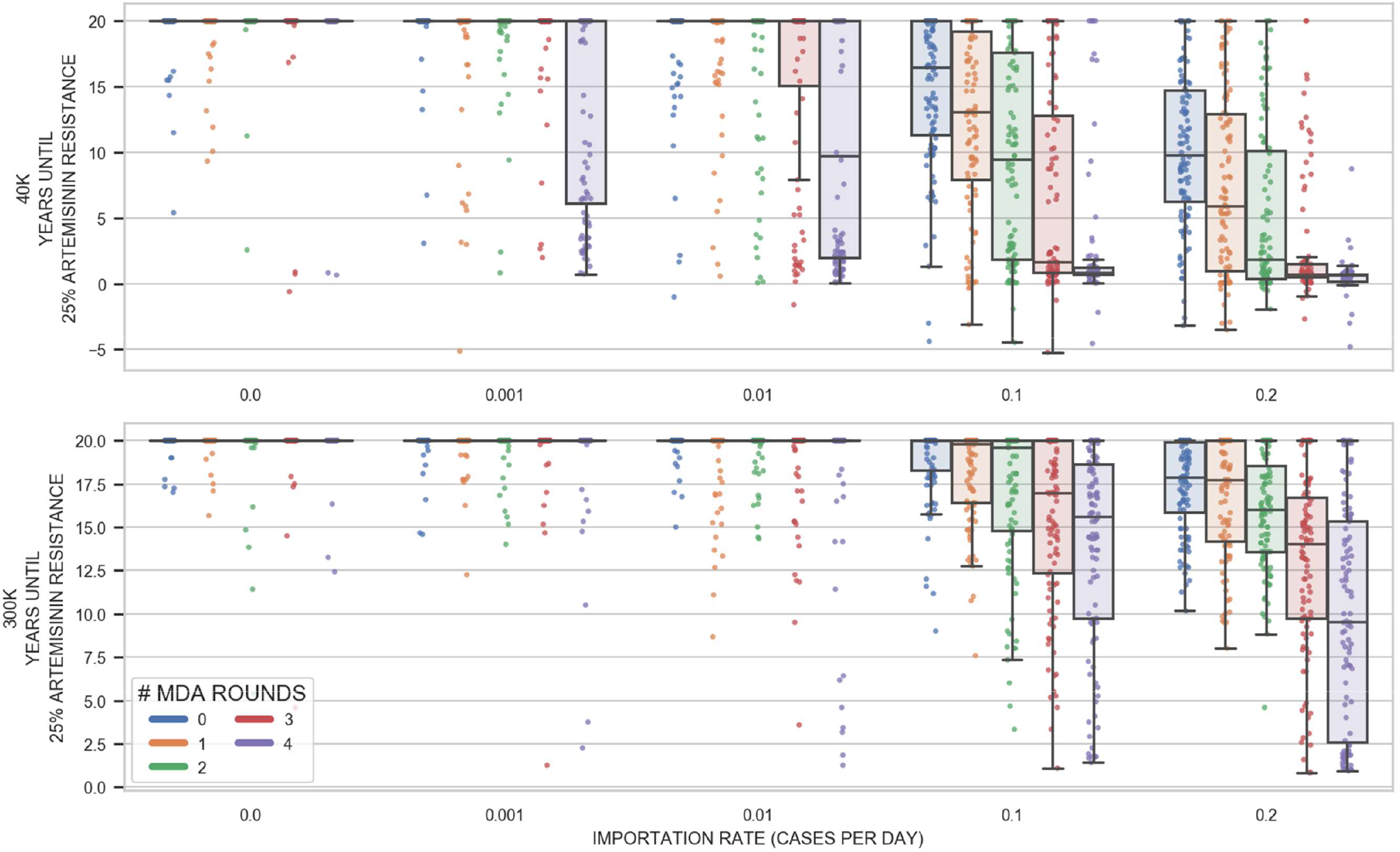
Effect of importation on artemisinin-resistance evolution in an MDA scenario with PfPR_2-10_=2%. The box+scatter plots show how long it takes for the allele frequency of 580Y to reach 0.25 in the simulation. Note that both the drug used in MDA (DHA-PPQ) and the first-line drug used to treat symptomatic *P. falciparum* malaria (AL) contain an artemisinin derivative, so selection on 580Y is occurring throughout the simulation. Five importation rates were considered (*x*-axis). The imported genotype has an equal 25% probability of being any of the four genotypes C580/580Y and single/double-copy plasmepsin. Alleles at other loci are randomly chosen. One hundred simulations were run for each importation rate, each of the two population sizes, and each possibility of zero to four rounds of MDA (5000 simulations in all). One dot in the graph corresponds to one simulation, and the boxes show the interquartile range for 100 simulations. With no or very low importation (0.001/day), there is little genetic variation in the population and the bottleneck period typically has no effect on selection. When importation of drug-resistance is regular, it is likely that drug-resistant genotypes will be present during the bottlenecking process; in these scenarios, more rounds or MDA lead to smaller bottlenecks and smaller bottlenecks result in more rapid selection. Treatment coverage is 55% for individuals over the age of 5, and 60% for children <5, and there is no improvement of treatment coverage in the post-MDA period. Overall, the bottleneck effect is stronger at smaller population sizes (40K v. 300K).

### 3.2 Improved treatment coverage (ITC) after the MDA

After an MDA program has completed all rounds, an opportunity presents itself to extinguish remaining chains of malaria transmission and achieve regional elimination. In September 2018, a WHO-convened evidence review group concluded that “maintaining reductions in transmission after the last round of MDA requires additional interventions, including vector control, case management, and intensified surveillance and response” [25]. These recommendations were reviewed by the Malaria Policy Advisory Committee (MPAC) in April 2019 emphasizing “that MDA must be thought of as a package together with other interventions” with a goal of reducing “transmission to the point that intensive case- and focus-based activities can be initiated” [33]. Field studies have confirmed that improved treatment coverage both pre- and post-MDA is likely to increase the chances of elimination and reduce the probability of an epidemiological rebound [3,20].

In a scenario with no re-importation of malaria, our model showed that improved treatment coverage (ITC) post-MDA can lead to malaria elimination if prevalence is low enough (Figure 2 GHI) but not if the starting prevalence is high (Figure 3 GHI). When re-importation of malaria cases is expected, a post-MDA ITC policy can still lead to elimination. Elimination in these settings is defined as a scenario where malaria may be imported but will not generate enough secondary cases to initiate an outbreak or establish endemically; imported cases in this scenario are eventually cured and/or treated as are the small number of secondary cases they generate. Figure 5 shows prevalence reductions five years post-MDA in a setting with PfPR_2-10_ = 1% and frequent re-importation. In a population of 40,000 individuals, four rounds of MDA and no ITC leads to a majority of simulations (53%) having fewer than 100 infected individuals (practically speaking, an elimination outcome), but 94% and 96% of simulations reach the <100 infections point when antimalarial treatment coverage post-MDA reaches 65% and 70%, respectively. In a population of 300,000 individuals, the probability of reaching elimination improves from 7% (no ITC) to 75% (ITC at 65%) and 98% (ITC at 70%) when improving access and coverage of antimalarials after the MDA. As long as baseline prevalence is low enough, MDA+ITC can lead to elimination even under a scenario of constant re-importation of drug-resistant malaria.

**Figure 5.**
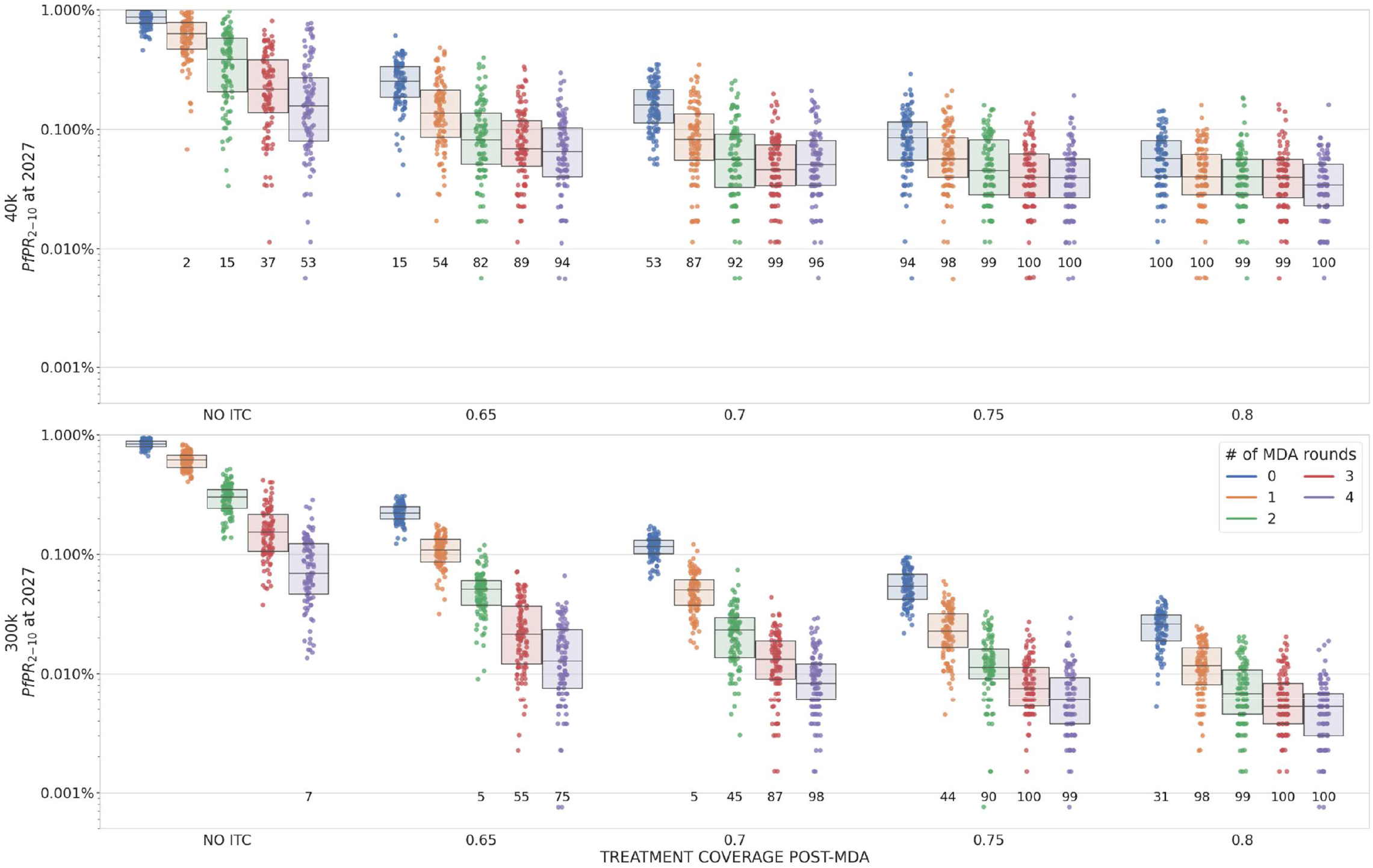
Box plots (*N*=100 simulations each) showing malaria prevalence five years after MDA; baseline PfPR_2-10_= 1%. The numbers printed below each plot indicate the number of simulations that ended in 2030 with fewer than 100 malaria cases present in the population (most of which are likely imports), i.e. a proxy for the number of simulations that achieved elimination. Importation of 580Y alleles is 0.1 cases/day. Numbers of rounds of MDA are shown by different colors, and four levels of improved treatment access are considered on the *x*-axis (65% access post-MDA to 80% access post-MDA). Improved treatment coverage post-MDA increases the chances of local elimination.

When baseline prevalence is high (PfPR_2-10_ ≥ 5%), malaria infections will rebound post MDA, even when ITC is made available (Figure 3 G-L). When re-importation is likely, this rebound will carry with it a high frequency of drug resistance (higher than if no MDA took place), and this is the countervailing evolutionary-epidemiological effect that must be avoided when considering MDA in higher prevalence settings. Under these circumstances, MDA would lead to higher resistance levels in the short-term and long-term, unless MDA participation and/or ITC levels could be pushed to higher levels than are modeled in the present analysis. This underscores the importance of knowing prevalence and reducing prevalence before implementing MDA.

## 4 Discussion

This work was commissioned by the WHO Evidence Review Group on Mass Drug Administration for Malaria (Sep 2018 [25]) and is intended to provide guidance for MDA scenarios in Africa – in particular, in locations where there may be importation risks of artemisinin-resistant genotypes from Rwanda [34] – and higher-prevalence SE Asia scenarios where artemisinin resistance is common and widespread [35]. Target population sizes of 40,000 and 300,000 individuals were evaluated as this was stated as the likely range of larger MDA programs supported by WHO [25]. In these settings, we find that strong population bottlenecks are associated with substantial uncertainty with respect to future patterns of antimalarial drug resistance, and we provide guidance on how to classify epidemiological scenarios when evaluating MDA-related drug-resistance risks. We discuss below some of the factors that are associated with the best possible outcome of an MDA, local malaria elimination, as well as the worst possible outcome, a rebound in malaria prevalence with higher drug-resistance levels than those expected under no MDA.

A combination of migration, drift, and selection – when all three are present simultaneously – can sometimes lead to a paradoxical epidemiological outcome where an MDA worsens long-term population-level health outcomes. In an epidemiological scenario with (1) a small bottleneck caused by the MDA, (2) likely re-importation of drug-resistant parasites, and (3) persistent selection pressure by the recommended first-line antimalarial post-MDA, long-term dynamics post-MDA will lead to an epidemiological rebound and higher frequencies of drug-resistance alleles when compared to a scenario with no MDA. Note that these detrimental outcomes typically appear several years after the MDA, allowing for a course correction if the risk has been identified. In addition, the re-importation requirement means that one would need to identify two highly connected populations with substantial drug resistance and perform MDA in only one of them to generate a worse-than-status-quo drug-resistance outcome. This serves as an obvious reminder to scrutinize regional epidemiological patterns before initiating MDA activities [3,20], and to focus MDA on higher-resistance areas first if prevalence levels there are sufficiently low.

In many African malaria settings conducive to an MDA-based prevalence reduction, importation of artemisinin-resistance alleles is unlikely and there would be little risk that an MDA leaves the population worse off than before. However, understanding the three requirements above is of special importance to many local malaria management questions in Southeast Asia as some areas may meet all three conditions. In a Southeast Asian setting with high local prevalence and a high chance of re-importation of artemisinin resistance, mass drug administration may exacerbate drug resistance instead of making it better. This risk assessment helps us underline the obvious conclusion that in a group of connected populations all of which have high drug-resistance levels, an MDA should be conducted for the entire region or not at all. A partial MDA implementation in this scenario would cause re-imported drug resistance to nullify the near-term beneficial effects of MDA. Fortunately, the two SE Asian MDA projects to date that have recorded *Pfkelch13* allele frequencies to assess artemisinin-resistance risk did not detect an increase in resistance post-MDA [3,11]. We propose that these three criteria be evaluated in future considerations of MDA in large populations. Criterion 1 will likely be met by any successful MDA program. The focus would thus be on questions of drug-resistance importation and the ability to substitute the first-line therapy once the MDA is complete.

Identification of these three conditions and their relationship to drug-resistance risk clarifies that the primary evolutionary force driving drug resistance is the first-line therapy used for routine case management post-MDA – and not the MDA drug itself – as the post-MDA bottleneck period is much longer than the MDA. This presents a new policy option of substituting out the first-line therapy after an MDA has completed, in order to remove selection pressure on any recently generated drug resistance. This policy option will need to be evaluated using specific geographic and treatment scenarios with a simulation approach similar to the one taken here. The principle that diverse drug environments should slow the evolutionary process should apply. The more treatment heterogeneity we can introduce into the parasites’ environment during and post-MDA, the more difficult it will be for newly emergent drug-resistant genotypes to spread successfully [36–39].

The most direct approach to avoiding drug-resistance risks that materialize during and after an MDA bottleneck is to drive parasite numbers to zero while the number of parasite-carrying individuals is still low. In our analysis, the key factor in reaching elimination is the treatment coverage post-MDA. If the infrastructure and knowledge put in place for the MDA – newly trained staff, larger catchment area for a local malaria post or community health workers, stable procurement of antimalarials, awareness among febrile individuals on how to access effective antimalarials – is able to be maintained in a way that improves treatment coverage in the population, elimination can be achieved in low-prevalence settings. The cost-effectiveness of post-MDA ITC needs to be assessed. As twelve months of a large-scale population health program are unlikely to cost twice as much as six months of the same program, an important question in the health economics of malaria management will be determining how long to run an ITC campaign for after an MDA has completed.

One current limitation of the modeling analyses presented here is the generalizability of these scenarios to other malaria-endemic settings. Many important epidemiological characteristics such as seasonal changes in mosquito populations, geographic extent of the MDA program [40], variation in household biting rates, and participation and compliance in each MDA round [14] will have important effects on the success of MDA programs. While it is true that African and Southeast Asian regions will have markedly different risks corresponding to different resistance profiles of circulating *P. falciparum* populations, partner-drug resistance will be present in both settings and will need to be accounted for as a potential cause of current and future treatment failures. As elimination plans move forward regionally in SE Asia, as well as elimination-ready regions in Africa and South America, MDA scenarios like the ones outlined here will need to be tailored to the exact populations and epidemiological characteristics of regions whose prevalence has been reduced to the point where MDA programs can realistically be used to accelerate towards elimination.

In the months immediately following a mass-drug administration program, prevalence management strategies yield more programmatic certainty than resistance management strategies. Low parasite numbers immediately post-MDA present an opportune period to drive prevalence to zero, but they do not provide good predictability on near-term evolutionary outcomes related to drug resistance. As more malariaendemic regions are brought to low-prevalence status, MDA will become more common as a policy option. Success will depend on assessment and management of re-importation risk and ability of health systems to commit resources to ensure high participation, long follow-up, and continual improvements in malaria control post MDA.

## Supporting information

Supplementary Materials 2 - Drug-by-Genotype Table

## Acknowledgements

This work was commissioned for WHO Evidence Review Group (ERG) on Mass Drug Administration for Malaria (Sep 11-13, 2018, Geneva). The work was funded by the Malaria Modeling Consortium (www.malariamodelingconsortium.org) via the University of Washington’s MMC grant from the Bill and Melinda Gates Foundation (OPP159934). Model simulations were done at Penn State’s Institute for Computational Data Sciences (ICDS) Advance Cyberinfrastructure (ACI) computing cluster. DMP is supported by funds from the Bill and Melinda Gates Foundation (OPP1211806) and the National Institutes of Allergy and Infectious Disease (FAIN: U19AI089672; Subaward number: 6123-1247-00-D). Thanks to Lorenz von Seidlein and François Nosten for valuable discussions.

## Author Contributions

TDN and MFB designed the analysis. TDN developed the model and ran the simulations. TN-AT reviewed literature and assembled the drug-by-genotype efficacy table. DMP provided grounding in field data and helped with parameterization of individual participation in MDA. NJW helped with literature review, field context, and likely future MDA application scenarios. MFB wrote the first draft of the paper.

