## Supplementary Materials 1 - Supp Methods and Figures for "Antimalarial mass drug administration in large populations and the evolution of drug resistance"

### Supplementary Appendix 1 to "Antimalarial mass drug administration in large populations and the evolution of drug resistance" by Nguyen, Tran, Parker, et al

#### 1. Updates to model since Nguyen et al (2015) [1]

Most model details are in the supplement (open-access) to the original paper which can be easily downloaded here: https://www.thelancet.com/cms/10.1016/S2214-109X(15)00162-X/attachment/dc5c729e-02cc-4d4e-9d61-35ad6894299c/mmc1.pdf

Some changes and updates are described below. Model behaviors that have not changed since 2015 are reviewed briefly.

##### 1.1. New Locus Structure

To allow for partial resistance and multiple different levels of drug resistance due to different mutations at different loci, a new locus representation was introduced into the model. This locus-based model includes positions N86Y and Y184F in the *pfmdr1* gene on chromosome 5. Copy number variation (CNV) is allowed for *pfmdr1*, but the alleles on the second copy are restricted to be the same as the alleles on the first copy. Eight configurations (haplotype) are possible for chromosome 5, and they can be coded or summarized in the following shorthand: `NY--, YY--, NF--, YF--, NYNY, YYYY, NFNF, YFYF`. The dashed lines indicate an absence of a second copy of *pfmdr1*. The model does not distinguish between two copies or more than two copies.

On chromosome 7, the K76T alleles are included for the *pfcrt* gene. On chromosome 13, the C580Y alleles are included for the *pfkelch13* gene. And on chromosome 14, copy number variation is allowed for the *plasmepsin-2,3* genes.

This results in 64 total genotypes, where the wild-type can be coded as `KNY--Y1` or `KNY1Y1` if we choose to use the "1" to indicate one copy of *pfmdr1*. Details of how these 64 genotypes map to phenotypes (resistant phenotypes associated with particular treatment efficacies) is in Supplementary Appendix 2.

Details of the resistant alleles and relationships between genotypes and phenotypes (drug efficacies) were describes in the "Sumplement-TT.pdf" file.

The chromosomal positions of the loci are important because our model includes recombination (sexual reproduction) at the point of transmission to the mosquito, when a mosquito bites a multi-clonally infected host. The "force of infection" of an individual infected with multiple clones is the sum of all forces of infection for all possible recombinant offspring that could be produced with the host's parasites. The potential recombinant offspring are listed using a standard recombination table, and the entries in the table (the probabilities of forming a particular recombinant) account for the current parasite densities of clones in an individual's blood. No interrupted mosquito feeding is modeled, so recombination only occurs if there is multi-clonality in the populations.

#### 1.2. Parasite Density

Symptomatic individuals start with an asexual parasite density between 2000 parasites per microliter (/µl) and 200,000/µl. For symptomatic hosts, this number is drawn uniformly on a $\log_{10}$-scale between 3.301 and 5.301. For new bites that progress to asymptomatic or sub-clinical infection, $\log_{10}$-parasite density is drawn from a log-normal distribution with mean 3.0 and standard deviation 0.5. Asymptomatic density will get progressively lower as the immune system works to reduce the overall parasite density. Asymptomatic infections last between 60 and 281 days, depending on the host's initial immune status. Within host, drug-resistant clones experience a cost of resistance in the absence of the drug. For a single-resistant clone (i.e. a genotype with one resistant allele or one CNV), the daily cost of resistance is $c_R = 0.0005$, which results in a 17% genotype frequency drop over one year compared to a wild-type parasite population. The daily cost of resistance for a parasite clone with *n* genetic mechanisms (non-wild-type) conferring resistance is $c_R = 1 - (1 - 0.0005)^n$.

#### 1.3. Public and Private Market

In the new model, individuals can seek and receive malaria treatment either in the public sector or in the private market. In the baseline scenario set-up, the fraction of public market use starts at 5% in 2008 and increases linearly to 80% in 2042, which means the fraction of private market use will wane from 95% to 20% from 2008 to 2042. This means that in 2022, approximately 36% of malaria treatment seeking occurs in the public sector (recommended first-line therapy used, RFT) while 64% occurs in private markets or pharmacies (off-policy purchases, OPP). In the baseline set-up presented here, artemether-lumefantrine (AL) is the RFT in the public sector, while private-market drug purchases include sulfadoxine-pyrimethamine (30%), amodiaquine monotherarpy (30%), chloroquine (30%), and AL (10%).

#### 1.4. Importation

The new model allows for importation of new genotypes into the population. In the baseline setting for importation scenarios, a new parasite importation occurs every 10 days, on average, according to a Poisson process; the imported parasite has a 50% probability of carrying the 580Y allele and a 50% probability of carrying multiple copies of the *plasmepsin-2,3* genes, independently. All 64 genotypes have an equal 1/64 probability of being imported into the population.

##### 1.5. Mass Drug Administration

The new model contains a mechanism for mass drug administration (MDA). Not all individuals in a population will participate in each round of the MDA. Participation rates vary depending on the communication strategy prior to the MDA, the size of the population, the number of study staff, and the occupations and livelihoods of the residents living in the area. Low participation rates are in the 60% range while a high participation rate would be around 85%. Children and elderly individuals are likely to have higher participation rates. Individuals that miss one round of MDA are likely to miss another round, due to their occupation, travel patterns, hesitancy, or some other reason. Therefore, the missingness across rounds is not random across individuals in the population. To model this feature, each individual in the model is given a probability of MDA participation. These probabilities are drawn from a beta distribution with mean 0.85 (children and elderly) or 0.75 (working age adults, and older children, ages 10-40). The standard distribution of the beta-distribution is 0.3 to capture the variation in participation and non-participation across individuals in the population.

Four rounds of MDA take place in the scenarios modeled here. In the simulation, the duration of each round of MDAs is 14 days and rounds are 5 weeks apart. Individual will be assigned uniformly a day in 14 days of MDA period to receive an antimalarial.

##### 1.6. Improved Treatment Coverage (ITC)

During the MDA campaign, the treatment coverage for symptomatic individuals increases from 50% to ~80% for a few months. We evaluated scenarios where the infrastructure from MDA scale-up was reused to maintain the high treatment coverage, with the highest levels modeled at 80% drug coverage for routine malaria infections. After the MDA, treatment coverage improved linearly over a six-month period before reaching its maximum of 80%.

##### 1.7 Pharmacokinetics and Pharmacodynamics of OZ439 and Ferroquine

These drugs were used in an experimental set of simulations (Figures S3.1 to S3.4) where we examined the benefits of substituting out artemether-lumefantrine (AL) as first-line therapy after the MDA and replacing it with a combination therapy of OZ439 and ferroquine (FQ). Half-life of OZ439 was assumed to be 2.5 days [2] and half-life of FQ was taken as 10 days [3]. In McCarthy et al (J Antimicrob Chemother, 2016 [4]), OZ439 plasma concentration half-life was reported as 260h (geometric mean) for a 500mg dose, while in the 200mg dose group the half-life was 51h. Hence, we chose 2.5 days (60h) as the plausible value for OZ439 half-life.

The 48-hour parasite reduction ratio ($PRR_{48}$) of OZ439 was estimated to be 10176 (95% CI 5757 - 17986) for a 500mg dose and 165 (95% CI 124 -222) for 200 mg doses [4]. Then, the $PRR_{24}$ estimate will be 100.86 (95% CI 75.87 – 134.11) for a 500mg dose; i.e. the fractional parasite killing per day will be in the range 0.9700 to 0.9925. In Phyo et al [2], the killing rate per hour was estimated to be 0.13 and 0.17, for 400mg and 800 mg doses, respectively; which yields a range of parasite killing per day between 0.9646 and 0.9886. In the simulation, we set the maximum fraction of parasites that can be killed per day ($p_{\max}$) to be 0.9990 for a host with a very high (40% above average) OZ439 drug concentration. For an average drug concentration ($C_0$=1.0) with $EC_{50}$ = 0.75, the factor $\left(\frac{C_t^n}{C_t^n+EC_{50}^n}\right)$ results in a killing rate of 0.9467, just below the indicated range above.

For FQ, the $PRR_{48}$ was reported as 163 (95 % CI 141–188) which is equivalent to $PRR_{24}$ of 12.77 (95% CI 11.87 – 13.71) [3]. Daily fractional parasite killing is estimated to be between 0.9157 and 0.9271. In the simulation we choose the maximum fraction of parasites that can be killed per day to be $p_{\max}$ = 0.99 for the host which results in > 0.9 killing rates for high drug concentrations.

The efficacy of the combination OZ439-FQ was calculated by running five simulations on 10,000 patients each. Each patient had a per-μl parasitaemia drawn from a log-uniform distribution between 2000 and 200,000. 28-day efficacies ranged from 95.5% to 95.9% in the five trials.

Summary of the PK/PD parameters used in the simulation for OZ439 and FQ:

| | OZ439 | FQ |
|---|---|---|
| Half-life ($t_{0.5}$) | 2.5 days (Phyo et al., 2016) | 10 days (McCarthy et.al., 2016) |
| The maximum fraction of parasites that can be killed per day ($p_{max}$) | 0.999 | 0.99 |
| Slope of the concentration-effect curve ($n$) | 10 | 5 |
| The drug concentration at which the parasite killing is 50% of $p_{\max}$ ($EC_{50}$) | 0.75 | 0.75 |

### 2. Supplementary Figures

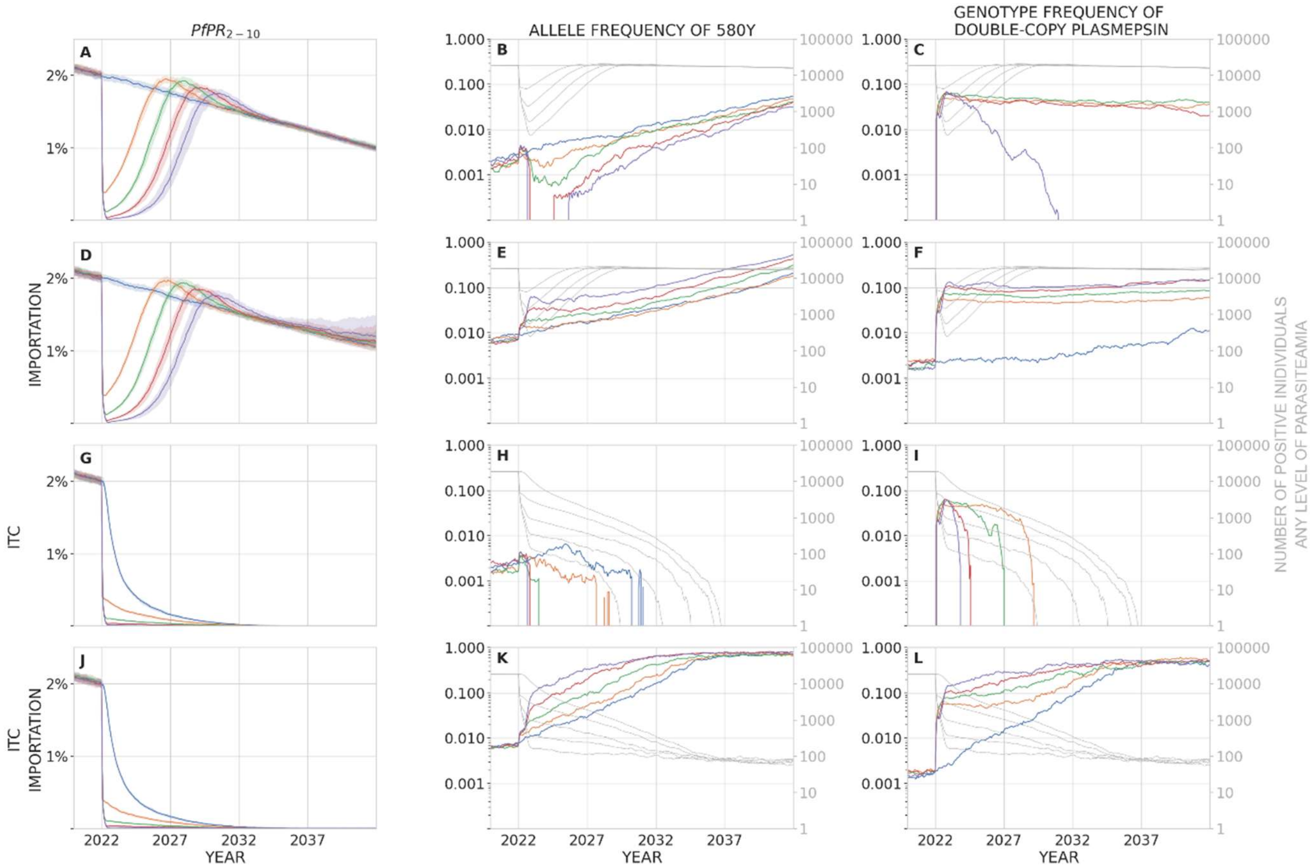


**Figure S1.** In a population of 300,000 individuals, panels show malaria prevalence ($PfPR_{2\text{-}10}$, left), allele frequency of 580Y (middle), and the frequency of parasites with two or more copies of the plasmepsin-2,3 genes (right) over a period of 20 years after MDA has been carried out. In this scenario, baseline $PfPR_{2\text{-}10}$ = 2%. Median trajectories are shown from 100 simulations, and the shaded areas (left column only) show the interquartile range. Simulations are colored by the number of rounds of MDA carried out: blue (0), orange (1), green (2), red (3), purple (4). The top row shows a scenario with no importation of drug-resistant genotypes and no improvement in treatment coverage (ITC) after the MDA is carried out. The second row (panels DEF) shows a scenario where a new parasite importation occurs every 10 days, on average; the imported parasite has a 50% probability of carrying the 580Y allele and a 50% probability of carrying multiple copies of the plasmepsin-2,3 genes, independently. The third row (panels GHI) shows a scenario with no importation, but where treatment coverage is increased post-MDA to 80% of the symptomatic patient population. The fourth row (panels JKL) shows a scenario with both importation and ITC. In the middle and right columns, the light gray lines show the absolute number of infected individuals in the simulation (of any parasitaemia level) and correspond to the right-hand gray tick marks on each panel. Note for example that in panels K and L, artemisinin-resistant and piperaquine-resistant genotype frequencies are very high, but in a population of only about 100 infected individuals, most of which are imports that occur during the course of the simulation; this region appears to have eliminated malaria but imported parasite-positive cases can still be found in the region. MDA begins on January 1 2022, and the drop in prevalence in early 2022 can be seen in all plots in the left column. The bottleneck period lasts months to years, depending on the number of rounds of MDA carried out. Rapid selection of 580Y can be seen during the bottleneck period when importation is present (panels E and K). Double-copy plasmepsin variants are maintained in the population through linkage disequilibrium with 580Y genotypes (panels F and L, see Figure S4.1). The bottleneck period is risky when importation of 580Y alleles is expected; under these conditions, more rounds of MDA result in worse long-term drug-resistance outcomes.

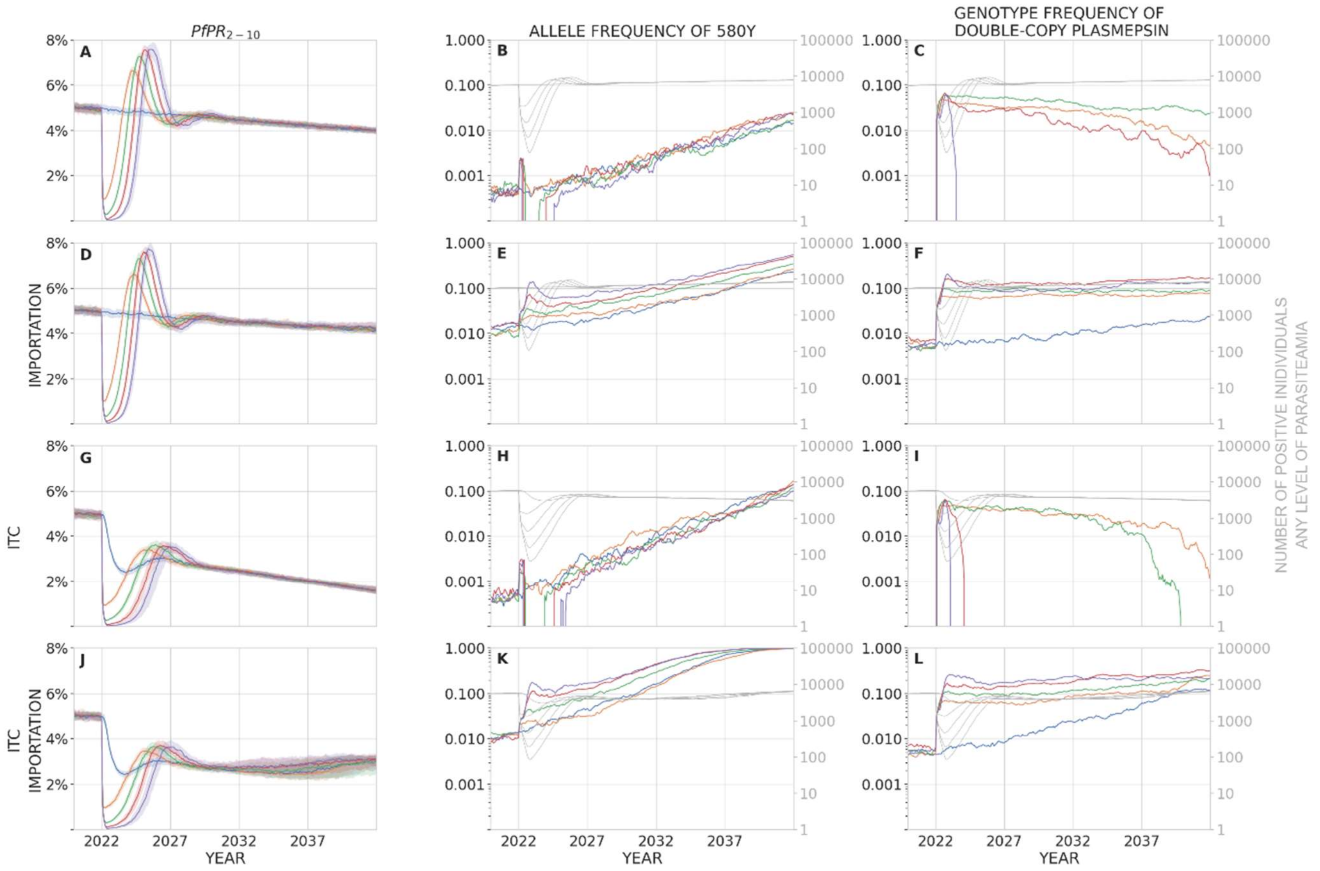


**Figure S2.** In a population of 40,000 individuals, panels show malaria prevalence ($PfPR_{2\text{-}10}$, left), allele frequency of 580Y (middle), and the frequency of parasites with two or more copies of the plasmepsin-2,3 genes (right) over a period of 20 years after a mass drug administration has been carried out. In this scenario, baseline $PfPR_{2\text{-}10}$ = 5%. Median trajectories are shown from 100 simulations, and the shaded areas (left column only) show the interquartile range. Simulations are colored by the number of rounds of MDA carried out: blue (0), orange (1), green (2), red (3), purple (4). The top row shows a scenario with no importation of drug-resistant genotypes and no improvement in treatment coverage (ITC) after the MDA is carried out. The second row (panels DEF) shows a scenario where a new parasite importation occurs every 10 days, on average; the imported parasite has a 50% probability of carrying the 580Y allele and a 50% probability of carrying multiple copies of the plasmepsin-2,3 genes, independently. The third row (panels GHI) shows a scenario with no importation, but where treatment coverage is increased post-MDA to 80% of the symptomatic patient population. The fourth row (panels JKL) shows a scenario with both importation and ITC. In the middle and right columns, the light gray lines show the absolute number of infected individuals in the simulation (of any parasitaemia level) and correspond to the right-hand gray tick marks on each panel. MDA begins on January 1 2022, and the drop in prevalence in early 2022 can be seen in all plots in the left column. The bottleneck period lasts months to years, depending on the number of rounds of MDA carried out. Selection of 580Y can be seen during the bottleneck period when importation is present (panels E and K), although this bottleneck effect is weaker in a higher prevalence scenario (this figure) than in a low prevalence scenario (Figure 2E and 2K). Double-copy plasmepsin variants are maintained in the population through linkage disequilibrium with 580Y genotypes (panels C, F, L, see Figure S4.2). The bottleneck period is risky when importation of 580Y alleles is expected; under these conditions, more rounds of MDA result in worse long-term drug-resistance outcomes.

**Figure S3.1 – S3.4.** In a population of 40,000 individuals, artemisinin pressure was removed by deploying an OZ439-Ferroquine combination to replace AL as first-line therapy in 2022. Panels show malaria prevalence ($PfPR_{2-10}$, left), allele frequency of 580Y (middle), and the frequency of parasites with two or more copies of the plasmepsin-2,3 genes (right) over a period of 20 years after a mass drug administration has been carried out. In this scenario, baseline $PfPR_{2-10}$ = 2%. Median trajectories are shown from 100 simulations, and the shaded areas show the interquartile range. Simulations are colored by the number of rounds of MDA carried out: blue (0), orange (1), green (2), red (3), purple (4). In these scenarios, a new parasite importation occurs every 10 days, on average; the imported parasite has a 50% probability of carrying the 580Y allele and a 50% probability of carrying multiple copies of the plasmepsin-2,3 genes, independently. In each figure, the top row shows a scenario with no improvement in treatment coverage (ITC) after the MDA is carried out and the bottom row shows a scenario with ITC. In the middle and right columns, the light gray lines show the absolute number of infected individuals in the simulation (of any parasitaemia level) and correspond to the right-hand gray tick marks on each panel. For all sub-scenarios **S3.1** to **S3.4**, the fixation of 580Y and plasmempsin-2 copy in the E and F panel was due to the 25% chance of importing a new double-copy plasmepsin and 580Y genotype while the number of positive individuals dropped below 100 in the population.

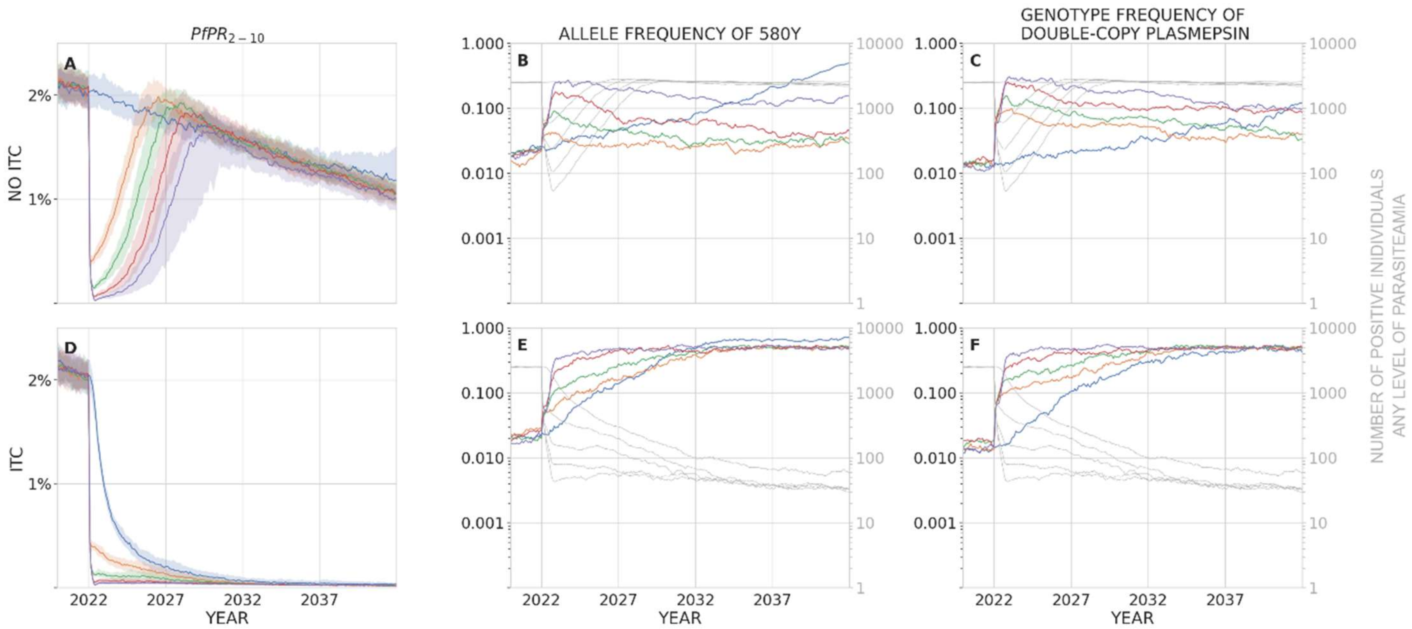


**Figure S3.1.** AL in public sector was replaced by OZ439+FQ after the final round of MDA. For the no-MDA scenario (blue line), AL continues to be used through 2042. AL is still used in the private sector.

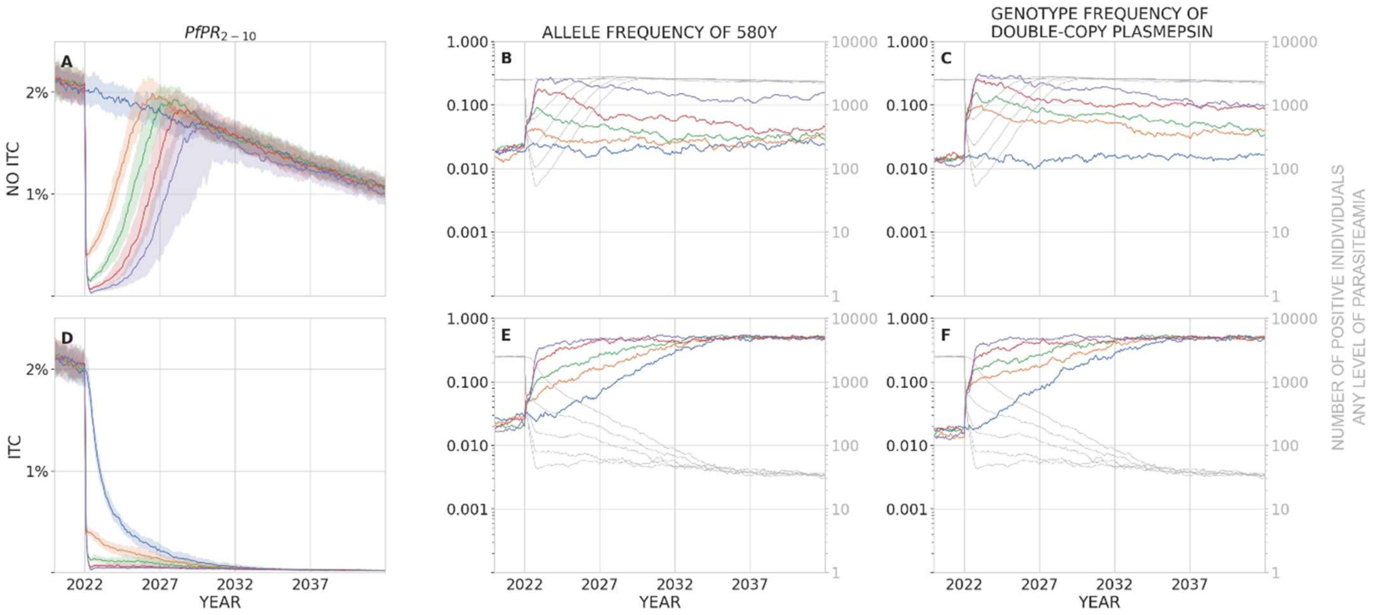


**Figure S3.2.** As Figure S3.1 but AL is replaced by OZ439-FQ under the no MDA scenario (blue line) as well. AL is still used in the private sector.

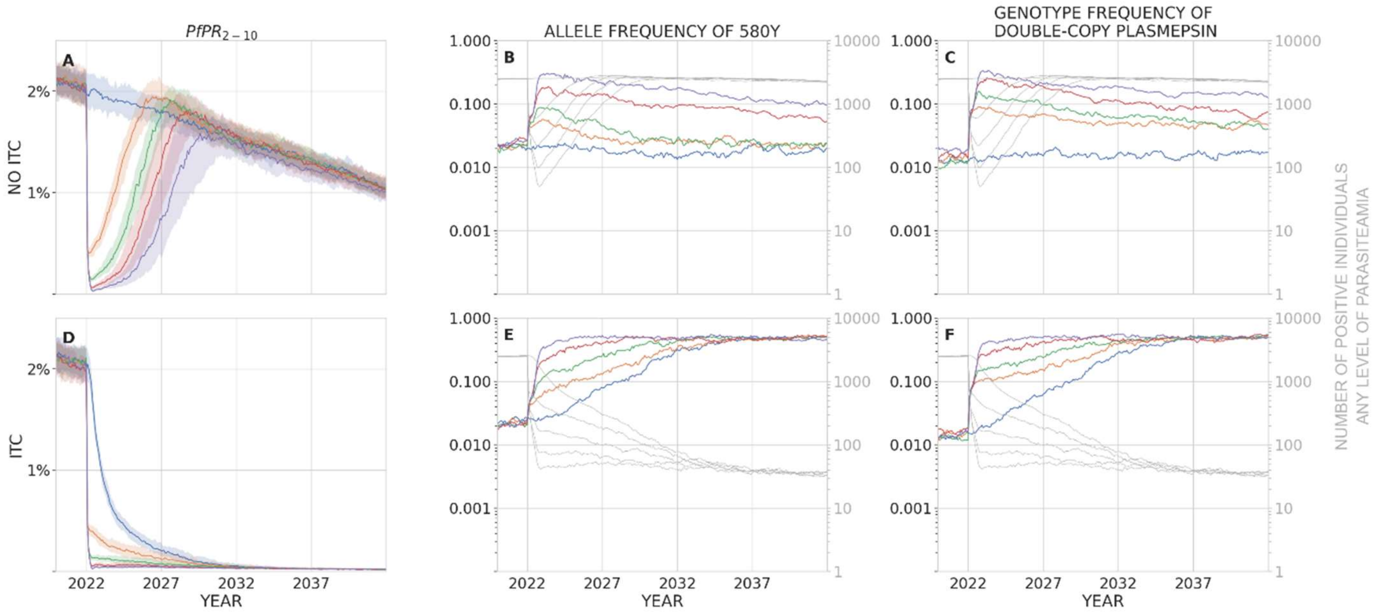


**Figure S3.3.** Unlike in S3.1 and S3.2, AL is replaced by OZ439-FQ prior to the MDA. AL is still used in the private sector.

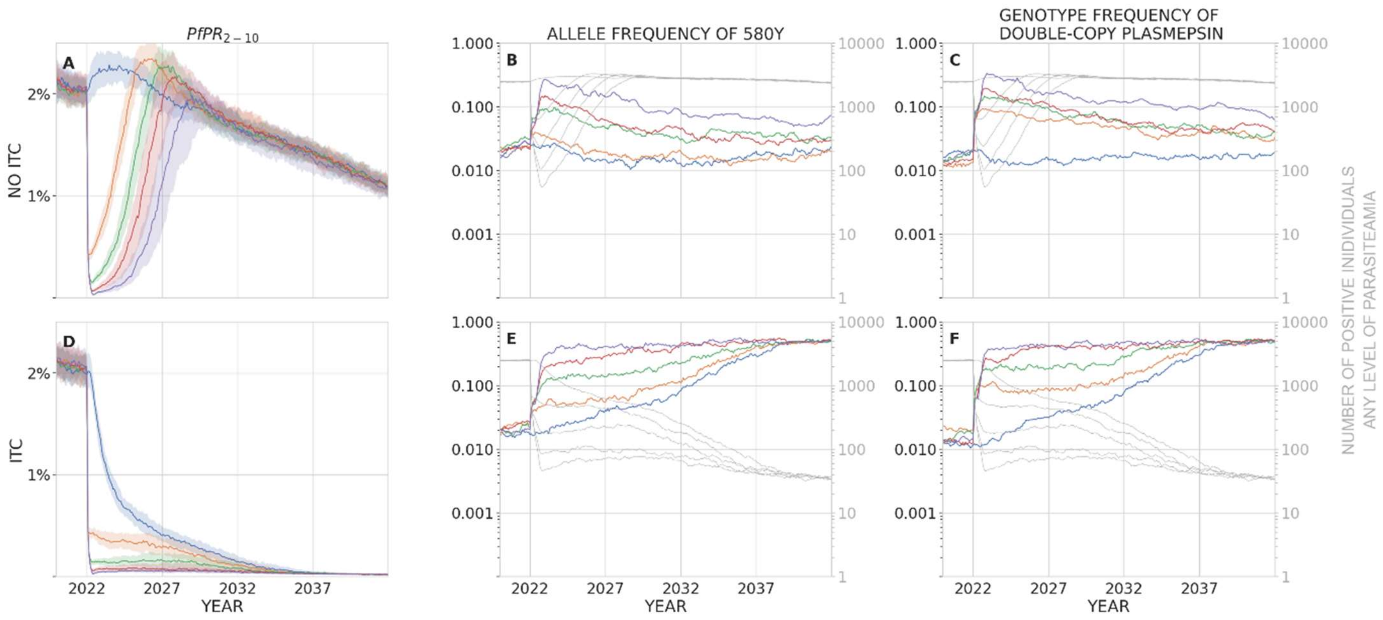


**Figure S3.4.** AL is replaced by OZ439-FQ prior to the MDA, and AL is also removed from the private sector. 580Y frequencies in Panel B are lower when comparing to Figure S3.3.

Note that 580Y frequencies recede in all four scenarios. They recede most slowly in **Fig S3.1** and most quickly in **Fig S3.4** where all artemisinin pressure is removed. 580Y frequencies are still highest under four rounds of MDA, but they are not associated with treatment failure because the first-line therapy post-MDA was changed to OZ439-FQ.

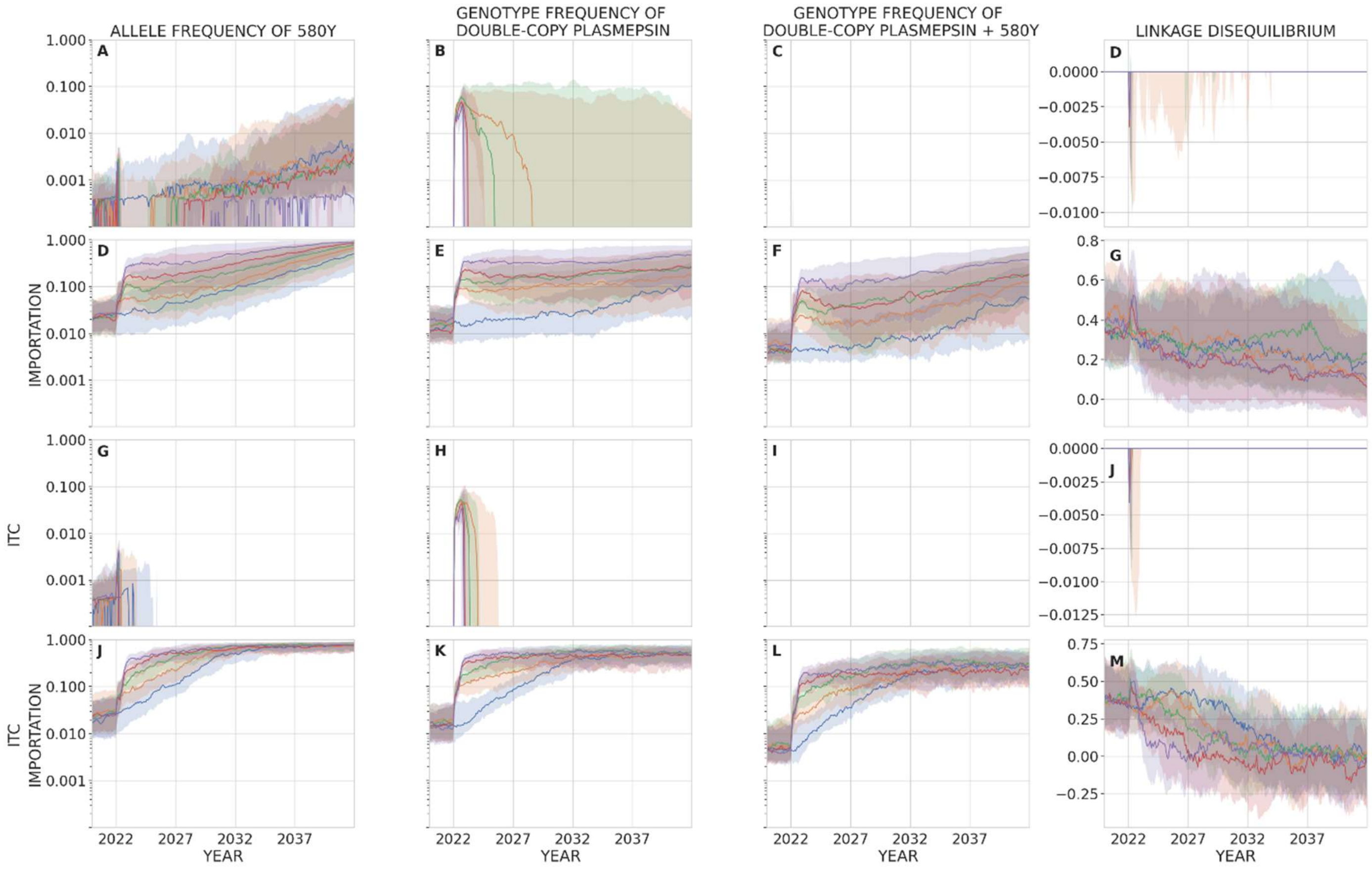


**Figure S4.1.** Scenario settings are population size of 300,000 and 2% PfPR. Panels shows allele frequency of 580Y (first column), the frequency of parasites with two or more copies of the plasmepsin-2,3 genes (second column), the frequency of parasites with both 580Y and two or more copies of the plasmepsin-2,3 genes (third column), and linkage disequilibrium between 580Y and two or more copies of the plasmepsin-2,3 genes (fourth column) over a period of 20 years after a mass drug administration has been carried out. In the second row LD is positive between the two loci (due to selection), and this helps maintain piperaquine resistance in the population even when PPQ is not used as part of the first-line therapy.

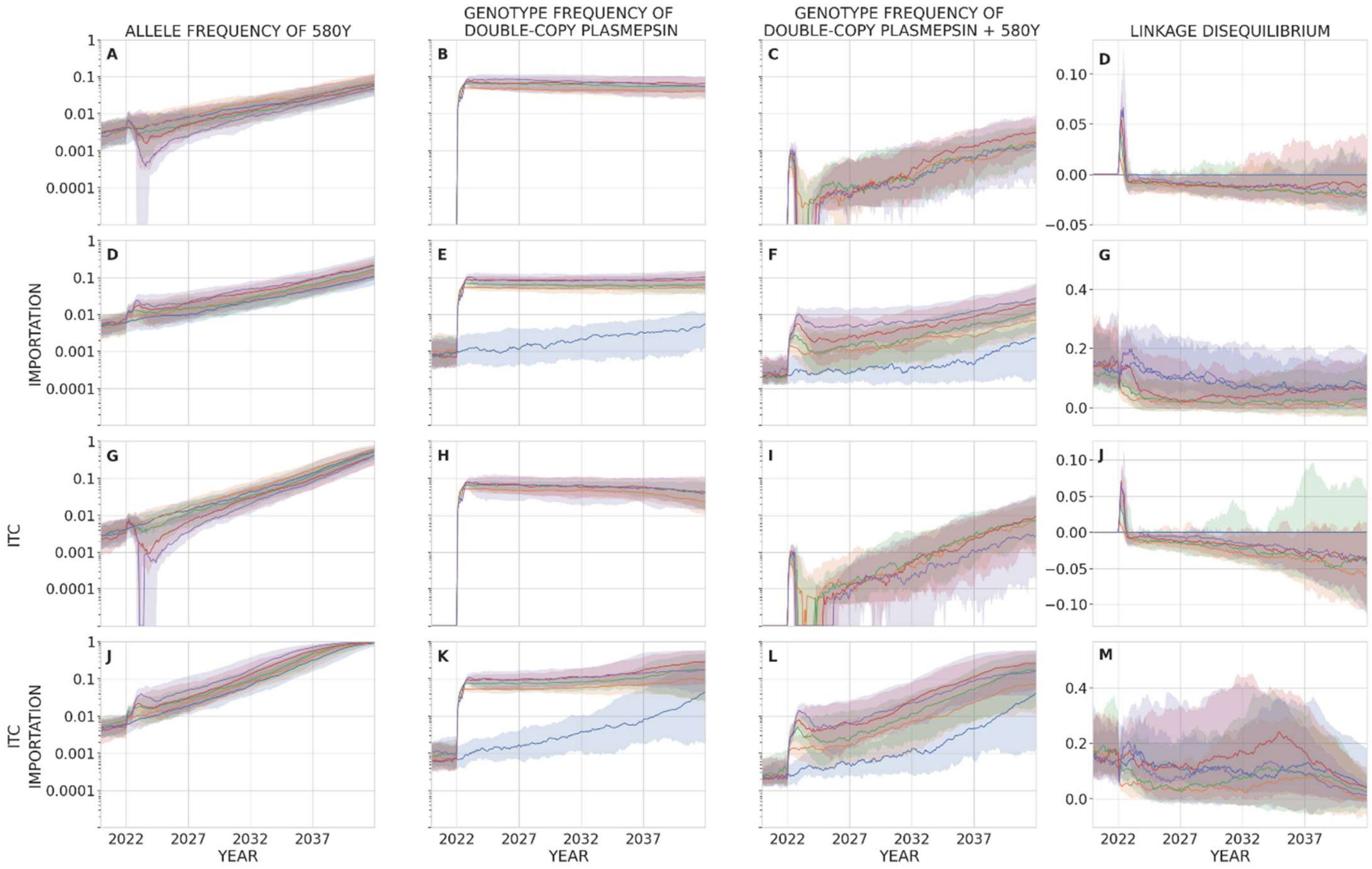


**Figure S4.2.** Scenario settings are population size of 300,000 and 5% PFPR (as in Figure 3 of main text). Panels shows allele frequency of 580Y (first column), the frequency of parasites with two or more copies of the plasmepsin-2,3 genes (second column), the frequency of parasites with both 580Y and two or more copies of the plasmepsin-2,3 genes (third column), and linkage disequilibrium between 580Y and two or more copies of the plasmepsin-2,3 genes (fourth column) over a period of 20 years after a mass drug administration has been carried out. In the second and fourth rows LD is positive between the two loci (due to selection), and this helps maintain piperaquine resistance in the population even when PPQ is not used as part of the first-line therapy.
