## Supplementary Materials 2 - Drug-by-Genotype Table for "Antimalarial mass drug administration in large populations and the evolution of drug resistance"

**Supplementary Appendix 2 to "Antimalarial mass drug administration in large populations and the evolution of drug resistance" by Nguyen, Tran, Parker, et al**

### Construction of Drug-by-Genotype Table

#### Table of Contents


***Table of Contents*** **1**

***1 Background and Objectives*** **1**

***2 Estimating EC50 values and resulting efficacy of oral artesunate monotherapy*** **5**

**2.1 Inferring an EC50 for AS and AM on C580 genotypes** **5**

**2.2 Inferring an EC50 for AS and AM on 580Y genotypes** **9**

2.2.1 PPQ monotherapy and DHAPPQ on *kelch13*-C580 and single *plasmepsin-2,3* genotypes 9

2.2.2 EC50 for AS on 580Y genotypes and PPQ on multiple-copy *plasmepsin-2,3* genotype 11

***3 Estimating EC50 for lumefantrine monotherapy using trial data on artemether-lumefantrine (AL) combination therapy*** **15**

**3.1 AL therapeutic efficacy studies** **15**

**3.2 Validation of Results with Older Trials of Lumefantrine Monotherapy** **18**

***4 MQ, ASMQ, CQ*** **23**

**4.1 MQ, ASMQ** **23**

**4.2 CQ** **27**

***5 AQ, ASAQ*** **29**

***6 References*** **35**


### 1 Background and Objectives

In this document, we approximate the efficacy of six antimalarial monotherapies and four combination therapies on 64 *P. falciparum* genotypes. The purpose of this exercise is to generate a drug-by-genotype (DxG) table – essentially a genotype-by-environment interaction matrix – that will allow mathematical models of *P. falciparum* drug resistance evolution to model any antimalarial therapy use on any genotype, as any combination of therapy and genotype should be possible in a simulation or model run.

We consider the following ten antimalarials: artesunate (AS), lumefantrine (LM), amodiaquine (AQ), piperaquine (PPQ), mefloquine (MQ), chloroquine (CQ), artemether-lumefantrine (AL), dihydroartemisinin-piperaquine (DHA-PPQ), artesunate-amodiaquine (ASAQ), and artesunate-mefloquine (ASMQ).

We consider four loci and two copy-number variations. The four loci are K76T on *pfcrt* gene (chromosome 7), N86Y on *pfmdr1* gene (chromosome 5), Y184F *pfmdr1* gene, and C580Y on *kelch13* gene (chromosome 13). We consider parasites that have multiple copies of the *pfmdr1* gene, but we only allow for multiple copies that have the same alleles at the 86 and 184 loci. Additionally, we include copy-number variation of the *plasmepsin-2,3* genes (chromosome 14). These loci were chosen as they have known drug-resistance effects on many commonly used antimalarials. Some of the allowable genotypes have been simplified in this analysis to keep the problem tractable. A total of 64 genotypes are allowed for in this analysis as copy-number variants (CNV) are simply allowed to be 'single' or 'multiple'.

Our goal is to approximate the entries in a table where each cell represents the (PCR-corrected) day-28 efficacy of a particular antimalarial monotherapy or combination therapy on an uncomplicated case of *Plasmodium falciparum* with a particular genotype. There will be 640 entries to approximate. We will start with known genotype-specific efficacies from the literature. The main challenge here is that there the vast majority of trials do not report parasite genotypes, and indeed "therapeutic efficacy studies are not powered to test for the association between parasite genotypes and treatment outcome" (Ljolje et al. 2018). Some historical assumptions will be made about what genotypes were likely to be circulating on a particular continent in a particular decade. Some approximate inference will be carried out (using a stochastic model) when, for example, the efficacy of a combination therapy and one of the underlying monotherapies is known.

After the first version of the DxG table was completed, we asked two clinicians with extensive experience in malaria therapeutic efficacy studies (TESs) and drug-resistance genotypes to review the table. We sat down with each clinician independently and did not show them the other clinician's opinions. Without revealing the entries in the table, we asked for expected treatment efficacies for certain drug-genotype combinations where our literature search showed the most variation, and we asked for the best estimate of these particular efficacies in a modern context with appropriate dosing, quality control, and follow-up measurements. Resulting from these conversations, some of the efficacies were adjusted manually, and these instances are indicated in this document.

| Antimalarial | Allele that it selects | References |
|---|---|---|
| AS | *kelch13*-580Y | (Ariey et al. 2013; Ashley et al. 2014) |
| LM | *pfcrt*-K76 | (Humphreys et al. 2007; Sisowath et al. 2009; Mwai et al. 2009; Raman et al. 2011; Duah et al. 2013; Thomsen et al. 2013; Hemming-Schroeder et al. 2018) |
| | *pfmdr1*-N86 | (Sisowath et al. 2005; Humphreys et al. 2007; Sisowath et al. 2009; C. T. Happi et al. 2009; Mwai et al. 2009; Raman et al. 2011; Duah et al. 2013; Thomsen et al. 2013; Hemming-Schroeder et al. 2018) |
| | *pfmdr1*-184F | (Humphreys et al. 2007; Sisowath et al. 2007; C. T. Happi et al. 2009; Malmberg et al. 2013; Thomsen et al. 2013; Hemming-Schroeder et al. 2018) |
| | multiple copies of *pfmdr1* | (Sidhu et al. 2006; R. N. Price et al. 2006; Mungthin et al. 2010; Duah et al. 2013; Venkatesan et al. 2014) |
| AQ | *pfcrt*-76T | (Humphreys et al. 2007; Holmgren et al. 2007; Tinto et al. 2008; Mandi et al. 2008) |
| | *pfmdr1*-86Y | (Humphreys et al. 2007; Holmgren et al. 2007; Nsobya et al. 2007; Tinto et al. 2008) |
| | *pfmdr1*-Y184 | (Humphreys et al. 2007; Holmgren et al. 2007) |
| PPQ | multiple copies of *plasmepsin-2,3* | (Amato et al. 2017; Witkowski et al. 2016) |
| MQ | multiple copies of *pfmdr1* | (Wilson et al. 1993; Ric N. Price et al. 2004; Sidhu et al. 2006) |
| CQ | *pfcrt*-76T | (Fidock et al. 2000; Djimdé et al. 2001; Dorsey et al. 2001) |
| | *pfmdr1*-86Y | (Djimdé et al. 2001) |

***Table S1*** *The following selection scheme is implemented in our model. In our allele-locus-allele notation, as usual, the left-hand allele is the wild-type (ancestral) allele and the right-hand allele is the mutant (derived) allele. However, note that the derived allele is not always the resistant allele.*

The table above shows the six drug compounds considered here as well as the alleles and CNVs that they select for. In the individual-based simulation used in the main paper – which has a daily time-step and in some cases will only approximate pharmacokinetics (PK) and pharmacodynamics (PD) – we assume a basic one-compartmental PK-model, i.e. instantaneous absorption to the initial drug concentration $C_0$ and then drug decay as described below

$$C_t = C_0 \cdot e^{\frac{-ln2}{T_{0.5}}t}$$

where

$t$ = elapsed time (in days) since the time of drug administration.

$C_t$ = drug concentration in the patient's plasma at day $t$.

$T_{0.5}$ = elimination half-life of the drug in day unit (Table S6 (Nguyen et al. 2015)).

Drug concentration changes with the simulation's daily time step. A person with average drug absorption ability would have initial drug concentration $C_0 = 1.0$. $C_0 < 1.0$ and $C_0 > 1.0$ indicate below-average and above-average drug absorption, respectively.

For PD-component, we assume drugs act independently of one another. The PD equations used in the simulation are as follows:

$$\text{fraction\_parasites\_removed\_per\_day} = p(C_t) = p_{max} \cdot \left(\frac{C_t^n}{C_t^n + EC_{50}^n}\right)$$

$$\text{parasite\_density\_at\_day\_t} = P_t = \left(1 - p(C_t)\right) \cdot P_{t-1}$$

where

$p_{max}$ = the maximum fraction of parasites that can be killed by a monotherapy in a single day (see Table S5 (Nguyen et al. 2015)).

$n$ = slope of the concentration-effect curve.

$EC_{50}$ = the drug concentration at which the parasite killing reaches 50%, i.e. $\frac{p_{max}}{2}$.

If a total of $m$ antimalarials are present in the patient's plasma simultaneously (e.g. in the case of combination therapies), each drug $i$ would have a separate set of values for $p_{max}$, $C_t$, $n$, $EC_{50}$ and the resulting parasite density in the patient's blood at the end of day $t$ would be:

$$P_t = P_{t-1} \cdot \prod_{i=1}^{m} \left(1 - p(C_{i,t})\right)$$

As we can see, varying any of the three PD parameters of an antimalarial would affect the parasite killing effect of its monotherapy as well as its combinations with other drugs. Additionally, we do not distinguish different derivatives of artemisinin, i.e. we use the same values of PK/PD parameters for artesunate, artemether, and dihydroartemisinin. For convenience, we use 'artesunate' as the representative of all artemisinin derivatives.

The drug-by-genotype table is constructed in order to help us calibrate $EC_{50}$ values of each antimalarial on each genotype. Other parameters such as slope $n$, and $p_{max}$ are kept unchanged, i.e. identical to those being used in (Nguyen et al. 2015).

On the following page, we show an empty drug-by-genotype (DxG) table with 640 blank entries. We will fill out the table section by section, starting with literature on artesunate and artemether monotherapy. In the first column of the table, we differentiate 64 genotypes by 7-character strings whose character 1, 2, 3, and 6 denote alleles at loci 76 on *pfcrt*, 86 on *pfmdr1*, 184 on *pfmdr1*, and 580 on *kelch13*. Character 4, and 5 in a genotype string show whether the genotype has single (labeled "--") or multiple copies of *pfmdr1* gene. In the case of multiple copies of *pfmdr1*, the pair of alleles represented at character 4 and 5 must match those at character 2 and 3. The last character in the genotype string tells us how many copies of *plasmepsin-2,3* gene are present in the genotype. When filling out the table step by step, we will use red to indicate efficacies which can be verified by previous therapeutic efficacy studies and orange to indicate inferred efficacies.

| Genotype | **AS** | **LM** | **AQ** | **PPQ** | **MQ** | **CQ** | **AL** | **ASAQ** | **DHAPPQ** | **ASMQ** |
|---|---|---|---|---|---|---|---|---|---|---|
| KNY--C1 | | | | | | | | | | |
| KNY--C2 | | | | | | | | | | |
| KNY--Y1 | | | | | | | | | | |
| KNY--Y2 | | | | | | | | | | |
| KYY--C1 | | | | | | | | | | |
| KYY--C2 | | | | | | | | | | |
| KYY--Y1 | | | | | | | | | | |
| KYY--Y2 | | | | | | | | | | |
| KNF--C1 | | | | | | | | | | |
| KNF--C2 | | | | | | | | | | |
| KNF--Y1 | | | | | | | | | | |
| KNF--Y2 | | | | | | | | | | |
| KYF--C1 | | | | | | | | | | |
| KYF--C2 | | | | | | | | | | |
| KYF--Y1 | | | | | | | | | | |
| KYF--Y2 | | | | | | | | | | |
| KNYNYC1 | | | | | | | | | | |
| KNYNYC2 | | | | | | | | | | |
| KNYNYY1 | | | | | | | | | | |
| KNYNYY2 | | | | | | | | | | |
| KYYYYC1 | | | | | | | | | | |
| KYYYYC2 | | | | | | | | | | |
| KYYYYY1 | | | | | | | | | | |
| KYYYYY2 | | | | | | | | | | |
| KNFNFC1 | | | | | | | | | | |
| KNFNFC2 | | | | | | | | | | |
| KNFNFY1 | | | | | | | | | | |
| KNFNFY2 | | | | | | | | | | |
| KYFYFC1 | | | | | | | | | | |
| KYFYFC2 | | | | | | | | | | |
| KYFYFY1 | | | | | | | | | | |
| KYFYFY2 | | | | | | | | | | |
| TNY--C1 | | | | | | | | | | |
| TNY--C2 | | | | | | | | | | |
| TNY--Y1 | | | | | | | | | | |
| TNY--Y2 | | | | | | | | | | |
| TYY--C1 | | | | | | | | | | |
| TYY--C2 | | | | | | | | | | |
| TYY--Y1 | | | | | | | | | | |
| TYY--Y2 | | | | | | | | | | |
| TNF--C1 | | | | | | | | | | |
| TNF--C2 | | | | | | | | | | |
| TNF--Y1 | | | | | | | | | | |
| TNF--Y2 | | | | | | | | | | |
| TYF--C1 | | | | | | | | | | |
| TYF--C2 | | | | | | | | | | |
| TYF--Y1 | | | | | | | | | | |
| TYF--Y2 | | | | | | | | | | |
| TNYNYC1 | | | | | | | | | | |
| TNYNYC2 | | | | | | | | | | |
| TNYNYY1 | | | | | | | | | | |
| TNYNYY2 | | | | | | | | | | |
| TYYYYC1 | | | | | | | | | | |
| TYYYYC2 | | | | | | | | | | |
| TYYYYY1 | | | | | | | | | | |
| TYYYYY2 | | | | | | | | | | |
| TNFNFC1 | | | | | | | | | | |
| TNFNFC2 | | | | | | | | | | |
| TNFNFY1 | | | | | | | | | | |
| TNFNFY2 | | | | | | | | | | |
| TYFYFC1 | | | | | | | | | | |
| TYFYFC2 | | | | | | | | | | |
| TYFYFY1 | | | | | | | | | | |
| TYFYFY2 | | | | | | | | | | |

### 2 Estimating EC50 values and resulting efficacy of oral artesunate monotherapy

#### 2.1 Inferring an EC50 for AS and AM on C580 genotypes

Among the ten therapies we include in the drug-by-genotype table, four of them are artemisinin combinations; hence, calibrating $EC_{50}$of artesunate should be prioritized. Several studies allow us to measure the efficacy of artesunate monotherapy on *P. falciparum* infections. These studies describe 3-day, 5-day, and 7-day courses of treatment. They are summarized below.

**Gabon 2001, children 4-15yo**. From January 2001 to April 2001, 50 P.*falciparum*-infected children aged 4-15 at Albert Schweitzer Hospital in Lambarene, Gabon were included in a study to evaluate the efficacy of 3-day artesunate monotherapy (Borrmann et al. 2003). Mean weight of the patients was 25.3 kg and the geometric mean of initial parasitaemia was 22000/µl (range: 2100-200000). Artesunate was given under supervision at 4 mg/kg dose once daily for 3 days and day-28 PCR-corrected cure rate was 36/50=72%.

**Thailand 1998-1999**. From April 1998 to March 1999, (Ittarat et al. 2003) treated 104 patients, aged 13-49, at Bangkok Hospital for Tropical Diseases with artesunate monotherapy at total dose of 600 mg over 3 days. The inclusion criteria did not restrict the range of initial parasitaemia, hence, parasite density at admission was reported to be as high as 776000/µl. The patients remained in the hospital for the whole study period. At the study's endpoint, there were 32 recrudescence cases and the 28-day efficacy of 3-day artesunate monotherapy was 72/104=69.2%.

**Hainan Island (China) 1982-1984.** In Hainan Island, China, from 1982 to 1984, (Li et al. 1984) recruited 80 *P.falciparum*-infected patients, aged 9-57, into a 4-arm trial to compare efficacy of mefloquine plus Fansidar (group A), mefloquine plus qinghaosu (group B), mefloquine plus Fansidar and qinghaosu (group C), and qinghaosu alone (group D); each arm had 20 patients. The initial parasitemia ranged from 1840 to 353157 parasite/µl, with mean of 57414 parasite/µl. All regimens were given as one single dose except for group D which was 3 doses over 3 days. The total dose of mefloquine was 750mg, and that of Fansidar was 75 mg pyrimerthamine plus 1500 mg sulfadoxine in all regimens. Total dose of qinghaosu was 1000 mg in group B, and C, and 2000 mg in group D. The patients remained in the hospital for the first 7 days of the study, then came back for follow-up on day 14, 21, and 28. Day-28 radical cure rates, after excluding vivax cases, were 100% for mefloquine-Fansidar, 100% for mefloquine-qinghaosu, 100% for mefloquine-Fansidar-qinghaosu, and 10/17=58.8% for qinghaosu 3-day monotherapy. The cure rates were calculated based on World Health Organization (WHO) 1973 grading scale for resistance (World Health Organization 1973), hence, they shall be consider as PCR-uncorrected.

**China 1994.** A series of studies in China from the 1990s and early 2000s during Novartis' early development of Coartem (artemether-lumefantrine) contain results on the efficacy of artemether monotherapy; these are summarized in FDA New Drug Application (NDA) 22-268 submitted in 2008 (Novartis Pharmaceuticals Corporation 2009). Study AB/MO2 conducted in 1994 in China reported 28-day parasitological cure rates (not PCR-corrected) in three trial arms: 3 days AL (N=51), 3 days artemether monotherapy (N=52), 3 days lumefantrine monotherapy (N=52). In all three arms, dosing occurred at 0h, 8h, 24h, 48h. The 155 patients were aged 13 to 57, and all cases were confirmed *P. falciparum*. Inclusion criterion for parasitaemia was parasite density between 1,000/µL and 100,000/µL, with 12 patients included in the trial despite having parasite densities >100,000/µL. Median baseline parasite densities in the three arms were between 19,000 and 27,000 parasites/µL. Reported efficacies were 100.0% for Coartem, 92.2% for lumefantrine, and 54.5% efficacy for 3-day artemether (per protocol, PCR-uncorrected). The summary review document of this NDA also mentioned a trial conducted by Chinese Academy of Military Medical Sciences (AMMS2 study) with three arms similar to those in study AB/MO2 above. Sixty patients aged 14-46 were recruited (twenty for each arm) and day-28 cure rates were 18/20=90% for the combination, 8/20=45% for artemether monotherapy, and 15/20=75% for lumefantrine monotherapy. Other inclusion criteria such as weight, initial parasitaemia as well as whether the efficacy was PCR-corrected were not mentioned in the document.

**Thailand, probably 1990-1991.** (Bunnag et al. 1991) showed that 5-day monotherapy of a 600mg artesunate dose given either once daily (n=25) or twice daily with half-doses (n=25) to *P.falciparum*-confirmed Thai adults (mean age 24.6-24.7 years old, mean weight 52.1-54.1 kg). For both groups, the first day's total dose was 200mg and subsequent doses were 100mg daily. Day-28 efficacies were 72% and 76%, respectively. Mean initial parasitaemia counts in both arms were 12129 and 16443 parasites/µl. The patients remained in the hospital during the course of treatment (28 days).

**Thailand 1991 or 1992.** (Karbwang et al. 1992) carried out a trial in Thailand in to compare efficacy of 5-day mefloquine monotherapy and 5-day oral artemether monotherapy. Total dose in each arm was 1250mg and 700mg, respectively. Patients in the study were 46 males aged 15-50, weighing 45-65 kg, with acute uncomplicated *P.falciparum*. 12 patients were assigned to the mefloquine arm, and 34 to artemether; they remained in the hospital for 42 and 28 days, respectively. Parasitaemia ranged from 3900 to 149260 parasites/µl; geometric mean of initial parasitaemia was 23438/µl for mefloquine arm and 13490/µl for artemether. Drugs were given at 0h, 6h, 30h, 54h,

78h, and 102h. The outcomes were assessed based on WHO criteria used at the time (World Health Organization 1973), and treatment success was defined as parasitological cure on day 28. Efficacy of 5-day artemether was 29/30 = 96.7%; day-42 efficacy for mefloquine arm was 7/11 = 64%.

**Thailand 1991.** (Looareesuwan et al. 1992) carried out a study evaluating the efficacies of artesunate (AS), mefloquine (MQ) and artesunate mefloquine (ASMQ) combinations in Thailand 1991. 127 patients were recruited and assigned to 3 different treatment arms: oral AS (100mg immediately, then 50mg every 12h for 5 days, total dose 600mg), oral MQ (750mg immediately, then 500mg at 6h; total dose 1250mg), and ASMQ (same oral 5-day AS regimen; two doses of oral MQ given on day 6 as in MQ arm). The baseline parasitaemia ranged from 172/µl to 184,400/µl, with geometric mean between 14,195/µl to 25,825/µl. The patients were 16-60 years old, weighed 45-60kg, and agreed to remain in hospital for 28 days. 28-day efficacies, measured as parasitological cure, were 88% for 5-day mono AS (n=40), 81% for MQ (n=37), and 100% for ASMQ (n=39).

**Thailand 1993.** A study in Bangkok (Karbwang et al. 1995) enrolled patients into two arms: artemether monotherapy (300mg on day 0, then 100mg on day 1-4) and sequential artemether (300mg at hour 0) followed by MQ (750mg at hour 24), in 109 Thai male patients with uncomplicated multi-drug resistant (MDR) falciparum (resistant to CQ and SP, with probable resistance to quinine and MQ). Ages 13-47. 53 patients enrolled in the AM monotherapy arm, and 56 in the AM-MQ arm. Baseline parasitaemia ranged from 360/µL to 403,340/µL, with mean parasitaemia 43,000 and 52,000 in the two arms. Efficacy of 5-day artemether was 88%. Efficacy of 1-day AM plus 1-day MQ was 94%. Only patients who completed 28-day for AM and 42-day for ASMQ hospital follow-up were included in the efficacy calculation. There is no molecular/genotyping detail as to the extent of drug-resistance of parasites in the enrolled patients, but clearly they could not have been very resistant to MQ as the AL-MQ arm had high efficacy. It is probably safe to assume that these are chloroquine-resistant parasites (76T in *pfcrt*), and the MDR label probably refers to CQ, Q, and SP.

**Vietnam 1994.** A 2-arm study (Giao et al. 2001) with 227 patients was conducted in southern Vietnam (Binh Thuan) in 1994. Patients with uncomplicated *P. falciparum* were given oral artemisinin monotherapy (0h, 8h, then once daily) for 5 days (arm 1) or 7 days (arm 2). Parasite density inclusion criterion was 1,000-100,000/µL; nevertheless, some patients with hyperparasitaemia >100,000/µL were enrolled. Primary outcome was 'radical cure' defined as parasite clearance by day 7 with no recrudescence (determined by microscopy) by day 28 (World Health Organization 1973). There is no information as to whether recrudescences were PCR-corrected, but prevalence in Binh Thuan province would have been high enough (>10%) in some communes that re-infections may have occurred (Hung et al, Bull WHO, 2002; Nam et al, TMIH, 2005). Ages 4 to 58 enrolled. Per-protocol artemisinin efficacies were 74.8% (5-day) and 77.7% (7-day) with no statistical difference between the two.

**Bangladesh 2008-2009.** (Starzengruber et al. 2012) conducted a trial in southeastern Bangladesh. Males and non-pregnant females aged 8 to 65 years with uncomplicated *P. falciparum* were recruited; the inclusion criterion for parasitaemia was parasite density between 1000 and 100,000/µl. The primary endpoints were defined as adequate clinical and parasitological response (ACPR, defined in (World Health Organization 2003) on day 28 and on day 42 (PCR-corrected). There were 3 treatment arms in the trial, artesunate 2mg/kg once daily for 7 days, artesunate 4mg/kg once daily for 7 days, and quinine (10mg/kg) combined with doxycycline (4mg/kg) for 7 days. A total of 106 recruited patients reached the primary endpoint. The geometric mean of the initial parasite density ranged from 7,910 to 9,137 parasites/µl. Efficacies did not differ between 28-day and 42-day endpoints. Efficacies at day 28 were 97.8% for 7-day AS (2mg/kg), 100% for 7-day AS (4mg/kg), and 100% for Q+DOXY.

To approximate the optimal EC50 value for artesunate, we simulated cohorts of 10,000 patients, each cohort with a different EC50 value and a different dosing regimen (3-day, 5-day, 7-day); see Figure S1 below. For each 10,000-patient simulated trial, we computed the squared error between the simulated efficacy and the observed efficacy. Using the 13 arms in the 10 trials listed here, this gives a sum-squared error over 13 trials, which is shown on the y-axis in Figure S1. The optimal EC50 was 0.71 which corresponded to a 3-day AS efficacy of 72.2%. After consultation with two independent reviewers (clinicians with field experience in malaria) and comparison with the 3-day data only, we manually adjusted this efficacy so that it fell below 70%. The final chosen EC50 was 0.75 which corresponded to a 3-day AS efficacy of 68.9%. A balance was needed to keep the efficacy of ACTs high and the efficacy of 5-day AS and 7-day AS high.

Hence, we fill in 32 entries in the table – all genotypes that are C580 – with the value 0.689.

(A similar approach was taken for other compounds in subsequent sections).

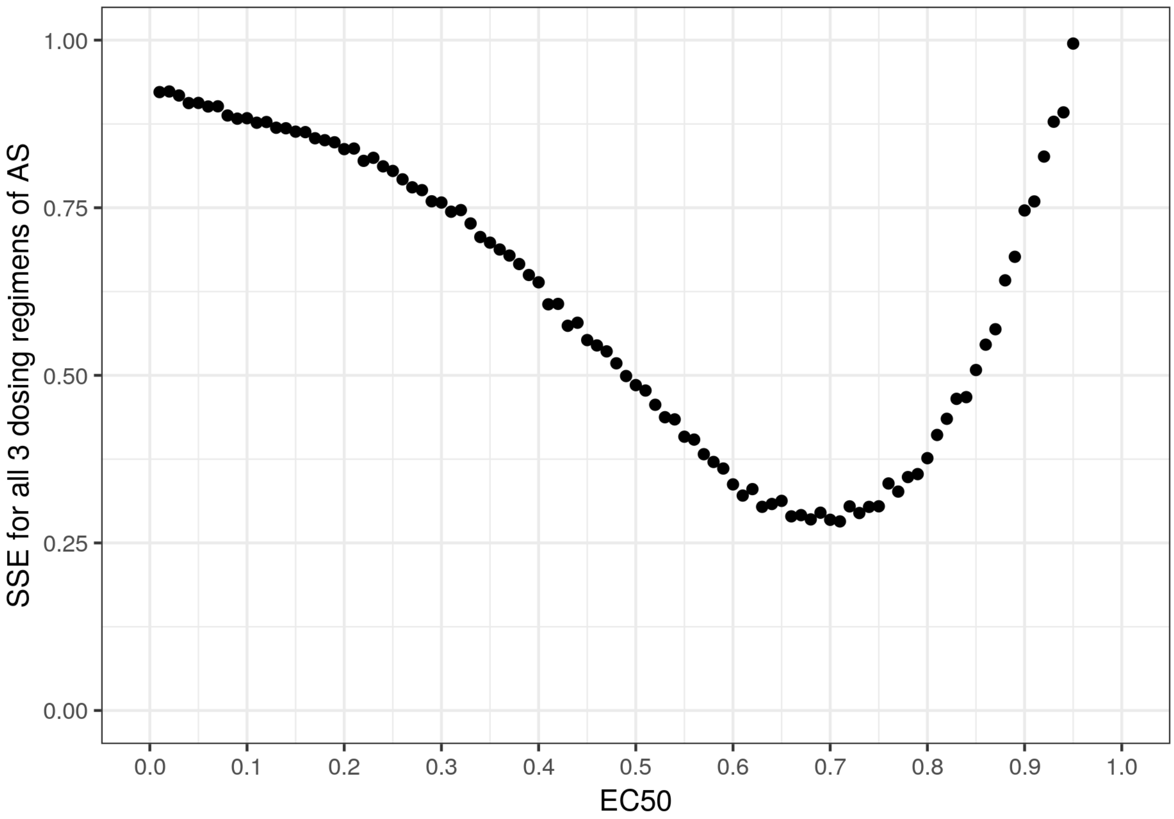


*Figure S1* Sum of squared errors for 3-day, 5-day, and 7-day AS.

| Genotype | AS | LM | AQ | PPQ | MQ | CQ | AL | ASAQ | DHAPPQ | ASMQ |
|---|---|---|---|---|---|---|---|---|---|---|
| KNY--C1 | 0.689 | | | | | | | | | |
| KNY--C2 | 0.689 | | | | | | | | | |
| KNY--Y1 | | | | | | | | | | |
| KNY--Y2 | | | | | | | | | | |
| KYY--C1 | 0.689 | | | | | | | | | |
| KYY--C2 | 0.689 | | | | | | | | | |
| KYY--Y1 | | | | | | | | | | |
| KYY--Y2 | | | | | | | | | | |
| KNF--C1 | 0.689 | | | | | | | | | |
| KNF--C2 | 0.689 | | | | | | | | | |
| KNF--Y1 | | | | | | | | | | |
| KNF--Y2 | | | | | | | | | | |
| KYF--C1 | 0.689 | | | | | | | | | |
| KYF--C2 | 0.689 | | | | | | | | | |
| KYF--Y1 | | | | | | | | | | |
| KYF--Y2 | | | | | | | | | | |
| KNYNYC1 | 0.689 | | | | | | | | | |
| KNYNYC2 | 0.689 | | | | | | | | | |
| KNYNYY1 | | | | | | | | | | |
| KNYNYY2 | | | | | | | | | | |
| KYYYYC1 | 0.689 | | | | | | | | | |
| KYYYYC2 | 0.689 | | | | | | | | | |
| KYYYYY1 | | | | | | | | | | |
| KYYYYY2 | | | | | | | | | | |
| KNFNFC1 | 0.689 | | | | | | | | | |
| KNFNFC2 | 0.689 | | | | | | | | | |
| KNFNFY1 | | | | | | | | | | |
| KNFNFY2 | | | | | | | | | | |
| KYFYFC1 | 0.689 | | | | | | | | | |
| KYFYFC2 | 0.689 | | | | | | | | | |
| KYFYFY1 | | | | | | | | | | |
| KYFYFY2 | | | | | | | | | | |
| TNY--C1 | 0.689 | | | | | | | | | |
| TNY--C2 | 0.689 | | | | | | | | | |
| TNY--Y1 | | | | | | | | | | |
| TNY--Y2 | | | | | | | | | | |
| TYY--C1 | 0.689 | | | | | | | | | |
| TYY--C2 | 0.689 | | | | | | | | | |
| TYY--Y1 | | | | | | | | | | |
| TYY--Y2 | | | | | | | | | | |
| TNF--C1 | 0.689 | | | | | | | | | |
| TNF--C2 | 0.689 | | | | | | | | | |
| TNF--Y1 | | | | | | | | | | |
| TNF--Y2 | | | | | | | | | | |
| TYF--C1 | 0.689 | | | | | | | | | |
| TYF--C2 | 0.689 | | | | | | | | | |
| TYF--Y1 | | | | | | | | | | |
| TYF--Y2 | | | | | | | | | | |
| TNYNYC1 | 0.689 | | | | | | | | | |
| TNYNYC2 | 0.689 | | | | | | | | | |
| TNYNYY1 | | | | | | | | | | |
| TNYNYY2 | | | | | | | | | | |
| TYYYYC1 | 0.689 | | | | | | | | | |
| TYYYYC2 | 0.689 | | | | | | | | | |
| TYYYYY1 | | | | | | | | | | |
| TYYYYY2 | | | | | | | | | | |
| TNFNFC1 | 0.689 | | | | | | | | | |
| TNFNFC2 | 0.689 | | | | | | | | | |
| TNFNFY1 | | | | | | | | | | |
| TNFNFY2 | | | | | | | | | | |
| TYFYFC1 | 0.689 | | | | | | | | | |
| TYFYFC2 | 0.689 | | | | | | | | | |
| TYFYFY1 | | | | | | | | | | |
| TYFYFY2 | | | | | | | | | | |

#### 2.2 Inferring an EC50 for AS and AM on 580Y genotypes

This is done in several stages which are described below.

##### 2.2.1 PPQ monotherapy and DHAPPQ on *kelch13*-C580 and single *plasmepsin-2,3* genotypes

**Cochrane review**. (Zani et al. 2014) reviewed 27 trials comparing DHAPPQ with AL, ASMQ, and ASAQ in Asia, Africa, and South America from 2002 to 2010. In overall, day-28 PCR-corrected cure rate of 3-day DHAPPQ was consistently high (>95%) across studies. This gave us an estimate of DHAPPQ efficacy on 16 artemisinin- and piperaquine-sensitive genotypes (those with *kelch13*-C580 allele and single copy of *plasmepsin-2,3* gene). With the prior knowledge on efficacy of artemisinin monotherapy on *kelch13*-C580 *P.falciparum* (section 2.1 ), we were able to infer efficacy of PPQ monotherapy on these genotypes. By setting EC50 of PPQ to 0.58, we obtained an estimate of 89.9% for day-28 efficacy of PPQ monotherapy on sensitive strains and an estimate of 97.2% efficacy of DHA-PPQ on sensitive genotypes. This PPQ efficacy agrees with some trials of 3-day piperaquine monotherapy (total dose of 1500mg) in China in 1986 which reported day-30 cure rate as high as 86.8% (Lan et al. 1989; Guo and Fu 1989).

| Genotype | AS | LM | AQ | PPQ | MQ | CQ | AL | ASAQ | DHAPPQ | ASMQ |
|---|---|---|---|---|---|---|---|---|---|---|
| KNY--C1 | 0.689 | | | 0.899 | | | | | 0.972 | |
| KNY--C2 | 0.689 | | | | | | | | | |
| KNY--Y1 | | | | 0.899 | | | | | | |
| KNY--Y2 | | | | | | | | | | |
| KYY--C1 | 0.689 | | | 0.899 | | | | | 0.972 | |
| KYY--C2 | 0.689 | | | | | | | | | |
| KYY--Y1 | | | | 0.899 | | | | | | |
| KYY--Y2 | | | | | | | | | | |
| KNF--C1 | 0.689 | | | 0.899 | | | | | 0.972 | |
| KNF--C2 | 0.689 | | | | | | | | | |
| KNF--Y1 | | | | 0.899 | | | | | | |
| KNF--Y2 | | | | | | | | | | |
| KYF--C1 | 0.689 | | | 0.899 | | | | | 0.972 | |
| KYF--C2 | 0.689 | | | | | | | | | |
| KYF--Y1 | | | | 0.899 | | | | | | |
| KYF--Y2 | | | | | | | | | | |
| KNYNYC1 | 0.689 | | | 0.899 | | | | | 0.972 | |
| KNYNYC2 | 0.689 | | | | | | | | | |
| KNYNYY1 | | | | 0.899 | | | | | | |
| KNYNYY2 | | | | | | | | | | |
| KYYYYC1 | 0.689 | | | 0.899 | | | | | 0.972 | |
| KYYYYC2 | 0.689 | | | | | | | | | |
| KYYYYY1 | | | | 0.899 | | | | | | |
| KYYYYY2 | | | | | | | | | | |
| KNFNFC1 | 0.689 | | | 0.899 | | | | | 0.972 | |
| KNFNFC2 | 0.689 | | | | | | | | | |
| KNFNFY1 | | | | 0.899 | | | | | | |
| KNFNFY2 | | | | | | | | | | |
| KYFYFC1 | 0.689 | | | 0.899 | | | | | 0.972 | |
| KYFYFC2 | 0.689 | | | | | | | | | |
| KYFYFY1 | | | | 0.899 | | | | | | |
| KYFYFY2 | | | | | | | | | | |
| TNY--C1 | 0.689 | | | 0.899 | | | | | 0.972 | |
| TNY--C2 | 0.689 | | | | | | | | | |
| TNY--Y1 | | | | 0.899 | | | | | | |
| TNY--Y2 | | | | | | | | | | |
| TYY--C1 | 0.689 | | | 0.899 | | | | | 0.972 | |
| TYY--C2 | 0.689 | | | | | | | | | |
| TYY--Y1 | | | | 0.899 | | | | | | |
| TYY--Y2 | | | | | | | | | | |
| TNF--C1 | 0.689 | | | 0.899 | | | | | 0.972 | |
| TNF--C2 | 0.689 | | | | | | | | | |
| TNF--Y1 | | | | 0.899 | | | | | | |
| TNF--Y2 | | | | | | | | | | |
| TYF--C1 | 0.689 | | | 0.899 | | | | | 0.972 | |
| TYF--C2 | 0.689 | | | | | | | | | |
| TYF--Y1 | | | | 0.899 | | | | | | |
| TYF--Y2 | | | | | | | | | | |
| TNYNYC1 | 0.689 | | | 0.899 | | | | | 0.972 | |
| TNYNYC2 | 0.689 | | | | | | | | | |
| TNYNYY1 | | | | 0.899 | | | | | | |
| TNYNYY2 | | | | | | | | | | |
| TYYYYC1 | 0.689 | | | 0.899 | | | | | 0.972 | |
| TYYYYC2 | 0.689 | | | | | | | | | |
| TYYYYY1 | | | | 0.899 | | | | | | |
| TYYYYY2 | | | | | | | | | | |
| TNFNFC1 | 0.689 | | | 0.899 | | | | | 0.972 | |
| TNFNFC2 | 0.689 | | | | | | | | | |
| TNFNFY1 | | | | 0.899 | | | | | | |
| TNFNFY2 | | | | | | | | | | |
| TYFYFC1 | 0.689 | | | 0.899 | | | | | 0.972 | |
| TYFYFC2 | 0.689 | | | | | | | | | |
| TYFYFY1 | | | | 0.899 | | | | | | |
| TYFYFY2 | | | | | | | | | | |

##### 2.2.2 EC50 for AS on 580Y genotypes and PPQ on multiple-copy *plasmepsin-2,3* genotype

A number of therapeutic efficacy studies in Southeast Asia in late 2000s and 2010s have allowed us to estimate efficacies of DHA-PPQ on resistant genotypes.

**Cambodia 2009-2015**. (Witkowski et al. 2016) used piperaquine survival assay (PSA) (Duru et al. 2015) and whole-genome sequencing technique to analyze samples from Cambodian patients collected from 2009 to 2015 (Leang et al. 2013; 2015) and found multicopy of *plasmepsin-2,3* to be highly associated with PPQ resistance. In the therapeutic efficacy study, patients were treated under supervision with 3-day DHA-PPQ at the total adult dose of around 7 mg/kg DHA + 56 mg/kg PPQ. As (Witkowski et al. 2016) presented in their Figure 5, day-28 PCR-corrected efficacies of DHA-PPQ on *PfK13*-C580 + *PfPM2*-single, *PfK13*-580Y+ *PfPM2*-single, *PfK13*-C580 + *PfPM2*-multi, and *PfK13*-580Y+ *PfPM2*-multi were estimated to be 268/268=100%, 206/208=99%, 12/14=85.7%, and 154/235=65.5%, respectively.

**Cambodia 2011-2013**. (Amato et al. 2017) conducted a genome-wide association study on 297 *P.falciparum*-infected isolates from two parasite clearance rate studies in Cambodia from 2010 to 2013 (Amaratunga et al. 2012; Ashley et al. 2014) and identified *exo*-E415G mutation and amplification of *plasmepsin-2,3* gene as markers for piperaquine resistance. When further investigating the association between these two markers and parasite recrudescence via survival analysis of 133 samples from (Amaratunga et al. 2016) efficacy study, the authors estimated that day-28 efficacy of DHA-PPQ (3-day regimen at total adult dose of around 7 mg/kg DHA + 56 mg/kg PPQ) was 63% [95% CI 50%-70%] in the presence of *plasmepsin-2,3* amplification (Figure 2 (Amato et al. 2017)) and across any *kelch13* allele. In the case of single copy of *plasmepsin2,3*, they estimated DHAPPQ efficacy to be 97% [95% CI 90%-99%].

**Cambodia 2012-2014**. (Saunders et al. 2012; Spring et al. 2015). *P.falciparum* or mixed *P.falciparum*/*P.vivax*, inclusion criterion for parasitemia was 1000-200000 parasites/µL. DHA-PPQ (n=51) vs DHA-PPQ+Primaquine (n=50). 3-day (0h, 24h, 48h) DHA-PPQ (total 360mg DHA + 2880mg PPQ, standard regimen), under supervision. Collected samples were genotyped for *kelch13*-C580Y and *kelch13*-R539T. Estimated mean day to recrudescence in DHA-PPQ was 36 (95% CI 34-39). In Spring et al. (2015)'s Table, among 63 per-protocol patients infected with *kelch13*-580Y, 21 were classified as adequate clinical and parasitological response on day 42, and 37 as *P.falciparum* recrudescence (the parasites also carried MAL10:688956 and MAL13:1718319 SNP); however, it was not clear whether all of these 63 patients were infected with only *P.falciparum* or mixed *P.falciparum*/*P.vivax* at the time of treatment. Our estimate for day-42 cure rate of DHA-PPQ on patients with *kelch13*-580Y is 21/(21+37)=36.2%, and patients without *kelch13*-580Y is 23/(23+5)=82.1%. There was no piperaquine resistance marker reported in the study but in vitro results showed a decrease in piperaquine susceptibility in parasites presented with *kelch13*-580Y.

**Vietnam 2015**. (Phuc et al. 2017) enrolled 46 *P.falciparum*-infected patients (aged 14-53, initial parasitaemia 1514 – 97454 parasites/µL) to a 3-day DHAPPQ study, 44 of whom finished 42-day follow-up. 38/42 had *PfK13*-580Y, 25/46 had multiple copies of *PfPM2*; 22 had both *PfK13*-580Y and multiple copies *PfPM2*. 14 patients experienced recrudescence at day 42; 10 of which carried *PfK13*-580Y and multiple copies *PfPM2*, 3 with only *PfK13*-580Y. Hence, day-42 cure rate of DHAPPQ on *PfK13*-580Y + multiple copies *PfPM2* was (22-10)/22=54.6%, and on *PfK13*-580Y + single copy *PfPM2* (38-22-3)/(38-22)=81.3%.

As previously presented in section 2.1 , we estimated day-28 efficacy of 3-day artesunate monotherapy on *kelch13*-C580 to be 68.9%. Therefore, day-28 efficacy of 3-day DHA-PPQ on *kelch13*-C580 and *plasmepsin2,3*-multicopy genotype should be higher than 68.9%. Additionally, based on analyses from (Witkowski et al. 2016; Amato et al. 2017), we assumed that a plausible range for the value of this particular efficacy could be 70%-86%. By fixing EC50 of AS at 0.75 and varying EC50 of PPQ, we found that PPQ's EC50 value of 1.4 would yield day-28 efficacy of DHA-PPQ on *kelch13*-C580 and multiple copies of *plasmepsin-2,3* within the mentioned range of 70%-86%. Additionally, with EC50 of 1.4, the simulated efficacy of 3-day PPQ monotherapy on this genotype was 21.3%. Since the efficacies of PPQ monotherapy on resistant strains cannot be verified with the available clinical data, we use orange to distinguish them from the others in the following table.

In a similar manner, we assumed the efficacy range of DHA-PPQ on *kelch13*-580Y and single copy of *plasmepsin-2,3* shall be higher than 89.9% (efficacy of PPQ monotherapy on this genotype) and lower than 97% (estimation from (Amato et al. 2017)). We found an EC50 of 1.2 for AS would satisfy this condition, resulting in a 24.1% efficacy of 3-day AS on *kelch13*-580Y.

| Genotype | AS | LM | AQ | PPQ | MQ | CQ | AL | ASAQ | DHAPPQ | ASMQ |
|---|---|---|---|---|---|---|---|---|---|---|
| KNY--C1 | 0.689 | | | 0.899 | | | | | 0.972 | |
| KNY--C2 | 0.689 | | | 0.213 | | | | | 0.768 | |
| KNY--Y1 | | | | 0.899 | | | | | 0.929 | |
| KNY--Y2 | | | | 0.213 | | | | | 0.415 | |
| KYY--C1 | 0.689 | | | 0.899 | | | | | 0.972 | |
| KYY--C2 | 0.689 | | | 0.213 | | | | | 0.768 | |
| KYY--Y1 | | | | 0.899 | | | | | 0.929 | |
| KYY--Y2 | | | | 0.213 | | | | | 0.415 | |
| KNF--C1 | 0.689 | | | 0.899 | | | | | 0.972 | |
| KNF--C2 | 0.689 | | | 0.213 | | | | | 0.768 | |
| KNF--Y1 | | | | 0.899 | | | | | 0.929 | |
| KNF--Y2 | | | | 0.213 | | | | | 0.415 | |
| KYF--C1 | 0.689 | | | 0.899 | | | | | 0.972 | |
| KYF--C2 | 0.689 | | | 0.213 | | | | | 0.768 | |
| KYF--Y1 | | | | 0.899 | | | | | 0.929 | |
| KYF--Y2 | | | | 0.213 | | | | | 0.415 | |
| KNYNYC1 | 0.689 | | | 0.899 | | | | | 0.972 | |
| KNYNYC2 | 0.689 | | | 0.213 | | | | | 0.768 | |
| KNYNYY1 | | | | 0.899 | | | | | 0.929 | |
| KNYNYY2 | | | | 0.213 | | | | | 0.415 | |
| KYYYYC1 | 0.689 | | | 0.899 | | | | | 0.972 | |
| KYYYYC2 | 0.689 | | | 0.213 | | | | | 0.768 | |
| KYYYYY1 | | | | 0.899 | | | | | 0.929 | |
| KYYYYY2 | | | | 0.213 | | | | | 0.415 | |
| KNFNFC1 | 0.689 | | | 0.899 | | | | | 0.972 | |
| KNFNFC2 | 0.689 | | | 0.213 | | | | | 0.768 | |
| KNFNFY1 | | | | 0.899 | | | | | 0.929 | |
| KNFNFY2 | | | | 0.213 | | | | | 0.415 | |
| KYFYFC1 | 0.689 | | | 0.899 | | | | | 0.972 | |
| KYFYFC2 | 0.689 | | | 0.213 | | | | | 0.768 | |
| KYFYFY1 | | | | 0.899 | | | | | 0.929 | |
| KYFYFY2 | | | | 0.213 | | | | | 0.415 | |
| TNY--C1 | 0.689 | | | 0.899 | | | | | 0.972 | |
| TNY--C2 | 0.689 | | | 0.213 | | | | | 0.768 | |
| TNY--Y1 | | | | 0.899 | | | | | 0.929 | |
| TNY--Y2 | | | | 0.213 | | | | | 0.415 | |
| TYY--C1 | 0.689 | | | 0.899 | | | | | 0.972 | |
| TYY--C2 | 0.689 | | | 0.213 | | | | | 0.768 | |
| TYY--Y1 | | | | 0.899 | | | | | 0.929 | |
| TYY--Y2 | | | | 0.213 | | | | | 0.415 | |
| TNF--C1 | 0.689 | | | 0.899 | | | | | 0.972 | |
| TNF--C2 | 0.689 | | | 0.213 | | | | | 0.768 | |
| TNF--Y1 | | | | 0.899 | | | | | 0.929 | |
| TNF--Y2 | | | | 0.213 | | | | | 0.415 | |
| TYF--C1 | 0.689 | | | 0.899 | | | | | 0.972 | |
| TYF--C2 | 0.689 | | | 0.213 | | | | | 0.768 | |
| TYF--Y1 | | | | 0.899 | | | | | 0.929 | |
| TYF--Y2 | | | | 0.213 | | | | | 0.415 | |
| TNYNYC1 | 0.689 | | | 0.899 | | | | | 0.972 | |
| TNYNYC2 | 0.689 | | | 0.213 | | | | | 0.768 | |
| TNYNYY1 | | | | 0.899 | | | | | 0.929 | |
| TNYNYY2 | | | | 0.213 | | | | | 0.415 | |
| TYYYYC1 | 0.689 | | | 0.899 | | | | | 0.972 | |
| TYYYYC2 | 0.689 | | | 0.213 | | | | | 0.768 | |
| TYYYYY1 | | | | 0.899 | | | | | 0.929 | |
| TYYYYY2 | | | | 0.213 | | | | | 0.415 | |
| TNFNFC1 | 0.689 | | | 0.899 | | | | | 0.972 | |
| TNFNFC2 | 0.689 | | | 0.213 | | | | | 0.768 | |
| TNFNFY1 | | | | 0.899 | | | | | 0.929 | |
| TNFNFY2 | | | | 0.213 | | | | | 0.415 | |
| TYFYFC1 | 0.689 | | | 0.899 | | | | | 0.972 | |
| TYFYFC2 | 0.689 | | | 0.213 | | | | | 0.768 | |
| TYFYFY1 | | | | 0.899 | | | | | 0.929 | |
| TYFYFY2 | | | | 0.213 | | | | | 0.415 | |

| Genotype | AS | LM | AQ | PPQ | MQ | CQ | AL | ASAQ | DHAPPQ | ASMQ |
|---|---|---|---|---|---|---|---|---|---|---|
| KNY--C1 | 0.689 | | | 0.899 | | | | | 0.972 | |
| KNY--C2 | 0.689 | | | 0.213 | | | | | 0.768 | |
| KNY--Y1 | 0.241 | | | 0.899 | | | | | 0.929 | |
| KNY--Y2 | 0.241 | | | 0.213 | | | | | 0.415 | |
| KYY--C1 | 0.689 | | | 0.899 | | | | | 0.972 | |
| KYY--C2 | 0.689 | | | 0.213 | | | | | 0.768 | |
| KYY--Y1 | 0.241 | | | 0.899 | | | | | 0.929 | |
| KYY--Y2 | 0.241 | | | 0.213 | | | | | 0.415 | |
| KNF--C1 | 0.689 | | | 0.899 | | | | | 0.972 | |
| KNF--C2 | 0.689 | | | 0.213 | | | | | 0.768 | |
| KNF--Y1 | 0.241 | | | 0.899 | | | | | 0.929 | |
| KNF--Y2 | 0.241 | | | 0.213 | | | | | 0.415 | |
| KYF--C1 | 0.689 | | | 0.899 | | | | | 0.972 | |
| KYF--C2 | 0.689 | | | 0.213 | | | | | 0.768 | |
| KYF--Y1 | 0.241 | | | 0.899 | | | | | 0.929 | |
| KYF--Y2 | 0.241 | | | 0.213 | | | | | 0.415 | |
| KNYNYC1 | 0.689 | | | 0.899 | | | | | 0.972 | |
| KNYNYC2 | 0.689 | | | 0.213 | | | | | 0.768 | |
| KNYNYY1 | 0.241 | | | 0.899 | | | | | 0.929 | |
| KNYNYY2 | 0.241 | | | 0.213 | | | | | 0.415 | |
| KYYYYC1 | 0.689 | | | 0.899 | | | | | 0.972 | |
| KYYYYC2 | 0.689 | | | 0.213 | | | | | 0.768 | |
| KYYYYY1 | 0.241 | | | 0.899 | | | | | 0.929 | |
| KYYYYY2 | 0.241 | | | 0.213 | | | | | 0.415 | |
| KNFNFC1 | 0.689 | | | 0.899 | | | | | 0.972 | |
| KNFNFC2 | 0.689 | | | 0.213 | | | | | 0.768 | |
| KNFNFY1 | 0.241 | | | 0.899 | | | | | 0.929 | |
| KNFNFY2 | 0.241 | | | 0.213 | | | | | 0.415 | |
| KYFYFC1 | 0.689 | | | 0.899 | | | | | 0.972 | |
| KYFYFC2 | 0.689 | | | 0.213 | | | | | 0.768 | |
| KYFYFY1 | 0.241 | | | 0.899 | | | | | 0.929 | |
| KYFYFY2 | 0.241 | | | 0.213 | | | | | 0.415 | |
| TNY--C1 | 0.689 | | | 0.899 | | | | | 0.972 | |
| TNY--C2 | 0.689 | | | 0.213 | | | | | 0.768 | |
| TNY--Y1 | 0.241 | | | 0.899 | | | | | 0.929 | |
| TNY--Y2 | 0.241 | | | 0.213 | | | | | 0.415 | |
| TYY--C1 | 0.689 | | | 0.899 | | | | | 0.972 | |
| TYY--C2 | 0.689 | | | 0.213 | | | | | 0.768 | |
| TYY--Y1 | 0.241 | | | 0.899 | | | | | 0.929 | |
| TYY--Y2 | 0.241 | | | 0.213 | | | | | 0.415 | |
| TNF--C1 | 0.689 | | | 0.899 | | | | | 0.972 | |
| TNF--C2 | 0.689 | | | 0.213 | | | | | 0.768 | |
| TNF--Y1 | 0.241 | | | 0.899 | | | | | 0.929 | |
| TNF--Y2 | 0.241 | | | 0.213 | | | | | 0.415 | |
| TYF--C1 | 0.689 | | | 0.899 | | | | | 0.972 | |
| TYF--C2 | 0.689 | | | 0.213 | | | | | 0.768 | |
| TYF--Y1 | 0.241 | | | 0.899 | | | | | 0.929 | |
| TYF--Y2 | 0.241 | | | 0.213 | | | | | 0.415 | |
| TNYNYC1 | 0.689 | | | 0.899 | | | | | 0.972 | |
| TNYNYC2 | 0.689 | | | 0.213 | | | | | 0.768 | |
| TNYNYY1 | 0.241 | | | 0.899 | | | | | 0.929 | |
| TNYNYY2 | 0.241 | | | 0.213 | | | | | 0.415 | |
| TYYYYC1 | 0.689 | | | 0.899 | | | | | 0.972 | |
| TYYYYC2 | 0.689 | | | 0.213 | | | | | 0.768 | |
| TYYYYY1 | 0.241 | | | 0.899 | | | | | 0.929 | |
| TYYYYY2 | 0.241 | | | 0.213 | | | | | 0.415 | |
| TNFNFC1 | 0.689 | | | 0.899 | | | | | 0.972 | |
| TNFNFC2 | 0.689 | | | 0.213 | | | | | 0.768 | |
| TNFNFY1 | 0.241 | | | 0.899 | | | | | 0.929 | |
| TNFNFY2 | 0.241 | | | 0.213 | | | | | 0.415 | |
| TYFYFC1 | 0.689 | | | 0.899 | | | | | 0.972 | |
| TYFYFC2 | 0.689 | | | 0.213 | | | | | 0.768 | |
| TYFYFY1 | 0.241 | | | 0.899 | | | | | 0.929 | |
| TYFYFY2 | 0.241 | | | 0.213 | | | | | 0.415 | |

### 3 Estimating EC50 for lumefantrine monotherapy using trial data on artemether-lumefantrine (AL) combination therapy

#### 3.1 AL therapeutic efficacy studies

We used two major sources of data to estimate the efficacy of 3-day AL therapy. The first set of sources consisted of major recent reviews of AL therapy (with no genotype information), and the second set of sources included specific trials of AL therapy which also had genotype information for the infecting parasite population (currently, two trials).

A meta-analysis conducted by the Worldwide Antimalarial Resistance Network (WWARN) included 61 trials of AL therapy from 1998 to 2012 with efficacy data on a total of 14,740 patients in various malaria transmission settings (WWARN AL Dose Impact Study Group 2015). Major results of this analysis were the effects of AL dosage and child weight on the treatment efficacy (total 6 doses over 3 days with median total dose of artemether at 11.4 mg/kg and lumefantrine at 68.6 mg/kg). For children under 5 who were not underweight for their age, mean efficacy of AL was estimated to be from 96% to 97% depending on the exact age group. For underweight children, the mean efficacy was reported to be as low as 94% or 95%. Since the trials were conducted prior to 2012, we can assume that all (or the vast majority) of genotypes had the C580 allele on *kelch13* and a single copy of the *plasmepsin-2,3* gene (although multiple copies of *plasmepsin* should not have any effect on AL efficacy).

AL selects for N allele at the N86Y locus and for F allele at the Y184F locus on *pfmdr1* gene and for K allele at the K76T locus on *pfcrt* gene (**Table S1**). Therefore, the genotype that is most sensitive to AL treatment is TYY, with a single copy number of *pfmdr1* gene (TYY--). The next most sensitive genotypes are KYY, TNY, TYF. Among those three loci, the N86Y and K76T locus respectively has the largest and smallest (as seen in (Bassat et al. 2009; Baraka et al. 2015)) effect on dropping AL efficacy. However, when comparing the effect of *pfmdr1* CNV and alleles at locus 86 on predicting recrudescence, the presence of more than one copy of *pfmdr1* produces a higher hazard ratio than the presence of N86 does (Venkatesan et al. 2014).

**Burkina Faso 2005-2006**. (Bassat et al. 2009) AL and DHA-PPQ were given to *P.falciparum*-infected children with mean age of 2.4 years, mean weight of 11 kg, and geometric mean parasitaemia of 24,557 parasites/µL and 25,884 parasites/µL, respectively. AL was administered as 6-dose over 3-day and DHA-PPQ once daily for 3 days under supervision. Overall day-28 PCR-corrected ACPR were 92/97=94.85, and 195/198=98.48; day-42 PCR-corrected ACPR were 89/97=91.75, and 179/198=90.40 for AL, and DHA-PPQ in Burkina Faso. (Baraka et al. 2015) reported the genotyping results on Burkina Faso samples in (Bassat et al. 2009) study. Table 2 in (Baraka et al. 2015) shown that in the AL arm, day-28 efficacy for genotypes with *pfmdr1*-86Y appeared to be 35/35=100% (based on the samples that were sequenced). Day-28 efficacy for *pfmdr1*-N86 was 49/56=87.5%. Day-28 efficacy for *pfcrt*-76T was 57/62=91.9%, and for *pfcrt*-K76 25/27=92.6%. Efficacies on specific genotypes (e.g. *pfcrt*-K76 *pfmdr1*-N86 *pfmdr1*-Y184, *pfcrt*-K76 *pfmdr1*-86Y *pfmdr1*-Y184,...) are not available in this paper, thus, the red numbers in the table on page 17 are approximations.

**Angola 2011-2013**. (Kiaco et al. 2015) conducted a trial with 6-dose AL during high transmission season in Luanda, Angola following WHO 2010 guidelines. Inclusion criteria were age > 6 months, initial *P.falciparum* parasitaemia from 1,000 to 100,000 parasites/µL. Total 103 patients were enrolled, the number of PCR-corrected cured and recrudescent patients by day 28 were 94 and 8, respectively; so overall PCR-corrected efficacy was 94/102=92.2%. Before treatment, 93, 94, and 21 samples were successfully analyzed for *pfmdr1* copy number, polymorphisms on *pfmdr1* gene (at locus 86) and on *kelch13* gene (at loci 493, 539, 543, and 580), respectively. Table 4 in the paper allowed us to infer the effect of SNP on treatment outcomes; however, it seemed the reported "treatment failure" in this Table 4 was PCR-uncorrected recurrent cases (since there were 10 treatment failures carrying *Pfatp6*-S769 which matched the total number of failures reported under PCR-uncorrected section in Table 2). (Gama et al. 2010; Fançony et al. 2012; Ngane et al. 2015) provided information on prevalence of *pfcrt*-K76T, and *pfmdr1*-Y184F in Angola around 2007-2010. We then assumed that samples collected in (Kiaco et al. 2015) were likely to carry *pfcrt*-76T and *pfmdr1*-Y184; hence, we approximated efficacies of artemether-lumefantrine on *pfcrt*-76T *pfmdr1*-N86 *pfmdr1*-Y184 (TNY) and *pfcrt*-76T *pfmdr1*-86Y *pfmdr1*-Y184 (TYY) genotypes to be 63/69=91.3% and 15/17=88.2%, respectively. [Note: no conclusive relationship between genotypes and treatment failure; the study showed LM selecting for *pfmdr1*-86Y, and copy number]

**Angola 2015**. From January to June 2015, (Plucinski et al. 2015) recruited 586 *P.falciparum*-infected children (aged 0.5-12 years old, weighing 6-42 kg, initial parasitaemia 1,003 – 195,529/µL) from Benguela, Zaire, and Lunda Sul Province in Angola into three treatment arms, namely AL, DHA-PPQ, and ASAQ. All treatments were supervised and given according to the standard weight-based dosing regimens from the manufacturers. Overall, day-28 PCR-corrected efficacies of AL were 96.1% in Benguela and 86.5% in Zaire, efficacies of DHA-PPQ were 98.8% in Zaire and 100% in Lunda Sul, efficacies of ASAQ were 99.9% in Benguela and 100% in Lunda Sul. (Ljolje et al. 2018) analyzed 506 pre-treatment and 50 treatment failure samples from the therapeutic efficacy trial above for polymorphisms on *pfmdr1* and *kelch13* genes as well as copy number variation of *pfmdr1*. All genotyped samples in the study carried single copy of *pfmdr1*. The table in the supplementary material of (Ljolje et al. 2018) shown that day-28 PCR-corrected efficacy of AL on *pfmdr1*-N86 *pfmdr1*-Y184 was 92/(92+8)=92%, efficacy on *pfmdr1*-N86 *pfmdr1*-184F was 44/(44+4)=91.7%, efficacy on *pfmdr1*-86Y *pfmdr1*-Y184 was (35+5)/(35+5+1)=97.6%, and efficacy on *pfmdr1*-86Y *pfmdr1*-184F was (5+1)/(5+1+1)=85.7%. Furthermore, according to other molecular marker studies in Angola in early 2010s, the majority of *P.falciparum*-infected Angolan samples carried threonine at codon 76 on *pfcrt* gene; thus, we used these reported cure rates to calibrate our estimates for efficacies of AL on genotypes carrying *pfcrt*-76T. When comparing the first three cure rates mentioned above, namely 92% on TNY--C, 91.7% on TNF--C, and 97.6% on TYY--C, we saw that the reduction in efficacy of AL from the fully sensitive TYY--C to TNY—C (one-mutation away from TYY—C) was much larger than the reduction in efficacy of AL from TNY--C to TNF—C (two-mutation away from TYY—C). This means that either (1) the effect of *pfmdr1*-N86 on susceptibility to LM is greater than that of *pfmdr1*-184F and the interaction between the two alleles is additive, i.e. no epistasis, or (2) both alleles have similar effect on susceptibility to LM and their interaction is one of antagonistic epistasis. In our model, we chose the second approach, i.e. EC50 of LM on TNY--C equals EC50 of LM on TYF--C.

In the table on the next page, among genotypes carrying *kelch13*-C580 allele, we filled in the efficacy of AL on its most sensitive genotype (TYY) and on the three next most sensitive genotypes (one mutation away from TYY), namely, KYY, TNY, and TYF. These four efficacies are shown in red in the table below, and they are duplicated as there is no effect of the *plasmepsin-2,3* gene on AL efficacy.

By setting EC50 of LM to 0.6, we were able to obtain day-28 efficacy of AL on TYY--C1 and TYY--C2 (as in our assumptions, *plasmepsin-2,3* amplification does not confer resistance to either AS or LM) at 96.5%. An EC50 of LM of 0.75 gave us 92.9% efficacy of AL on TNY--C1 and TNY—C2. These two estimates agree with the genotype-stratified efficacies in (Ljolje et al. 2018). As stated previously, we set EC50 of LM on TYF--C to be identical to EC50 of LM on TNY--C which is 0.75, thus, our estimate for efficacy of AL on TYF--C is 92.9%.

When comparing the effect of alleles at locus 76 on *pfcrt* gene on treatment outcomes, it is confirmed that AL selects for *pfcrt*-K76, however, the presence of K76 in the genotype does not seem to substantially increase the number of recrudescence (Bassat et al. 2009; Baraka et al. 2015; Venkatesan et al. 2014; Mwaiswelo et al. 2017). In the drug-by-genotype table, we calibrated EC50 of LM so that the efficacy of AL on genotypes carrying *pfcrt*-K76 would be around 1% lower than the efficacy of the combination on genotypes carrying *pfcrt*-76T to reflect the effect mentioned above. Since we assume that the wild-type *pfcrt*-K76 has the least effect on reducing susceptibility to LM, EC50 of LM on KYY--C1 and KYY--C2 should be higher than 0.6 and lower than 0.75. In our model, this value of EC50 is set to 0.67 which results in 95.3% efficacy of AL on the two corresponding genotypes. This 95.3% efficacy is around 1% lower than the 96.5% efficacies of AL on the fully sensitive genotypes TYY--C1 and TYY--C2.

With values of EC50 as stated above, we were able to simulate for day-28 efficacies of LM monotherapy on the corresponding genotypes. Validation of these EC50 based on limited data from LM monotherapy trials is presented in section 3.2 .

| Genotype | AS | LM | AQ | PPQ | MQ | CQ | AL | ASAQ | DHAPPQ | ASMQ |
|---|---|---|---|---|---|---|---|---|---|---|
| KNY--C1 | 0.689 | | | 0.899 | | | | | 0.972 | |
| KNY--C2 | 0.689 | | | 0.213 | | | | | 0.768 | |
| KNY--Y1 | 0.241 | | | 0.899 | | | | | 0.929 | |
| KNY--Y2 | 0.241 | | | 0.213 | | | | | 0.415 | |
| KYY--C1 | 0.689 | | | 0.899 | | | 0.953 | | 0.972 | |
| KYY--C2 | 0.689 | | | 0.213 | | | 0.953 | | 0.768 | |
| KYY--Y1 | 0.241 | | | 0.899 | | | | | 0.929 | |
| KYY--Y2 | 0.241 | | | 0.213 | | | | | 0.415 | |
| KNF--C1 | 0.689 | | | 0.899 | | | | | 0.972 | |
| KNF--C2 | 0.689 | | | 0.213 | | | | | 0.768 | |
| KNF--Y1 | 0.241 | | | 0.899 | | | | | 0.929 | |
| KNF--Y2 | 0.241 | | | 0.213 | | | | | 0.415 | |
| KYF--C1 | 0.689 | | | 0.899 | | | | | 0.972 | |
| KYF--C2 | 0.689 | | | 0.213 | | | | | 0.768 | |
| KYF--Y1 | 0.241 | | | 0.899 | | | | | 0.929 | |
| KYF--Y2 | 0.241 | | | 0.213 | | | | | 0.415 | |
| KNYNYC1 | 0.689 | | | 0.899 | | | | | 0.972 | |
| KNYNYC2 | 0.689 | | | 0.213 | | | | | 0.768 | |
| KNYNYY1 | 0.241 | | | 0.899 | | | | | 0.929 | |
| KNYNYY2 | 0.241 | | | 0.213 | | | | | 0.415 | |
| KYYYYC1 | 0.689 | | | 0.899 | | | | | 0.972 | |
| KYYYYC2 | 0.689 | | | 0.213 | | | | | 0.768 | |
| KYYYYY1 | 0.241 | | | 0.899 | | | | | 0.929 | |
| KYYYYY2 | 0.241 | | | 0.213 | | | | | 0.415 | |
| KNFNFC1 | 0.689 | | | 0.899 | | | | | 0.972 | |
| KNFNFC2 | 0.689 | | | 0.213 | | | | | 0.768 | |
| KNFNFY1 | 0.241 | | | 0.899 | | | | | 0.929 | |
| KNFNFY2 | 0.241 | | | 0.213 | | | | | 0.415 | |
| KYFYFC1 | 0.689 | | | 0.899 | | | | | 0.972 | |
| KYFYFC2 | 0.689 | | | 0.213 | | | | | 0.768 | |
| KYFYFY1 | 0.241 | | | 0.899 | | | | | 0.929 | |
| KYFYFY2 | 0.241 | | | 0.213 | | | | | 0.415 | |
| TNY--C1 | 0.689 | | | 0.899 | | | 0.929 | | 0.972 | |
| TNY--C2 | 0.689 | | | 0.213 | | | 0.929 | | 0.768 | |
| TNY--Y1 | 0.241 | | | 0.899 | | | | | 0.929 | |
| TNY--Y2 | 0.241 | | | 0.213 | | | | | 0.415 | |
| TYY--C1 | 0.689 | | | 0.899 | | | 0.965 | | 0.972 | |
| TYY--C2 | 0.689 | | | 0.213 | | | 0.965 | | 0.768 | |
| TYY--Y1 | 0.241 | | | 0.899 | | | | | 0.929 | |
| TYY--Y2 | 0.241 | | | 0.213 | | | | | 0.415 | |
| TNF--C1 | 0.689 | | | 0.899 | | | 0.908 | | 0.972 | |
| TNF--C2 | 0.689 | | | 0.213 | | | 0.908 | | 0.768 | |
| TNF--Y1 | 0.241 | | | 0.899 | | | | | 0.929 | |
| TNF--Y2 | 0.241 | | | 0.213 | | | | | 0.415 | |
| TYF--C1 | 0.689 | | | 0.899 | | | 0.929 | | 0.972 | |
| TYF--C2 | 0.689 | | | 0.213 | | | 0.929 | | 0.768 | |
| TYF--Y1 | 0.241 | | | 0.899 | | | | | 0.929 | |
| TYF--Y2 | 0.241 | | | 0.213 | | | | | 0.415 | |
| TNYNYC1 | 0.689 | | | 0.899 | | | | | 0.972 | |
| TNYNYC2 | 0.689 | | | 0.213 | | | | | 0.768 | |
| TNYNYY1 | 0.241 | | | 0.899 | | | | | 0.929 | |
| TNYNYY2 | 0.241 | | | 0.213 | | | | | 0.415 | |
| TYYYYC1 | 0.689 | | | 0.899 | | | | | 0.972 | |
| TYYYYC2 | 0.689 | | | 0.213 | | | | | 0.768 | |
| TYYYYY1 | 0.241 | | | 0.899 | | | | | 0.929 | |
| TYYYYY2 | 0.241 | | | 0.213 | | | | | 0.415 | |
| TNFNFC1 | 0.689 | | | 0.899 | | | | | 0.972 | |
| TNFNFC2 | 0.689 | | | 0.213 | | | | | 0.768 | |
| TNFNFY1 | 0.241 | | | 0.899 | | | | | 0.929 | |
| TNFNFY2 | 0.241 | | | 0.213 | | | | | 0.415 | |
| TYFYFC1 | 0.689 | | | 0.899 | | | | | 0.972 | |
| TYFYFC2 | 0.689 | | | 0.213 | | | | | 0.768 | |
| TYFYFY1 | 0.241 | | | 0.899 | | | | | 0.929 | |
| TYFYFY2 | 0.241 | | | 0.213 | | | | | 0.415 | |

#### 3.2 Validation of Results with Older Trials of Lumefantrine Monotherapy

**China 1994-1996**. Novartis trial. Apart from study AB/MO2 in 1994 which shown the efficacy of 4-dose artemether-lumefantrine to be 100% (PCR-uncorrected), Novartis conducted another factorial trial coded A023 (Novartis Pharmaceuticals Corporation 2009) in China in 1996 to compare artemether-lumefantrine (same regimen as in study AB/MO2) with lumefantrine monotherapy, given either as tablets (total 1920 mg over 48 hours) or capsules (total 2000 mg over 72 hours). 153 *P.falciparum*-infected patients, aged 12-65, weighing 34-70 kg, were recruited to the trial; 52 of whom were assigned to artemether-lumefantrine arm, 51 to tablet lumefantrine, and 50 to capsule lumefantrine. The initial parasitaemia of patients in the three arms ranged from 1026 to 148626 parasites/µL, with geometric mean of 12885, 18695, and 16589 respectively. Day-28 uncorrected cure rate for artemether-lumefantrine was 50/51=98%, tablet lumefantrine 45/49=91.8%, and capsule lumefantrine 47/49=95.9%.

There was no genotyping information in Novartis' FDA application dossier, but we know CQ resistance was widespread in China in the 1990s, hence, it is reasonable to assume that most patients in the Novartis' trials were infected with parasites carrying (*pfcrt*-76T + *pfmdr1*-86Y) genotypes. Furthermore, since CQ does not select for phenylalanine at codon 184 (184F) on *pfmdr1* nor for amplification of this gene and MQ was not used in China in 1990s, it is likely that the *P.falciparum* strains circulating in the region around that time period had TYY-- (*pfcrt*-76T, single copy of *pfmdr1*-86Y and *pfmdr1*-Y184) genotypes.

By setting EC50 of LM on TYY--C (*pfcrt*-76T, single copy of *pfmdr1*-86Y and *pfmdr1*-Y184, *kelch13*-C580) genotypes to 0.6, we were able to obtain day-28 efficacy of AL to match previously presented data at 96.5% (filled in red in the table on page 18). This value of EC50 also gave us day-28 efficacy of 3-day LM monotherapy on the most LM-sensitive genotypes (TYY--) of 87.0% which is close to the reported cure rates in Novartis' trial A023 in China from 1994 to 1996. In a similar approach, we were able to estimate day-3 efficacies of LM monotherapy on TNY, TNF, and KYY to be 70%, 70%, and 82.8% (filled in orange in the table on page 20), respectively. Furthermore, since alleles on *kelch13* do not affect efficacies of LM monotherapy, we could use the latest EC50 of LM to simulate for efficacies of AL combination on artemisinin-resistant genotypes (i.e. TYY--Y, TNY--Y, TYY--Y, KYY--Y in the table on page 20)

| Genotype | AS | LM | AQ | PPQ | MQ | CQ | AL | ASAQ | DHAPPQ | ASMQ |
|---|---|---|---|---|---|---|---|---|---|---|
| KNY--C1 | 0.689 | | | 0.899 | | | | | 0.972 | |
| KNY--C2 | 0.689 | | | 0.213 | | | | | 0.768 | |
| KNY--Y1 | 0.241 | | | 0.899 | | | | | 0.929 | |
| KNY--Y2 | 0.241 | | | 0.213 | | | | | 0.415 | |
| KYY--C1 | 0.689 | 0.828 | | 0.899 | | | 0.953 | | 0.972 | |
| KYY--C2 | 0.689 | 0.828 | | 0.213 | | | 0.953 | | 0.768 | |
| KYY--Y1 | 0.241 | | | 0.899 | | | | | 0.929 | |
| KYY--Y2 | 0.241 | | | 0.213 | | | | | 0.415 | |
| KNF--C1 | 0.689 | | | 0.899 | | | | | 0.972 | |
| KNF--C2 | 0.689 | | | 0.213 | | | | | 0.768 | |
| KNF--Y1 | 0.241 | | | 0.899 | | | | | 0.929 | |
| KNF--Y2 | 0.241 | | | 0.213 | | | | | 0.415 | |
| KYF--C1 | 0.689 | | | 0.899 | | | | | 0.972 | |
| KYF--C2 | 0.689 | | | 0.213 | | | | | 0.768 | |
| KYF--Y1 | 0.241 | | | 0.899 | | | | | 0.929 | |
| KYF--Y2 | 0.241 | | | 0.213 | | | | | 0.415 | |
| KNYNYC1 | 0.689 | | | 0.899 | | | | | 0.972 | |
| KNYNYC2 | 0.689 | | | 0.213 | | | | | 0.768 | |
| KNYNYY1 | 0.241 | | | 0.899 | | | | | 0.929 | |
| KNYNYY2 | 0.241 | | | 0.213 | | | | | 0.415 | |
| KYYYYC1 | 0.689 | | | 0.899 | | | | | 0.972 | |
| KYYYYC2 | 0.689 | | | 0.213 | | | | | 0.768 | |
| KYYYYY1 | 0.241 | | | 0.899 | | | | | 0.929 | |
| KYYYYY2 | 0.241 | | | 0.213 | | | | | 0.415 | |
| KNFNFC1 | 0.689 | | | 0.899 | | | | | 0.972 | |
| KNFNFC2 | 0.689 | | | 0.213 | | | | | 0.768 | |
| KNFNFY1 | 0.241 | | | 0.899 | | | | | 0.929 | |
| KNFNFY2 | 0.241 | | | 0.213 | | | | | 0.415 | |
| KYFYFC1 | 0.689 | | | 0.899 | | | | | 0.972 | |
| KYFYFC2 | 0.689 | | | 0.213 | | | | | 0.768 | |
| KYFYFY1 | 0.241 | | | 0.899 | | | | | 0.929 | |
| KYFYFY2 | 0.241 | | | 0.213 | | | | | 0.415 | |
| TNY--C1 | 0.689 | 0.770 | | 0.899 | | | 0.929 | | 0.972 | |
| TNY--C2 | 0.689 | 0.770 | | 0.213 | | | 0.929 | | 0.768 | |
| TNY--Y1 | 0.241 | | | 0.899 | | | | | 0.929 | |
| TNY--Y2 | 0.241 | | | 0.213 | | | | | 0.415 | |
| TYY--C1 | 0.689 | 0.870 | | 0.899 | | | 0.965 | | 0.972 | |
| TYY--C2 | 0.689 | 0.870 | | 0.213 | | | 0.965 | | 0.768 | |
| TYY--Y1 | 0.241 | 0.870 | | 0.899 | | | | | 0.929 | |
| TYY--Y2 | 0.241 | 0.870 | | 0.213 | | | | | 0.415 | |
| TNF--C1 | 0.689 | 0.676 | | 0.899 | | | 0.908 | | 0.972 | |
| TNF--C2 | 0.689 | 0.676 | | 0.213 | | | 0.908 | | 0.768 | |
| TNF--Y1 | 0.241 | | | 0.899 | | | | | 0.929 | |
| TNF--Y2 | 0.241 | | | 0.213 | | | | | 0.415 | |
| TYF--C1 | 0.689 | 0.770 | | 0.899 | | | 0.929 | | 0.972 | |
| TYF--C2 | 0.689 | 0.770 | | 0.213 | | | 0.929 | | 0.768 | |
| TYF--Y1 | 0.241 | | | 0.899 | | | | | 0.929 | |
| TYF--Y2 | 0.241 | | | 0.213 | | | | | 0.415 | |
| TNYNYC1 | 0.689 | | | 0.899 | | | | | 0.972 | |
| TNYNYC2 | 0.689 | | | 0.213 | | | | | 0.768 | |
| TNYNYY1 | 0.241 | | | 0.899 | | | | | 0.929 | |
| TNYNYY2 | 0.241 | | | 0.213 | | | | | 0.415 | |
| TYYYYC1 | 0.689 | | | 0.899 | | | | | 0.972 | |
| TYYYYC2 | 0.689 | | | 0.213 | | | | | 0.768 | |
| TYYYYY1 | 0.241 | | | 0.899 | | | | | 0.929 | |
| TYYYYY2 | 0.241 | | | 0.213 | | | | | 0.415 | |
| TNFNFC1 | 0.689 | | | 0.899 | | | | | 0.972 | |
| TNFNFC2 | 0.689 | | | 0.213 | | | | | 0.768 | |
| TNFNFY1 | 0.241 | | | 0.899 | | | | | 0.929 | |
| TNFNFY2 | 0.241 | | | 0.213 | | | | | 0.415 | |
| TYFYFC1 | 0.689 | | | 0.899 | | | | | 0.972 | |
| TYFYFC2 | 0.689 | | | 0.213 | | | | | 0.768 | |
| TYFYFY1 | 0.241 | | | 0.899 | | | | | 0.929 | |
| TYFYFY2 | 0.241 | | | 0.213 | | | | | 0.415 | |

| Genotype | AS | LM | AQ | PPQ | MQ | CQ | AL | ASAQ | DHAPPQ | ASMQ |
|---|---|---|---|---|---|---|---|---|---|---|
| KNY--C1 | 0.689 | | | 0.899 | | | | | 0.972 | |
| KNY--C2 | 0.689 | | | 0.213 | | | | | 0.768 | |
| KNY--Y1 | 0.241 | | | 0.899 | | | | | 0.929 | |
| KNY--Y2 | 0.241 | | | 0.213 | | | | | 0.415 | |
| KYY--C1 | 0.689 | 0.828 | | 0.899 | | | 0.953 | | 0.972 | |
| KYY--C2 | 0.689 | 0.828 | | 0.213 | | | 0.953 | | 0.768 | |
| KYY--Y1 | 0.241 | 0.828 | | 0.899 | | | 0.878 | | 0.929 | |
| KYY--Y2 | 0.241 | 0.828 | | 0.213 | | | 0.878 | | 0.415 | |
| KNF--C1 | 0.689 | | | 0.899 | | | | | 0.972 | |
| KNF--C2 | 0.689 | | | 0.213 | | | | | 0.768 | |
| KNF--Y1 | 0.241 | | | 0.899 | | | | | 0.929 | |
| KNF--Y2 | 0.241 | | | 0.213 | | | | | 0.415 | |
| KYF--C1 | 0.689 | | | 0.899 | | | | | 0.972 | |
| KYF--C2 | 0.689 | | | 0.213 | | | | | 0.768 | |
| KYF--Y1 | 0.241 | | | 0.899 | | | | | 0.929 | |
| KYF--Y2 | 0.241 | | | 0.213 | | | | | 0.415 | |
| KNYNYC1 | 0.689 | | | 0.899 | | | | | 0.972 | |
| KNYNYC2 | 0.689 | | | 0.213 | | | | | 0.768 | |
| KNYNYY1 | 0.241 | | | 0.899 | | | | | 0.929 | |
| KNYNYY2 | 0.241 | | | 0.213 | | | | | 0.415 | |
| KYYYYC1 | 0.689 | | | 0.899 | | | | | 0.972 | |
| KYYYYC2 | 0.689 | | | 0.213 | | | | | 0.768 | |
| KYYYYY1 | 0.241 | | | 0.899 | | | | | 0.929 | |
| KYYYYY2 | 0.241 | | | 0.213 | | | | | 0.415 | |
| KNFNFC1 | 0.689 | | | 0.899 | | | | | 0.972 | |
| KNFNFC2 | 0.689 | | | 0.213 | | | | | 0.768 | |
| KNFNFY1 | 0.241 | | | 0.899 | | | | | 0.929 | |
| KNFNFY2 | 0.241 | | | 0.213 | | | | | 0.415 | |
| KYFYFC1 | 0.689 | | | 0.899 | | | | | 0.972 | |
| KYFYFC2 | 0.689 | | | 0.213 | | | | | 0.768 | |
| KYFYFY1 | 0.241 | | | 0.899 | | | | | 0.929 | |
| KYFYFY2 | 0.241 | | | 0.213 | | | | | 0.415 | |
| TNY--C1 | 0.689 | 0.770 | | 0.899 | | | 0.929 | | 0.972 | |
| TNY--C2 | 0.689 | 0.770 | | 0.213 | | | 0.929 | | 0.768 | |
| TNY--Y1 | 0.241 | 0.770 | | 0.899 | | | 0.829 | | 0.929 | |
| TNY--Y2 | 0.241 | 0.770 | | 0.213 | | | 0.829 | | 0.415 | |
| TYY--C1 | 0.689 | 0.870 | | 0.899 | | | 0.965 | | 0.972 | |
| TYY--C2 | 0.689 | 0.870 | | 0.213 | | | 0.965 | | 0.768 | |
| TYY--Y1 | 0.241 | 0.870 | | 0.899 | | | 0.908 | | 0.929 | |
| TYY--Y2 | 0.241 | 0.870 | | 0.213 | | | 0.908 | | 0.415 | |
| TNF--C1 | 0.689 | 0.676 | | 0.899 | | | 0.908 | | 0.972 | |
| TNF--C2 | 0.689 | 0.676 | | 0.213 | | | 0.908 | | 0.768 | |
| TNF--Y1 | 0.241 | 0.676 | | 0.899 | | | 0.753 | | 0.929 | |
| TNF--Y2 | 0.241 | 0.676 | | 0.213 | | | 0.753 | | 0.415 | |
| TYF--C1 | 0.689 | 0.770 | | 0.899 | | | 0.929 | | 0.972 | |
| TYF--C2 | 0.689 | 0.770 | | 0.213 | | | 0.929 | | 0.768 | |
| TYF--Y1 | 0.241 | 0.770 | | 0.899 | | | 0.829 | | 0.929 | |
| TYF--Y2 | 0.241 | 0.770 | | 0.213 | | | 0.829 | | 0.415 | |
| TNYNYC1 | 0.689 | | | 0.899 | | | | | 0.972 | |
| TNYNYC2 | 0.689 | | | 0.213 | | | | | 0.768 | |
| TNYNYY1 | 0.241 | | | 0.899 | | | | | 0.929 | |
| TNYNYY2 | 0.241 | | | 0.213 | | | | | 0.415 | |
| TYYYYC1 | 0.689 | | | 0.899 | | | | | 0.972 | |
| TYYYYC2 | 0.689 | | | 0.213 | | | | | 0.768 | |
| TYYYYY1 | 0.241 | | | 0.899 | | | | | 0.929 | |
| TYYYYY2 | 0.241 | | | 0.213 | | | | | 0.415 | |
| TNFNFC1 | 0.689 | | | 0.899 | | | | | 0.972 | |
| TNFNFC2 | 0.689 | | | 0.213 | | | | | 0.768 | |
| TNFNFY1 | 0.241 | | | 0.899 | | | | | 0.929 | |
| TNFNFY2 | 0.241 | | | 0.213 | | | | | 0.415 | |
| TYFYFC1 | 0.689 | | | 0.899 | | | | | 0.972 | |
| TYFYFC2 | 0.689 | | | 0.213 | | | | | 0.768 | |
| TYFYFY1 | 0.241 | | | 0.899 | | | | | 0.929 | |
| TYFYFY2 | 0.241 | | | 0.213 | | | | | 0.415 | |

Next, we consider genotypes with two alleles conferring resistance to LM, namely TNF, KNY, and KYF.

An EC50 of LM of 0.85 gives us 90.8% efficacy of AL on TNF--C1 and TNF--C2 which is in line with results from (Ljolje et al. 2018) as summarized in section 3.1 . We do not have good empirical data for efficacies of AL on the KNY--C genotype but we expect the presence of *pfmdr1*-N86 allele would drop the efficacy of AL by 5% (in absolute scale) from what the AL efficacy should be in the case of *pfmdr1*-86Y as observed in previous trials (Venkatesan et al. 2014). Therefore, we could approximate that day-28 efficacies of AL combination and LM monotherapy on KNY--C would be 91.5% and 71.9%, respectively (table on page 22). For efficacy of AL on KYF--C, we set EC50 of LM at 0.8 to obtain day-28 cure rate of AL at 91.5%, which is around 1% lower than the estimated efficacy of AL on TYF--C (see section 3.1 for the effect of *pfcrt*-K76 on susceptibility to LM).

Similarly, we do not have data for efficacy of AL on KNF--C, which is three-mutation away from the fully sensitive TYY--C, so we calibrated EC50 of LM on KNF--C based on EC50 of LM on TNF--C. Specifically, we set EC50 of LM on KNF--C to be 0.9 so that day-28 cure rate of AL on KNF--C reaches 89.0%, which is around 1% lower than day-28 efficacy of AL on TNF--C.

We do not have much evidence for efficacies of AL on genotypes carrying multiple copies of *pfmdr1* gene, but we expect the effect of *pfmdr1* amplification on reducing susceptibility to LM to be large (Venkatesan et al. 2014). In our model, we calibrated EC50 of LM on *pfmdr1* multi-copy genotypes based on EC50 of LM on *pfmdr1* single-copy counterparts so that the resulted efficacies of AL on *pfmdr1* multi-copy genotypes shall be 5-6% lower than those on *pfmdr1* single-copy. Table on page 22 shows all estimates for efficacies of LM and AL on all 64 genotypes.

| Genotype | AS | LM | AQ | PPQ | MQ | CQ | AL | ASAQ | DHAPPQ | ASMQ |
|---|---|---|---|---|---|---|---|---|---|---|
| KNY--C1 | 0.689 | 0.719 | | 0.899 | | | 0.915 | | 0.972 | |
| KNY--C2 | 0.689 | 0.719 | | 0.213 | | | 0.915 | | 0.768 | |
| KNY--Y1 | 0.241 | 0.719 | | 0.899 | | | 0.795 | | 0.929 | |
| KNY--Y2 | 0.241 | 0.719 | | 0.213 | | | 0.795 | | 0.415 | |
| KYY--C1 | 0.689 | 0.828 | | 0.899 | | | 0.953 | | 0.972 | |
| KYY--C2 | 0.689 | 0.828 | | 0.213 | | | 0.953 | | 0.768 | |
| KYY--Y1 | 0.241 | 0.828 | | 0.899 | | | 0.878 | | 0.929 | |
| KYY--Y2 | 0.241 | 0.828 | | 0.213 | | | 0.878 | | 0.415 | |
| KNF--C1 | 0.689 | 0.627 | | 0.899 | | | 0.890 | | 0.972 | |
| KNF--C2 | 0.689 | 0.627 | | 0.213 | | | 0.890 | | 0.768 | |
| KNF--Y1 | 0.241 | 0.627 | | 0.899 | | | 0.723 | | 0.929 | |
| KNF--Y2 | 0.241 | 0.627 | | 0.213 | | | 0.723 | | 0.415 | |
| KYF--C1 | 0.689 | 0.719 | | 0.899 | | | 0.915 | | 0.972 | |
| KYF--C2 | 0.689 | 0.719 | | 0.213 | | | 0.915 | | 0.768 | |
| KYF--Y1 | 0.241 | 0.719 | | 0.899 | | | 0.795 | | 0.929 | |
| KYF--Y2 | 0.241 | 0.719 | | 0.213 | | | 0.795 | | 0.415 | |
| KNYNYC1 | 0.689 | 0.523 | | 0.899 | | | 0.859 | | 0.972 | |
| KNYNYC2 | 0.689 | 0.523 | | 0.213 | | | 0.859 | | 0.768 | |
| KNYNYY1 | 0.241 | 0.523 | | 0.899 | | | 0.646 | | 0.929 | |
| KNYNYY2 | 0.241 | 0.523 | | 0.213 | | | 0.646 | | 0.415 | |
| KYYYYC1 | 0.689 | 0.662 | | 0.899 | | | 0.897 | | 0.972 | |
| KYYYYC2 | 0.689 | 0.662 | | 0.213 | | | 0.897 | | 0.768 | |
| KYYYYY1 | 0.241 | 0.662 | | 0.899 | | | 0.752 | | 0.929 | |
| KYYYYY2 | 0.241 | 0.662 | | 0.213 | | | 0.752 | | 0.415 | |
| KNFNFC1 | 0.689 | 0.422 | | 0.899 | | | 0.830 | | 0.972 | |
| KNFNFC2 | 0.689 | 0.422 | | 0.213 | | | 0.830 | | 0.768 | |
| KNFNFY1 | 0.241 | 0.422 | | 0.899 | | | 0.570 | | 0.929 | |
| KNFNFY2 | 0.241 | 0.422 | | 0.213 | | | 0.570 | | 0.415 | |
| KYFYFC1 | 0.689 | 0.523 | | 0.899 | | | 0.859 | | 0.972 | |
| KYFYFC2 | 0.689 | 0.523 | | 0.213 | | | 0.859 | | 0.768 | |
| KYFYFY1 | 0.241 | 0.523 | | 0.899 | | | 0.646 | | 0.929 | |
| KYFYFY2 | 0.241 | 0.523 | | 0.213 | | | 0.646 | | 0.415 | |
| TNY--C1 | 0.689 | 0.770 | | 0.899 | | | 0.929 | | 0.972 | |
| TNY--C2 | 0.689 | 0.770 | | 0.213 | | | 0.929 | | 0.768 | |
| TNY--Y1 | 0.241 | 0.770 | | 0.899 | | | 0.829 | | 0.929 | |
| TNY--Y2 | 0.241 | 0.770 | | 0.213 | | | 0.829 | | 0.415 | |
| TYY--C1 | 0.689 | 0.870 | | 0.899 | | | 0.965 | | 0.972 | |
| TYY--C2 | 0.689 | 0.870 | | 0.213 | | | 0.965 | | 0.768 | |
| TYY--Y1 | 0.241 | 0.870 | | 0.899 | | | 0.908 | | 0.929 | |
| TYY--Y2 | 0.241 | 0.870 | | 0.213 | | | 0.908 | | 0.415 | |
| TNF--C1 | 0.689 | 0.676 | | 0.899 | | | 0.908 | | 0.972 | |
| TNF--C2 | 0.689 | 0.676 | | 0.213 | | | 0.908 | | 0.768 | |
| TNF--Y1 | 0.241 | 0.676 | | 0.899 | | | 0.753 | | 0.929 | |
| TNF--Y2 | 0.241 | 0.676 | | 0.213 | | | 0.753 | | 0.415 | |
| TYF--C1 | 0.689 | 0.770 | | 0.899 | | | 0.929 | | 0.972 | |
| TYF--C2 | 0.689 | 0.770 | | 0.213 | | | 0.929 | | 0.768 | |
| TYF--Y1 | 0.241 | 0.770 | | 0.899 | | | 0.829 | | 0.929 | |
| TYF--Y2 | 0.241 | 0.770 | | 0.213 | | | 0.829 | | 0.415 | |
| TNYNYC1 | 0.689 | 0.575 | | 0.899 | | | 0.869 | | 0.972 | |
| TNYNYC2 | 0.689 | 0.575 | | 0.213 | | | 0.869 | | 0.768 | |
| TNYNYY1 | 0.241 | 0.575 | | 0.899 | | | 0.684 | | 0.929 | |
| TNYNYY2 | 0.241 | 0.575 | | 0.213 | | | 0.684 | | 0.415 | |
| TYYYYC1 | 0.689 | 0.719 | | 0.899 | | | 0.915 | | 0.972 | |
| TYYYYC2 | 0.689 | 0.719 | | 0.213 | | | 0.915 | | 0.768 | |
| TYYYYY1 | 0.241 | 0.719 | | 0.899 | | | 0.795 | | 0.929 | |
| TYYYYY2 | 0.241 | 0.719 | | 0.213 | | | 0.795 | | 0.415 | |
| TNFNFC1 | 0.689 | 0.474 | | 0.899 | | | 0.844 | | 0.972 | |
| TNFNFC2 | 0.689 | 0.474 | | 0.213 | | | 0.844 | | 0.768 | |
| TNFNFY1 | 0.241 | 0.474 | | 0.899 | | | 0.604 | | 0.929 | |
| TNFNFY2 | 0.241 | 0.474 | | 0.213 | | | 0.604 | | 0.415 | |
| TYFYFC1 | 0.689 | 0.575 | | 0.899 | | | 0.869 | | 0.972 | |
| TYFYFC2 | 0.689 | 0.575 | | 0.213 | | | 0.869 | | 0.768 | |
| TYFYFY1 | 0.241 | 0.575 | | 0.899 | | | 0.684 | | 0.929 | |
| TYFYFY2 | 0.241 | 0.575 | | 0.213 | | | 0.684 | | 0.415 | |

### 4 MQ, ASMQ, CQ

#### 4.1 MQ, ASMQ

Cochrane review 2005: ASMQ vs MQ. 8 trials, 1996 patients, low transmission settings (South-East Asia, Peruvian Amazon)

**Hainan Island (China) 1982-1984.** In Hainan Island, China, from 1982 to 1984, (Li et al. 1984) recruited 80 *P. falciparum*-infected patients, aged 9-57, into a 4-arm trial to compare efficacy of mefloquine plus Fansidar (group A), mefloquine plus qinghaosu (group B), mefloquine plus Fansidar and qinghaosu (group C), and qinghaosu alone (group D); each arm had 20 patients. The initial parasitaemia ranged from 1,840 to 353,157 parasite/µl, with mean of 57,414 parasite/µl. All regimens were given as one single dose except for group D which was 3-dose over 3 days. The total dose of mefloquine was 750mg, and that of Fansidar was 75 mg pyrimerthamine plus 1500 mg sulfadoxine in all regimens. Total dose of qinghaosu was 1000 mg in group B, and C, and 2000 mg in group D. Day-28 radical cure rate, after excluding vivax cases, were 100% for mefloquine-fansidar, 100% for mefloquine-qinghaosu, 100% for mefloquine-fansidar-qinghaosu, and 10/17=58.8% for qinghaosu 3-day monotherapy.

**Thailand 1980-1981**. (Harinasuta, Bunnag, and Wernsdorfer 1983) MQ on CQ-resistant, SP-resistant strain. 150 *P. falciparum*-confirmed (unknown *P. vivax* coinfection status) males, aged 15-65, weighing 42-73 kg, were enrolled in the study, 118 of whom completed 63-day follow-up. The three treatment arms administered a single dose of oral MQ at 500, 750, and 1000 mg to patients under supervision. Mean initial parasitaemia in three arms were 58,844, 42,679, 39,356 parasites/µl, respectively. *P. vivax* relapses were observed in all arms after day 28. When excluding *P. vivax* relapses, day-63 MQ cure rates were (38-11)/(40-11)=93.1% in 500mg arm, (37-16)/(40-16)=87.5% in 750mg arm, (38-16)/(38-16)=100% in 1000mg. However, day-28 efficacies were 38/40=95.0%, 37/40=92.5%, and 38/38=100% in the three arms.

**Thailand 1991 or 1992.** (Karbwang et al. 1992) conducted a trial at Hospital for Tropical Diseases in Bangkok, Thailand to compare efficacy of 1-day mefloquine monotherapy and 5-day oral artemether monotherapy. Total dose in each arm were 1250mg and 700mg respectively. Patients in the study were 46 males aged 15-50, weighing 45-65 kg, with acute uncomplicated *P. falciparum* (no information was given on whether the *P. falciparum* infection was confirmed or suspected). 12 patients were assigned to mefloquine arm, and 34 to artemether. Parasitaemia were ranging from 3,900 to 149,260 parasite/µl; geometric mean of initial parasitaemia were 23,438 parasite/µl for mefloquine arm and 13,490 for artemether. Drugs were given at 0h, 6h, 30h, 54h, 78h, and 102h. The outcomes were assessed based on WHO criteria used at the time (World Health Organization 1973), and treatment success was defined as parasitological cure on day 28. Efficacy of 5-day artemether was 29/30 = 96.7% (29/30); day-28 efficacy for mefloquine arm was 8/11 = 73%, (day-42: 7/11=64%).

**Thailand 1992.** (Looareesuwan et al. 1992) conducted a study to evaluate the efficacies of artesunate (AS), mefloquine (MQ) and artesunate mefloquine (ASMQ) combinations in Thailand in 1991. 127 patients were recruited and assigned to three different treatment arms: oral AS (100mg immediately, then 50mg every 12h for 5 days, total dose 600mg), oral MQ (750mg immediately, then 500mg at 6h; total dose 1250mg), and ASMQ (same oral 5-day AS regimen; two doses of oral MQ given on day 6 as in MQ arm). The baseline parasitaemia ranged from 172 to 184,400/µL, with geometric mean between 14,195/µl to 25,825/µl. The patients were 16-60 years old, weighing 45-60kg, and agreed to remain in the hospital for 28 days. 28-day efficacies, measured as parasitological cure, were 88% for 5-day mono AS (n=40), 81% for MQ (n=37), and 100% for ASMQ (n=39).

**Thailand 1993-1994**. To study the efficacy of combinations of artemisinin derivatives and mefloquine versus monotherapy of mefloquine, (R. N. Price et al. 1995) recruited 540 *P.falciparum*-infected patients with mean age of 17.6-20.2 years and mean initial parasitaemia of 3,003 – 7,178/µL from the Karen ethnic minority camp along Thailand-Myanmar border from June 1993 to May 1994. The three treatment regimens were (i) single dose 25 mg/kg MQ, (ii) 3-day 4 mg/kg AS plus single dose 25 mg/kg MQ on day 2, and (iii) 3-day 4 mg/kg AM plus single dose 25 mg/kg MQ on day 2. After adjusting for reinfection, day-28 efficacy of MQ monotherapy was 100-34.8=65.2%, efficacy of ASMQ was 100-3.9=96.1%, and efficacy of AMMQ was 100-2.8=97.2%.

**Thailand 1995-1996.** (Looareesuwan et al. 1999) recruited 252 *P.falciparum*-confirmed patients, aged 13-63, weighing 35-107 kg, at Bangkok Hospital of Tropical Diseases (Thailand) from 1995 to 1996 to compare the effect of artemether-lumefantrine (4 doses, 80 mg artemether plus 480 mg lumefantrine each, given at 0h, 8h, 24h, and 48h) against mefloquine (750 mg at 0h, then 500 mg at 8h). The initial parasitaemia ranged from 1,018 to 295,260 parasites/µL, geometric mean was 16,246 and 17,792 parasites/µL in artemether-lumefantrine and mefloquine arm respectively. The protocol required the patients to remain in the hospital for 28 days to avoid reinfection. Day-28 cure rates were 79/114=69.3% in artemether-lumefantrine arm, and 98/119=82.4% in mefloquine arm. Novartis NDA referred to this trial as study A004 in their FDA application for Coartem (Novartis Pharmaceuticals Corporation 2009).

**Thailand 1995-1996.** From 1995 to 1996, 617 *P. falciparum*-confirmed patients, aged 5-66, weighing 12-72 kg, were enrolled in a comparison trial of artemether-lumefantrine (n=309) with artesunate-mefloquine (n=308) in Mae La camp, Thailand (van Vugt et al. 1998). The patients' initial parasitaemia ranged from 10 to 512,297 parasites/µL with geometric mean of 4,456 parasites/µL. In the first arm, each dose of artemether-lumefantrine (1-2 mg artemether per kg plus 6-12 mg lumefantrine per kg) was given at 0h, 8h, 24h, and 48h. In the other arm, patients received 3 single daily doses of artesunate, 4 mg/kg each, plus mefloquine at 15 mg/kg on day 2 and at 10 mg/kg on day 3. Day-63 PCR-corrected cure rates were 187/223=83.9% for artemether-lumefantrine, and 202/211=95.7% for artesunate-mefloquine. Novartis NDA referred to this trial as study A008 in their FDA application for Coartem (Novartis Pharmaceuticals Corporation 2009).

**Thailand 1997-1998.** From 1997 to 1998, (van Vugt et al. 2000) enrolled 200 *P. falciparum*-confirmed patients, aged 2-63, weighing 8-81 kg, initial parasitaemia 264-254,490 parasites/µL, in Bangkok Hospital of Tropical Diseases and Mae La camp (Thailand) to a comparison trial of 3-day 6-dose artemether-lumefantrine with 3-day artesunate-mefloquine. Artemether-lumefantrine and artesunate-mefloquine regimens were identical to those in arm 2 in (van Vugt et al. 1998; 1999). Day-28 PCR-corrected cure rates were 130/133=97.7% in artemether-lumefantrine and 47/47=100% artesunate-mefloquine group. Novartis NDA referred to this trial as study A026 in their FDA application for Coartem (Novartis Pharmaceuticals Corporation 2009).

**Thailand 1992-1993**. From January 1992 to June 1993, (Nosten et al. 1994) conducted two studies in the Karen ethnic minority camp along Thailand-Myanmar border to determine suitable treatment for multi-drug resistant (to CQ, SP, and MQ) *P. falciparum* strains which were circulating in the area in the late 1980s-early 1990s. One of the arms in both studies was single dose MQ at 25 mg/kg. In study 1, the active comparator was single dose MQ at 25 mg/kg plus single dose AS at 4 mg/kg. In study 2, the active comparator was 3-day AS at total dose of 10 mg/kg plus single dose MQ at 25 mg/kg on day 2. All treatments were given under supervision. In total, 146+151=297 patients (aged 0.5-88 years, initial parasitaemia 2,111 – 4,687/µL) were enrolled in study 1 and 169+179=348 patients (aged 0.4-58 years, initial parasitaemia 4,196 – 7,480/µL) to study 2. Among those who completed 28-day follow-up, the PCR-uncorrected cure rates of MQ monotherapy in study 1 and study 2 were (115-22)/115=80.9% and (101-39+53-9)/(101+53)=68.83% respectively. Day-28 efficacy of single-dose ASMQ (study 1) was (124-21)/124=83.06% and efficacy of 3-day ASMQ (study 2) was (96-2+57-1)/(96+57)=98.04%.

**Thailand 1992-2002**. (Ric N. Price et al. 2004) recruited 1302 and 3322 *P. falciparum*-infected individuals in a Karen community in northwestern border of Thailand to MQ (total 25 mg/kg) and ASMQ (MQ 25 mg/kg + 3 days of AS at 4 mg/kg) treatment arm, respectively. Mean age of the patients in the two arms were 14 and 12 years, geometric mean of initial parasitaemia were 3,383/µL and 5,953/µL. The overall day-28 PCR-corrected efficacy was 802/1068=75% for MQ monotherapy and 2686/2886=93% for ASMQ combination. To investigate the effect of *pfmdr1* CNV on treatment outcomes, the authors genotyped 160 and 180 samples from each of the groups. According to figure 3 in this paper, day-28 efficacy of MQ on genotypes with a single copy of pfmdr1 was estimated to be around 92% and that on genotypes with multiple copies of pfmdr1 was 42-62%. With ASMQ, the estimates for day-28 efficacy were 95% and 65-87% for single and multiple copies of pfmdr1, respectively.

In the simulation, we set EC50 of MQ on genotypes carrying single and multiple copies of *pdmdr1* to be 0.45 and 1.1, respectively, to obtain day-28 efficacy of 3-day MQ monotherapy on MQ-sensitive genotypes (single copy of *pfmdr1*) at 94.5% and efficacy on MQ-resistant genotypes (multiple copies of *pfmdr1*) at 46.1% (tables on page 26-27) which resemble the reported outcomes in the trials above. Given the EC50 of AS presented in section 2 , we were able to approximate efficacies of ASMQ combination on 64 genotypes. Specifically, our estimate for efficacy of 3-day ASMQ on genotypes carrying *kelch13*-C580 + single copy *pfmdr1* is 98.3%, efficacy on *kelch13*-C580 + multiple copies *pfmdr1* is 84.6%, efficacy on *kelch13*-580Y + single copy *pfmdr1* is 96.2%, and efficacy on *kelch13*-580Y + multiple copies *pfmdr1* is 60.7% (tables on page 25-26).

| Genotype | AS | LM | AQ | PPQ | MQ | CQ | AL | ASAQ | DHAPPQ | ASMQ |
|---|---|---|---|---|---|---|---|---|---|---|
| KNY--C1 | 0.689 | 0.719 | | 0.899 | 0.945 | | 0.915 | | 0.972 | 0.983 |
| KNY--C2 | 0.689 | 0.719 | | 0.213 | 0.945 | | 0.915 | | 0.768 | 0.983 |
| KNY--Y1 | 0.241 | 0.719 | | 0.899 | 0.945 | | 0.795 | | 0.929 | 0.962 |
| KNY--Y2 | 0.241 | 0.719 | | 0.213 | 0.945 | | 0.795 | | 0.415 | 0.962 |
| KYY--C1 | 0.689 | 0.828 | | 0.899 | 0.945 | | 0.953 | | 0.972 | 0.983 |
| KYY--C2 | 0.689 | 0.828 | | 0.213 | 0.945 | | 0.953 | | 0.768 | 0.983 |
| KYY--Y1 | 0.241 | 0.828 | | 0.899 | 0.945 | | 0.878 | | 0.929 | 0.962 |
| KYY--Y2 | 0.241 | 0.828 | | 0.213 | 0.945 | | 0.878 | | 0.415 | 0.962 |
| KNF--C1 | 0.689 | 0.627 | | 0.899 | 0.945 | | 0.890 | | 0.972 | 0.983 |
| KNF--C2 | 0.689 | 0.627 | | 0.213 | 0.945 | | 0.890 | | 0.768 | 0.983 |
| KNF--Y1 | 0.241 | 0.627 | | 0.899 | 0.945 | | 0.723 | | 0.929 | 0.962 |
| KNF--Y2 | 0.241 | 0.627 | | 0.213 | 0.945 | | 0.723 | | 0.415 | 0.962 |
| KYF--C1 | 0.689 | 0.719 | | 0.899 | 0.945 | | 0.915 | | 0.972 | 0.983 |
| KYF--C2 | 0.689 | 0.719 | | 0.213 | 0.945 | | 0.915 | | 0.768 | 0.983 |
| KYF--Y1 | 0.241 | 0.719 | | 0.899 | 0.945 | | 0.795 | | 0.929 | 0.962 |
| KYF--Y2 | 0.241 | 0.719 | | 0.213 | 0.945 | | 0.795 | | 0.415 | 0.962 |
| KNYNYC1 | 0.689 | 0.523 | | 0.899 | | | 0.859 | | 0.972 | |
| KNYNYC2 | 0.689 | 0.523 | | 0.213 | | | 0.859 | | 0.768 | |
| KNYNYY1 | 0.241 | 0.523 | | 0.899 | | | 0.646 | | 0.929 | |
| KNYNYY2 | 0.241 | 0.523 | | 0.213 | | | 0.646 | | 0.415 | |
| KYYYYC1 | 0.689 | 0.662 | | 0.899 | | | 0.897 | | 0.972 | |
| KYYYYC2 | 0.689 | 0.662 | | 0.213 | | | 0.897 | | 0.768 | |
| KYYYYY1 | 0.241 | 0.662 | | 0.899 | | | 0.752 | | 0.929 | |
| KYYYYY2 | 0.241 | 0.662 | | 0.213 | | | 0.752 | | 0.415 | |
| KNFNFC1 | 0.689 | 0.422 | | 0.899 | | | 0.830 | | 0.972 | |
| KNFNFC2 | 0.689 | 0.422 | | 0.213 | | | 0.830 | | 0.768 | |
| KNFNFY1 | 0.241 | 0.422 | | 0.899 | | | 0.570 | | 0.929 | |
| KNFNFY2 | 0.241 | 0.422 | | 0.213 | | | 0.570 | | 0.415 | |
| KYFYFC1 | 0.689 | 0.523 | | 0.899 | | | 0.859 | | 0.972 | |
| KYFYFC2 | 0.689 | 0.523 | | 0.213 | | | 0.859 | | 0.768 | |
| KYFYFY1 | 0.241 | 0.523 | | 0.899 | | | 0.646 | | 0.929 | |
| KYFYFY2 | 0.241 | 0.523 | | 0.213 | | | 0.646 | | 0.415 | |
| TNY--C1 | 0.689 | 0.770 | | 0.899 | 0.945 | | 0.929 | | 0.972 | 0.983 |
| TNY--C2 | 0.689 | 0.770 | | 0.213 | 0.945 | | 0.929 | | 0.768 | 0.983 |
| TNY--Y1 | 0.241 | 0.770 | | 0.899 | 0.945 | | 0.829 | | 0.929 | 0.962 |
| TNY--Y2 | 0.241 | 0.770 | | 0.213 | 0.945 | | 0.829 | | 0.415 | 0.962 |
| TYY--C1 | 0.689 | 0.870 | | 0.899 | 0.945 | | 0.965 | | 0.972 | 0.983 |
| TYY--C2 | 0.689 | 0.870 | | 0.213 | 0.945 | | 0.965 | | 0.768 | 0.983 |
| TYY--Y1 | 0.241 | 0.870 | | 0.899 | 0.945 | | 0.908 | | 0.929 | 0.962 |
| TYY--Y2 | 0.241 | 0.870 | | 0.213 | 0.945 | | 0.908 | | 0.415 | 0.962 |
| TNF--C1 | 0.689 | 0.676 | | 0.899 | 0.945 | | 0.908 | | 0.972 | 0.983 |
| TNF--C2 | 0.689 | 0.676 | | 0.213 | 0.945 | | 0.908 | | 0.768 | 0.983 |
| TNF--Y1 | 0.241 | 0.676 | | 0.899 | 0.945 | | 0.753 | | 0.929 | 0.962 |
| TNF--Y2 | 0.241 | 0.676 | | 0.213 | 0.945 | | 0.753 | | 0.415 | 0.962 |
| TYF--C1 | 0.689 | 0.770 | | 0.899 | 0.945 | | 0.929 | | 0.972 | 0.983 |
| TYF--C2 | 0.689 | 0.770 | | 0.213 | 0.945 | | 0.929 | | 0.768 | 0.983 |
| TYF--Y1 | 0.241 | 0.770 | | 0.899 | 0.945 | | 0.829 | | 0.929 | 0.962 |
| TYF--Y2 | 0.241 | 0.770 | | 0.213 | 0.945 | | 0.829 | | 0.415 | 0.962 |
| TNYNYC1 | 0.689 | 0.575 | | 0.899 | | | 0.869 | | 0.972 | |
| TNYNYC2 | 0.689 | 0.575 | | 0.213 | | | 0.869 | | 0.768 | |
| TNYNYY1 | 0.241 | 0.575 | | 0.899 | | | 0.684 | | 0.929 | |
| TNYNYY2 | 0.241 | 0.575 | | 0.213 | | | 0.684 | | 0.415 | |
| TYYYYC1 | 0.689 | 0.719 | | 0.899 | | | 0.915 | | 0.972 | |
| TYYYYC2 | 0.689 | 0.719 | | 0.213 | | | 0.915 | | 0.768 | |
| TYYYYY1 | 0.241 | 0.719 | | 0.899 | | | 0.795 | | 0.929 | |
| TYYYYY2 | 0.241 | 0.719 | | 0.213 | | | 0.795 | | 0.415 | |
| TNFNFC1 | 0.689 | 0.474 | | 0.899 | | | 0.844 | | 0.972 | |
| TNFNFC2 | 0.689 | 0.474 | | 0.213 | | | 0.844 | | 0.768 | |
| TNFNFY1 | 0.241 | 0.474 | | 0.899 | | | 0.604 | | 0.929 | |
| TNFNFY2 | 0.241 | 0.474 | | 0.213 | | | 0.604 | | 0.415 | |
| TYFYFC1 | 0.689 | 0.575 | | 0.899 | | | 0.869 | | 0.972 | |
| TYFYFC2 | 0.689 | 0.575 | | 0.213 | | | 0.869 | | 0.768 | |
| TYFYFY1 | 0.241 | 0.575 | | 0.899 | | | 0.684 | | 0.929 | |
| TYFYFY2 | 0.241 | 0.575 | | 0.213 | | | 0.684 | | 0.415 | |

| Genotype | AS | LM | AQ | PPQ | MQ | CQ | AL | ASAQ | DHAPPQ | ASMQ |
|---|---|---|---|---|---|---|---|---|---|---|
| KNY--C1 | 0.689 | 0.719 | | 0.899 | 0.945 | | 0.915 | | 0.972 | 0.983 |
| KNY--C2 | 0.689 | 0.719 | | 0.213 | 0.945 | | 0.915 | | 0.768 | 0.983 |
| KNY--Y1 | 0.241 | 0.719 | | 0.899 | 0.945 | | 0.795 | | 0.929 | 0.962 |
| KNY--Y2 | 0.241 | 0.719 | | 0.213 | 0.945 | | 0.795 | | 0.415 | 0.962 |
| KYY--C1 | 0.689 | 0.828 | | 0.899 | 0.945 | | 0.953 | | 0.972 | 0.983 |
| KYY--C2 | 0.689 | 0.828 | | 0.213 | 0.945 | | 0.953 | | 0.768 | 0.983 |
| KYY--Y1 | 0.241 | 0.828 | | 0.899 | 0.945 | | 0.878 | | 0.929 | 0.962 |
| KYY--Y2 | 0.241 | 0.828 | | 0.213 | 0.945 | | 0.878 | | 0.415 | 0.962 |
| KNF--C1 | 0.689 | 0.627 | | 0.899 | 0.945 | | 0.890 | | 0.972 | 0.983 |
| KNF--C2 | 0.689 | 0.627 | | 0.213 | 0.945 | | 0.890 | | 0.768 | 0.983 |
| KNF--Y1 | 0.241 | 0.627 | | 0.899 | 0.945 | | 0.723 | | 0.929 | 0.962 |
| KNF--Y2 | 0.241 | 0.627 | | 0.213 | 0.945 | | 0.723 | | 0.415 | 0.962 |
| KYF--C1 | 0.689 | 0.719 | | 0.899 | 0.945 | | 0.915 | | 0.972 | 0.983 |
| KYF--C2 | 0.689 | 0.719 | | 0.213 | 0.945 | | 0.915 | | 0.768 | 0.983 |
| KYF--Y1 | 0.241 | 0.719 | | 0.899 | 0.945 | | 0.795 | | 0.929 | 0.962 |
| KYF--Y2 | 0.241 | 0.719 | | 0.213 | 0.945 | | 0.795 | | 0.415 | 0.962 |
| KNYNYC1 | 0.689 | 0.523 | | 0.899 | 0.461 | | 0.859 | | 0.972 | 0.846 |
| KNYNYC2 | 0.689 | 0.523 | | 0.213 | 0.461 | | 0.859 | | 0.768 | 0.846 |
| KNYNYY1 | 0.241 | 0.523 | | 0.899 | 0.461 | | 0.646 | | 0.929 | 0.607 |
| KNYNYY2 | 0.241 | 0.523 | | 0.213 | 0.461 | | 0.646 | | 0.415 | 0.607 |
| KYYYYC1 | 0.689 | 0.662 | | 0.899 | 0.461 | | 0.897 | | 0.972 | 0.846 |
| KYYYYC2 | 0.689 | 0.662 | | 0.213 | 0.461 | | 0.897 | | 0.768 | 0.846 |
| KYYYYY1 | 0.241 | 0.662 | | 0.899 | 0.461 | | 0.752 | | 0.929 | 0.607 |
| KYYYYY2 | 0.241 | 0.662 | | 0.213 | 0.461 | | 0.752 | | 0.415 | 0.607 |
| KNFNFC1 | 0.689 | 0.422 | | 0.899 | 0.461 | | 0.830 | | 0.972 | 0.846 |
| KNFNFC2 | 0.689 | 0.422 | | 0.213 | 0.461 | | 0.830 | | 0.768 | 0.846 |
| KNFNFY1 | 0.241 | 0.422 | | 0.899 | 0.461 | | 0.570 | | 0.929 | 0.607 |
| KNFNFY2 | 0.241 | 0.422 | | 0.213 | 0.461 | | 0.570 | | 0.415 | 0.607 |
| KYFYFC1 | 0.689 | 0.523 | | 0.899 | 0.461 | | 0.859 | | 0.972 | 0.846 |
| KYFYFC2 | 0.689 | 0.523 | | 0.213 | 0.461 | | 0.859 | | 0.768 | 0.846 |
| KYFYFY1 | 0.241 | 0.523 | | 0.899 | 0.461 | | 0.646 | | 0.929 | 0.607 |
| KYFYFY2 | 0.241 | 0.523 | | 0.213 | 0.461 | | 0.646 | | 0.415 | 0.607 |
| TNY--C1 | 0.689 | 0.770 | | 0.899 | 0.945 | | 0.929 | | 0.972 | 0.983 |
| TNY--C2 | 0.689 | 0.770 | | 0.213 | 0.945 | | 0.929 | | 0.768 | 0.983 |
| TNY--Y1 | 0.241 | 0.770 | | 0.899 | 0.945 | | 0.829 | | 0.929 | 0.962 |
| TNY--Y2 | 0.241 | 0.770 | | 0.213 | 0.945 | | 0.829 | | 0.415 | 0.962 |
| TYY--C1 | 0.689 | 0.870 | | 0.899 | 0.945 | | 0.965 | | 0.972 | 0.983 |
| TYY--C2 | 0.689 | 0.870 | | 0.213 | 0.945 | | 0.965 | | 0.768 | 0.983 |
| TYY--Y1 | 0.241 | 0.870 | | 0.899 | 0.945 | | 0.908 | | 0.929 | 0.962 |
| TYY--Y2 | 0.241 | 0.870 | | 0.213 | 0.945 | | 0.908 | | 0.415 | 0.962 |
| TNF--C1 | 0.689 | 0.676 | | 0.899 | 0.945 | | 0.908 | | 0.972 | 0.983 |
| TNF--C2 | 0.689 | 0.676 | | 0.213 | 0.945 | | 0.908 | | 0.768 | 0.983 |
| TNF--Y1 | 0.241 | 0.676 | | 0.899 | 0.945 | | 0.753 | | 0.929 | 0.962 |
| TNF--Y2 | 0.241 | 0.676 | | 0.213 | 0.945 | | 0.753 | | 0.415 | 0.962 |
| TYF--C1 | 0.689 | 0.770 | | 0.899 | 0.945 | | 0.929 | | 0.972 | 0.983 |
| TYF--C2 | 0.689 | 0.770 | | 0.213 | 0.945 | | 0.929 | | 0.768 | 0.983 |
| TYF--Y1 | 0.241 | 0.770 | | 0.899 | 0.945 | | 0.829 | | 0.929 | 0.962 |
| TYF--Y2 | 0.241 | 0.770 | | 0.213 | 0.945 | | 0.829 | | 0.415 | 0.962 |
| TNYNYC1 | 0.689 | 0.575 | | 0.899 | 0.461 | | 0.869 | | 0.972 | 0.846 |
| TNYNYC2 | 0.689 | 0.575 | | 0.213 | 0.461 | | 0.869 | | 0.768 | 0.846 |
| TNYNYY1 | 0.241 | 0.575 | | 0.899 | 0.461 | | 0.684 | | 0.929 | 0.607 |
| TNYNYY2 | 0.241 | 0.575 | | 0.213 | 0.461 | | 0.684 | | 0.415 | 0.607 |
| TYYYYC1 | 0.689 | 0.719 | | 0.899 | 0.461 | | 0.915 | | 0.972 | 0.846 |
| TYYYYC2 | 0.689 | 0.719 | | 0.213 | 0.461 | | 0.915 | | 0.768 | 0.846 |
| TYYYYY1 | 0.241 | 0.719 | | 0.899 | 0.461 | | 0.795 | | 0.929 | 0.607 |
| TYYYYY2 | 0.241 | 0.719 | | 0.213 | 0.461 | | 0.795 | | 0.415 | 0.607 |
| TNFNFC1 | 0.689 | 0.474 | | 0.899 | 0.461 | | 0.844 | | 0.972 | 0.846 |
| TNFNFC2 | 0.689 | 0.474 | | 0.213 | 0.461 | | 0.844 | | 0.768 | 0.846 |
| TNFNFY1 | 0.241 | 0.474 | | 0.899 | 0.461 | | 0.604 | | 0.929 | 0.607 |
| TNFNFY2 | 0.241 | 0.474 | | 0.213 | 0.461 | | 0.604 | | 0.415 | 0.607 |
| TYFYFC1 | 0.689 | 0.575 | | 0.899 | 0.461 | | 0.869 | | 0.972 | 0.846 |
| TYFYFC2 | 0.689 | 0.575 | | 0.213 | 0.461 | | 0.869 | | 0.768 | 0.846 |
| TYFYFY1 | 0.241 | 0.575 | | 0.899 | 0.461 | | 0.684 | | 0.929 | 0.607 |
| TYFYFY2 | 0.241 | 0.575 | | 0.213 | 0.461 | | 0.684 | | 0.415 | 0.607 |

## 4.2 CQ

**Malawi 2005**. (Laufer et al. 2006). 80 children aged 0.4-4.8 years, geometric mean parasitaemia of 19,379 parasites/µL received total dose of 25 mg/kg (10 mg/kg over the first two days, then 5 mg/kg on the third day) mono chloroquine sulfate over 3 days. 87 children aged 0.7-5.1 years, geometric mean parasitaemia of 18,856/µL received single dose of Fansidar (1.25 mg/kg sulfadoxine + 25 mg/kg pyrimerthamine). Drugs were administered under direct supervision. Day-28 ACPR rate of chloroquine was 99% and that of SP was 21%. All samples in the study had *pfcrt*-K76 allele.

**Mali 1997**. (Djimdé et al. 2001). 514 patients (median age 10 years, median initial parasitaemia 12,800/µL) were enrolled in a 3-day CQ at 25 mg/kg trial, 469 of whom finished 14-day follow up. Treatment outcomes were assessed using (World Health Organization 1996) classification. Overall, day-14 cure rate of CQ was 86%; Table 2 in this paper shown day-14 efficacy (PCR-uncorrected) *pfcrt*-76T and *pfmdr1*-86Y to be 22/(22+43)=33.8%

**Ghana 2000**. In 2000, (Ehrhardt et al. 2002) evaluated the efficacy of CQ in part of northern Ghana by recruiting *P. falciparum*-infected children who were under 5 years old and treating them with 3-day CQ (total dose 25 mg/kg) under supervision. The study included children with residual CQ from previous antimalarial ingestion and those with malnutrition. Treatment outcomes were classified as early treatment failure (ETF), late treatment failure (LTF), and adequate clinical response (ACR) according to WHO recommendations (World Health Organization 1996). In total, 225 children with mean age of 36 months old, mean weight of 11.6 kg, and geometric mean of initial parasitaemia of 28,184/µL finished 2-week follow-up; the overall day-14 PCR-uncorrected efficacy of CQ was 160/225=71.1%. (Mockenhaupt et al. 2005) analyzed samples from (Ehrhardt et al. 2002) trial to determine the role of *pfcrt* and *pfmdr1* genes in CQ treatment outcomes and found that recrudescence was associated with *pfcrt*-K76T and *pfmdr1*-N86Y mutations. After stratifying the samples by genotype, (Mockenhaupt et al. 2005) observed that day-14 ACPR rate (as defined in (World Health Organization 2003)) of CQ on *pfcrt*-K76 + *pfmdr1*-N86 was (32-11)/32=65.6%, ACPR on *pfcrt*-K76 + *pfmdr1*-86Y (10-4)/10=60%, ACPR on *pfcrt*-76T + *pfmdr1*-N86 was (65-34)/65=47.7%, and ACPR on *pfcrt*-76T + *pfmdr1*-86Y was (117-82)/117=29.9%. PCR correction for new infection was done in 58 of 131 treatment failures. If PCR correction had been done in all failure cases, the genotype-stratified efficacies could have been higher.

**Nigeria around 2003**, (T. C. Happi et al. 2003) recruited 60 *P. falciparum*-infected children aged 1-12 years with geometric mean of initial parasitaemia of 5,897/µL and treated them with 3-day CQ at total dose of 25 mg/kg. The overall cure rate (according to (World Health Organization 1973; 1996)) at day 28 was 31/60=51.7%. 15 samples from each of treatment success and treatment failure group were randomly selected to be further analyzed via genotyping. Samples from the treatment failure group were confirmed to be recrudescence. According to Table 3 in the paper, day-28 efficacy of 3-day CQ on *pfcrt*-K76 + *pfmdr1*-N86 was 3/3=100%, efficacy on *pfcrt*-K76 + *pfmdr1*-86Y was 3/4=75%, efficacy on *pfcrt*-76T + *pfmdr1*-N86 was 3/5=60%, and efficacy on *pfcrt*-76T + *pfmdr1*-86Y was 6/17=32.3%.

**India 2008-2009**. To study the role of genetic polymorphisms in CQ treatment failures, (Das et al. 2014) enrolled 126 patients with *P. falciparum*-confirmed monoinfection in eastern India from 2008 to 2009 and treated them with 3-day CQ monotherapy at a total dose of 25 mg/kg under supervision. Baseline characteristics of the patients were not presented. According to Table 3 in the paper, day-28 PCR-corrected efficacy of CQ on *pfcrt*-K76 + *pfmdr1*-N86 was 5/5=100%, efficacy on *pfcrt*-K76 + *pfmdr1*-86Y was (5+1)/(37+11) = 12.5%, efficacy on *pfcrt*-76T + *pfmdr1*-N86 was (17+5+2+10+2)/(21+5+2+12+4) = 81.8%, and efficacy on *pfcrt*-76T + *pfmdr1*-86Y was (9+0+11)/(14+3+12) = 69.0%.

By setting EC50 of chloroquine on *pfcrt*-K76 + *pfmdr1*-N86 to 0.72, we obtained day-28 efficacy of 3-day CQ monotherapy at 81.0%. With EC50=0.9 on *pfcrt*-K76 + *pfmdr1*-86Y, we obtained day-28 efficacy of CQ monotherapy at 63.9%. EC50=1.19 for CQ on *pfcrt*-76T + *pfmdr1*-N86 yielded an efficacy of 33.7%. When a patient is infected with parasites carrying mutants on both loci *pfcrt*-76 and *pfmdr1*-86, our estimate for day-28 efficacy of CQ monotherapy is 19.5%, which can be achieved by setting EC50 of CQ on pfcrt-76T + pfmdr1-86Y to 1.35.

| Genotype | AS | LM | AQ | PPQ | MQ | CQ | AL | ASAQ | DHAPPQ | ASMQ |
|---|---|---|---|---|---|---|---|---|---|---|
| KNY--C1 | 0.689 | 0.719 | | 0.899 | 0.945 | 0.810 | 0.915 | | 0.972 | 0.983 |
| KNY--C2 | 0.689 | 0.719 | | 0.213 | 0.945 | 0.810 | 0.915 | | 0.768 | 0.983 |
| KNY--Y1 | 0.241 | 0.719 | | 0.899 | 0.945 | 0.810 | 0.795 | | 0.929 | 0.962 |
| KNY--Y2 | 0.241 | 0.719 | | 0.213 | 0.945 | 0.810 | 0.795 | | 0.415 | 0.962 |
| KYY--C1 | 0.689 | 0.828 | | 0.899 | 0.945 | 0.639 | 0.953 | | 0.972 | 0.983 |
| KYY--C2 | 0.689 | 0.828 | | 0.213 | 0.945 | 0.639 | 0.953 | | 0.768 | 0.983 |
| KYY--Y1 | 0.241 | 0.828 | | 0.899 | 0.945 | 0.639 | 0.878 | | 0.929 | 0.962 |
| KYY--Y2 | 0.241 | 0.828 | | 0.213 | 0.945 | 0.639 | 0.878 | | 0.415 | 0.962 |
| KNF--C1 | 0.689 | 0.627 | | 0.899 | 0.945 | 0.810 | 0.890 | | 0.972 | 0.983 |
| KNF--C2 | 0.689 | 0.627 | | 0.213 | 0.945 | 0.810 | 0.890 | | 0.768 | 0.983 |
| KNF--Y1 | 0.241 | 0.627 | | 0.899 | 0.945 | 0.810 | 0.723 | | 0.929 | 0.962 |
| KNF--Y2 | 0.241 | 0.627 | | 0.213 | 0.945 | 0.810 | 0.723 | | 0.415 | 0.962 |
| KYF--C1 | 0.689 | 0.719 | | 0.899 | 0.945 | 0.639 | 0.915 | | 0.972 | 0.983 |
| KYF--C2 | 0.689 | 0.719 | | 0.213 | 0.945 | 0.639 | 0.915 | | 0.768 | 0.983 |
| KYF--Y1 | 0.241 | 0.719 | | 0.899 | 0.945 | 0.639 | 0.795 | | 0.929 | 0.962 |
| KYF--Y2 | 0.241 | 0.719 | | 0.213 | 0.945 | 0.639 | 0.795 | | 0.415 | 0.962 |
| KNYNYC1 | 0.689 | 0.523 | | 0.899 | 0.461 | 0.810 | 0.859 | | 0.972 | 0.846 |
| KNYNYC2 | 0.689 | 0.523 | | 0.213 | 0.461 | 0.810 | 0.859 | | 0.768 | 0.846 |
| KNYNYY1 | 0.241 | 0.523 | | 0.899 | 0.461 | 0.810 | 0.646 | | 0.929 | 0.607 |
| KNYNYY2 | 0.241 | 0.523 | | 0.213 | 0.461 | 0.810 | 0.646 | | 0.415 | 0.607 |
| KYYYYC1 | 0.689 | 0.662 | | 0.899 | 0.461 | 0.639 | 0.897 | | 0.972 | 0.846 |
| KYYYYC2 | 0.689 | 0.662 | | 0.213 | 0.461 | 0.639 | 0.897 | | 0.768 | 0.846 |
| KYYYYY1 | 0.241 | 0.662 | | 0.899 | 0.461 | 0.639 | 0.752 | | 0.929 | 0.607 |
| KYYYYY2 | 0.241 | 0.662 | | 0.213 | 0.461 | 0.639 | 0.752 | | 0.415 | 0.607 |
| KNFNFC1 | 0.689 | 0.422 | | 0.899 | 0.461 | 0.810 | 0.830 | | 0.972 | 0.846 |
| KNFNFC2 | 0.689 | 0.422 | | 0.213 | 0.461 | 0.810 | 0.830 | | 0.768 | 0.846 |
| KNFNFY1 | 0.241 | 0.422 | | 0.899 | 0.461 | 0.810 | 0.570 | | 0.929 | 0.607 |
| KNFNFY2 | 0.241 | 0.422 | | 0.213 | 0.461 | 0.810 | 0.570 | | 0.415 | 0.607 |
| KYFYFC1 | 0.689 | 0.523 | | 0.899 | 0.461 | 0.639 | 0.859 | | 0.972 | 0.846 |
| KYFYFC2 | 0.689 | 0.523 | | 0.213 | 0.461 | 0.639 | 0.859 | | 0.768 | 0.846 |
| KYFYFY1 | 0.241 | 0.523 | | 0.899 | 0.461 | 0.639 | 0.646 | | 0.929 | 0.607 |
| KYFYFY2 | 0.241 | 0.523 | | 0.213 | 0.461 | 0.639 | 0.646 | | 0.415 | 0.607 |
| TNY--C1 | 0.689 | 0.770 | | 0.899 | 0.945 | 0.337 | 0.929 | | 0.972 | 0.983 |
| TNY--C2 | 0.689 | 0.770 | | 0.213 | 0.945 | 0.337 | 0.929 | | 0.768 | 0.983 |
| TNY--Y1 | 0.241 | 0.770 | | 0.899 | 0.945 | 0.337 | 0.829 | | 0.929 | 0.962 |
| TNY--Y2 | 0.241 | 0.770 | | 0.213 | 0.945 | 0.337 | 0.829 | | 0.415 | 0.962 |
| TYY--C1 | 0.689 | 0.870 | | 0.899 | 0.945 | 0.195 | 0.965 | | 0.972 | 0.983 |
| TYY--C2 | 0.689 | 0.870 | | 0.213 | 0.945 | 0.195 | 0.965 | | 0.768 | 0.983 |
| TYY--Y1 | 0.241 | 0.870 | | 0.899 | 0.945 | 0.195 | 0.908 | | 0.929 | 0.962 |
| TYY--Y2 | 0.241 | 0.870 | | 0.213 | 0.945 | 0.195 | 0.908 | | 0.415 | 0.962 |
| TNF--C1 | 0.689 | 0.676 | | 0.899 | 0.945 | 0.337 | 0.908 | | 0.972 | 0.983 |
| TNF--C2 | 0.689 | 0.676 | | 0.213 | 0.945 | 0.337 | 0.908 | | 0.768 | 0.983 |
| TNF--Y1 | 0.241 | 0.676 | | 0.899 | 0.945 | 0.337 | 0.753 | | 0.929 | 0.962 |
| TNF--Y2 | 0.241 | 0.676 | | 0.213 | 0.945 | 0.337 | 0.753 | | 0.415 | 0.962 |
| TYF--C1 | 0.689 | 0.770 | | 0.899 | 0.945 | 0.195 | 0.929 | | 0.972 | 0.983 |
| TYF--C2 | 0.689 | 0.770 | | 0.213 | 0.945 | 0.195 | 0.929 | | 0.768 | 0.983 |
| TYF--Y1 | 0.241 | 0.770 | | 0.899 | 0.945 | 0.195 | 0.829 | | 0.929 | 0.962 |
| TYF--Y2 | 0.241 | 0.770 | | 0.213 | 0.945 | 0.195 | 0.829 | | 0.415 | 0.962 |
| TNYNYC1 | 0.689 | 0.575 | | 0.899 | 0.461 | 0.337 | 0.869 | | 0.972 | 0.846 |
| TNYNYC2 | 0.689 | 0.575 | | 0.213 | 0.461 | 0.337 | 0.869 | | 0.768 | 0.846 |
| TNYNYY1 | 0.241 | 0.575 | | 0.899 | 0.461 | 0.337 | 0.684 | | 0.929 | 0.607 |
| TNYNYY2 | 0.241 | 0.575 | | 0.213 | 0.461 | 0.337 | 0.684 | | 0.415 | 0.607 |
| TYYYYC1 | 0.689 | 0.719 | | 0.899 | 0.461 | 0.195 | 0.915 | | 0.972 | 0.846 |
| TYYYYC2 | 0.689 | 0.719 | | 0.213 | 0.461 | 0.195 | 0.915 | | 0.768 | 0.846 |
| TYYYYY1 | 0.241 | 0.719 | | 0.899 | 0.461 | 0.195 | 0.795 | | 0.929 | 0.607 |
| TYYYYY2 | 0.241 | 0.719 | | 0.213 | 0.461 | 0.195 | 0.795 | | 0.415 | 0.607 |
| TNFNFC1 | 0.689 | 0.474 | | 0.899 | 0.461 | 0.337 | 0.844 | | 0.972 | 0.846 |
| TNFNFC2 | 0.689 | 0.474 | | 0.213 | 0.461 | 0.337 | 0.844 | | 0.768 | 0.846 |
| TNFNFY1 | 0.241 | 0.474 | | 0.899 | 0.461 | 0.337 | 0.604 | | 0.929 | 0.607 |
| TNFNFY2 | 0.241 | 0.474 | | 0.213 | 0.461 | 0.337 | 0.604 | | 0.415 | 0.607 |
| TYFYFC1 | 0.689 | 0.575 | | 0.899 | 0.461 | 0.195 | 0.869 | | 0.972 | 0.846 |
| TYFYFC2 | 0.689 | 0.575 | | 0.213 | 0.461 | 0.195 | 0.869 | | 0.768 | 0.846 |
| TYFYFY1 | 0.241 | 0.575 | | 0.899 | 0.461 | 0.195 | 0.684 | | 0.929 | 0.607 |
| TYFYFY2 | 0.241 | 0.575 | | 0.213 | 0.461 | 0.195 | 0.684 | | 0.415 | 0.607 |

### 5 AQ, ASAQ

**Burkina Faso**. Original efficacy trial: (Zongo et al. 2005); molecular markers association study: (Dokomajilar et al. 2006). Data collection year was not mentioned explicitly in the two papers. The efficacy trial compared three treatments namely single dose of 25 mg/kg sulfadoxine + 1.25 mg/kg pyrimethamine (SP), 3-day of total 25 mg/kg amodiaquine (AQ), and SP+AQ (similar dosing regimen as in monotherapy arms). There were 264, 280, and 285 patients aged 0.5-52 years who completed 28-day follow-up in the three arms respectively. In the molecular markers association study, (Dokomajilar et al. 2006) randomly selected 200 samples, 80 from SP arm and 110 from AQ, to evaluate the effect of polymorphisms on *dhfr*, *dhps*, *pfcrt*, and *pfmdr1* gene on treatment outcomes. Based on Table 1 in (Dokomajilar et al. 2006), we estimated genotype-specific efficacies of AQ as follows:

- Among samples with *pfcrt* markers, 42 had K76 allele and 68 had 76T allele. Therefore, the sum of genotype-specific number of samples would be:
  - (pfcrt-K76 + *pfmdr1*-N86) and (*pfcrt*-K76 + *pfmdr1*-86Y) = 42.
  - (*pfcrt*-76T + *pfmdr1*-N86) and (*pfcrt*-76T + *pfmdr1*-86Y) = 68.
- Among samples with *pfmdr1* markers, 62 had N86 allele and 48 had 86Y allele. Therefore, the sum of genotype-specific number of samples would be:
  - (*pfcrt*-K76 + *pfmdr1*-N86) and (*pfcrt*-76T + *pfmdr1*-N86) = 62.
  - (*pfcrt*-K76 + *pfmdr1*-86Y) and (*pfcrt*-76T + *pfmdr1*-86Y) = 48.
- When *pfcrt* and *pfmdr1* were reported together, the number of samples carrying wild-type allele at any locus was 73. In other words:
  - (pfcrt-K76 + *pfmdr1*-N86) and (*pfcrt*-K76 + *pfmdr1*-86Y) and (p*fcrt*-76T + *pfmdr1*-N86) = 73.
- Hence, in total, we would have:
  - (*pfcrt*-76T + *pfmdr1*-N86) = 73 - 42 = 31
  - (*pfcrt*-K76 + *pfmdr1*-N86) = 62 - 31 = 31
  - (*pfcrt*-K76 + *pfmdr1*-86Y) = 42 - 31 = 11
  - (*pfcrt*-76T + *pfmdr1*-86Y) = 68 - 31 = 37
- Among recrudescent samples with *pfcrt* marker, 2 had K76 and 6 had 76T allele; thus:
  - recrudescence (*pfcrt*-K76 + *pfmdr1*-N86) and (*pfcrt*-K76 + *pfmdr1*-86Y) = 2.
  - recrudescence (*pfcrt*-76T + *pfmdr1*-N86) and (*pfcrt*-76T + *pfmdr1*-86Y) = 16.
- Among recrudescent samples with *pfmdr1* marker, 4 had N86 and 14 had 86Y allele; thus:
  - recrudescence (*pfcrt*-K76 + *pfmdr1*-N86) and (*pfcrt*-76T + *pfmdr1*-N86) = 4.
  - recrudescence (*pfcrt*-K76 + *pfmdr1*-86Y) + (*pfcrt*-76T + *pfmdr1*-86Y) = 14.
- Among recrudescent samples with *pfcrt* and *pfmdr1* markers, 14 had mixed or mutant at all alleles and 4 had wild-type at any alleles; thus:
  - recrudescence (*pfcrt*-76T + *pfmdr1*-86Y) = 14.
  - recrudescence (*pfcrt*-K76 & *pfmdr1*-N86) and (*pfcrt*-K76 + *pfmdr1*-86Y) and (*pfcrt*-76T + *pfmdr1*-N86) = 4.
- Hence, in recrudescent cases, we have:
  - (*pfcrt*-76T + *pfmdr1*-N86) = 4 - 2 = 2.
  - (*pfcrt*-K76 + *pfmdr1*-N86) = 4 - 2 = 2.
  - (*pfcrt*-K76 + *pfmdr1*-86Y) = 2 - 2 = 0.
  - (*pfcrt*-76T + *pfmdr1*-86Y) = 14.

According to the above numbers, PCR-corrected day-28 of AQ monotherapy on (*pfcrt*-K76 + *pfmdr1*-N86) was (31-2)/31=93.55%, on (*pfcrt*-K76 + *pfmdr1*-86Y) was (11-0)/11=100%, on (*pfcrt*-76T + *pfmdr1*-N86) was (31-2)/31=93.55%, and on (*pfcrt*-76T + *pfmdr1*-86Y) was (37-14)/37=62.16%.

**Burkina Faso**, year of trial is unclear. (Tinto et al. 2008) tested the effects of mutations at locus 76 on *pfcrt* gene and locus 86 on *pfmdr1* gene on clinical efficacy of AQ in children aged 6 months to 15 years. AQ was administered according to (World Health Organization 2003) recommendations, which ranged from 25 to 35 mg/kg. The paper, however, did not report in detail patients' characteristics such as weight and initial parasitaemia. According to Table 1 in the original publication, PCR-corrected efficacy of AQ on (*pfcrt*-K76 + *pfmdr1*-N86) was 56/(56+1)=98.2%, on (*pfcrt*-K76 + *pfmdr1*-86Y) was 13/13=100%, on (*pfcrt*-76T + *pfmdr1*-N86) was 65/(65+3)=95.6%, and on (*pfcrt*-76T + *pfmdr1*-86Y) was 35/(35+8)=81.4%.

**Burkina Faso 2005**. To evaluate the effect of the partner drug in ASAQ combination, which was officially adopted as first-line therapy in Burkina Faso in 2005, (Mandi et al. 2008) recruited 117 *P. falciparum*-confirmed children in northwestern Burkina Faso from September to November 2005 and treated them with 3-day AQ monotherapy at a total dose of 25 mg/kg (10 mg/kg on the first two days, 5 mg/kg on the third day) under supervision. The children were grouped into rural (n=62) and urban (n=55) area; mean age and mean initial parasitaemia (counted as trophozoites density) of the children in the two groups were 33.8 months, 26.4 months, 16,000/µl, and 18,000/µl, respectively. The overall day-28 PCR-corrected efficacy of AQ was 71/117=60.7%. To investigate the selection of *pfcrt* and *pfmdr1* after AQ treatment, (Danquah et al. 2010) genotyped samples from this trial for mutations at locus 76 on *pfcrt* and loci 86, 184, 1034, 1042, and 1246 on *pfmdr1*. In total, 109 and 111 samples were successfully determined for polymorphisms on *pfcrt* and *pfmdr1*, respectively. If we assume the two samples whose *pfcrt* genotyping failed carried *pfmdr1*-N86, we can estimate day-28 genotype-stratified efficacy of AQ based on Table 1 in (Danquah et al. 2010) as follows (mixed results are counted as mutants):

- Among samples with *pfcrt* markers, 52 had K76 allele and 57 had 76T allele. Therefore, the sum of genotype-specific number of samples would be:
  - (p*fcrt*-K76 + *pfmdr1*-N86) and (*pfcrt*-K76 + *pfmdr1*-86Y) = 52.
  - (p*fcrt*-76T + *pfmdr1*-N86) and (*pfcrt*-76T + *pfmdr1*-86Y) = 57.
- Among samples with *pfmdr1* markers, (76-2)=74 had N86 allele and 35 had 86Y allele. Therefore, the sum of genotype-specific number of samples would be:
  - (p*fcrt*-K76 + *pfmdr1*-N86) and (*pfcrt*-76T + *pfmdr1*-N86) = 74.
  - (p*fcrt*-K76 + *pfmdr1*-86Y) and (*pfcrt*-76T + *pfmdr1*-86Y) = 35.

- When *pfcrt* and *pfmdr1* were reported together, the number of samples carrying mutant allele at both loci was 25.; thus:
  - (p*fcrt*-76T + *pfmdr1*-86Y) = 25.

- Hence, in total, we would have:
  - (*pfcrt*-76T + *pfmdr1*-N86) = 57 – 25 = 32
  - (*pfcrt*-K76 + *pfmdr1*-N86) = 74 – 32 = 42
  - (*pfcrt*-K76 + *pfmdr1*-86Y) = 35 – 25 = 52 – 42 = 10
  - (*pfcrt*-76T + *pfmdr1*-86Y) = 25

- Among recrudescent samples with *pfcrt* marker, 0 had K76 and 32 had 76T allele; thus:
  - recrudescence (*pfcrt*-K76 + *pfmdr1*-N86) and (*pfcrt*-K76 + *pfmdr1*-86Y) = 17.
  - recrudescence (*pfcrt*-76T + *pfmdr1*-N86) and (*pfcrt*-76T + *pfmdr1*-86Y) = 15.

- Among recrudescent samples with *pfmdr1* marker, (22-1)=21 had N86 and 11 had 86Y allele; thus:
  - recrudescence (*pfcrt*-K76 + *pfmdr1*-N86) and (*pfcrt*-76T + *pfmdr1*-N86) = 21.
  - recrudescence (*pfcrt*-K76 + *pfmdr1*-86Y) + (*pfcrt*-76T + *pfmdr1*-86Y) = 11.

- Among recrudescent samples with *pfcrt* and *pfmdr1* markers, the number of samples carrying mutant allele at both loci was 20 ; thus:
  - recrudescence (*pfcrt*-76T + *pfmdr1*-86Y) = 7.
- Hence, in recrudescent cases, we have:
  - (*pfcrt*-76T + *pfmdr1*-N86) = 15 – 7 = 8.
  - (*pfcrt*-K76 + *pfmdr1*-N86) = 21 – 8 = 13.
  - (*pfcrt*-K76 + *pfmdr1*-86Y) = 11 – 7 = 17 – 13 = 4.
  - (*pfcrt*-76T + *pfmdr1*-86Y) = 7.

According to the above numbers, PCR-corrected day-28 of AQ monotherapy on (*pfcrt*-K76 + *pfmdr1*-N86) was (42-13)/42=69.1%, on (*pfcrt*-K76 + *pfmdr1*-86Y) was (10-4)/10=60%, on (*pfcrt*-76T + *pfmdr1*-N86) was (32-8)/32=75%, and on (*pfcrt*-76T + *pfmdr1*-86Y) was (25-7)/25=72%. Since the majority (87.4%) of samples in the trials had *pfmdr1*-184F at admission, we, for simplicity, use these efficacies to estimate AQ cure rates on genotypes carrying *pfmdr1*-184F.

**Nigeria 2005**. To investigate the effect of *pfcrt* and *pfmdr1* on susceptibility to AQ, (C. T. Happi et al. 2006) recruited and successfully followed 101 *P. falciparum*-infected children with mean age of 6 years old and geometric mean of initial parasitaemia of 23,033/μl (range: 2,070-180,390/μl) in Ibadan, Nigeria from April to November 2005. The patients were treated under supervision with AQ monotherapy at total dose of 30 mg/kg over three days. The overall PCR-corrected day-28 cure rate was 88/101=87.1%. We can infer genotype-stratified efficacies from Table 4 in the paper as follows (mixed results are counted as mutants):

- Among samples with *pfcrt* markers, 22 had K76 allele and (63+16)=79 had 76T allele. Therefore, the sum of genotype-specific number of samples would be:
  - (p*fcrt*-K76 + *pfmdr1*-N86) and (*pfcrt*-K76 + *pfmdr1*-86Y) = 22.
  - (p*fcrt*-76T + *pfmdr1*-N86) and (*pfcrt*-76T + *pfmdr1*-86Y) = 79.
- Among samples with *pfmdr1* markers, 44 had N86 allele and (29+28)=57 had 86Y allele. Therefore, the sum of genotype-specific number of samples would be:
  - (p*fcrt*-K76 + *pfmdr1*-N86) and (*pfcrt*-76T + *pfmdr1*-N86) = 44.
  - (p*fcrt*-K76 + *pfmdr1*-86Y) and (*pfcrt*-76T + *pfmdr1*-86Y) = 57.

- When *pfcrt* and *pfmdr1* were reported together, the number of samples carrying wild-type allele at both loci was 12 and the number of samples carrying mutant allele at both loci was 47.; thus:
  - (p*fcrt*-K76 + *pfmdr1*-N86) = 12.
  - (p*fcrt*-76T + *pfmdr1*-86Y) = 47.

- Hence, in total, we would have:
  - (*pfcrt*-K76 + *pfmdr1*-N86) = 12
  - (*pfcrt*-K76 + *pfmdr1*-86Y) = 22 – 12 = 57 – 47 = 10
  - (*pfcrt*-76T + *pfmdr1*-N86) = 79 – 47 = 44 – 12 = 32
  - (*pfcrt*-76T + *pfmdr1*-86Y) = 47

- Among recrudescent samples with *pfcrt* marker, 0 had K76 and (12+1)=13 had 76T allele; thus:
  - recrudescence (*pfcrt*-K76 + *pfmdr1*-N86) and (*pfcrt*-K76 + *pfmdr1*-86Y) = 0.

- recrudescence (*pfcrt*-76T + *pfmdr1*-N86) and (*pfcrt*-76T + *pfmdr1*-86Y) = 13.

- Among recrudescent samples with *pfmdr1* marker, 1 had N86 and (11+1)=12 had 86Y allele; thus:
  - recrudescence (*pfcrt*-K76 + *pfmdr1*-N86) and (*pfcrt*-76T + *pfmdr1*-N86) = 1.
  - recrudescence (*pfcrt*-K76 + *pfmdr1*-86Y) + (*pfcrt*-76T + *pfmdr1*-86Y) = 12.

- Among recrudescent samples with *pfcrt* and *pfmdr1* markers, tthe number of samples carrying wild-type allele at both loci was 0 and the number of samples carrying mutant allele at both loci was 12.; thus:
  - (p*fcrt*-K76 + *pfmdr1*-N86) = 0.
  - (*pfcrt*-76T + *pfmdr1*-86Y) = 12.
- Hence, in recrudescent cases, we have:
  - (*pfcrt*-K76 + *pfmdr1*-N86) = 0
  - (*pfcrt*-K76 + *pfmdr1*-86Y) = 0 – 0 = 12 -12 = 0
  - (*pfcrt*-76T + *pfmdr1*-N86) = 13 – 12 = 1 – 0 = 1
  - (*pfcrt*-76T + *pfmdr1*-86Y) = 12

According to the above numbers, PCR-corrected day-28 of AQ monotherapy on (*pfcrt*-K76 + *pfmdr1*-N86) was (12-0)/12=100%, on (*pfcrt*-K76 + *pfmdr1*-86Y) was (10-0)/10=100%, on (*pfcrt*-76T + *pfmdr1*-N86) was (32-1)/32=96.9%, and on (*pfcrt*-76T + *pfmdr1*-86Y) was (47-12)/42=83.3%.

**Uganda 2013-2014**. To compare the efficacy of AL and ASAQ, (Yeka et al. 2015) enrolled 602 *P.falciparum*-infected children under 5 years old in Northern, Central, and Western Uganda from 2013 to 2014 and treated them with 3-day standard weight-based of either AL (total dose of around 12 mg/kg artemether + 72 mg/kg lumefantrine) or ASAQ (total dose of around 3.7 mg/kg artesunate + 10 mg/kg amodiaquine). Mean age of the patients in both arms was from 2.4 to 3.0 years. Geometric mean of initial parasitaemia was lowest in the Western site (AL arm: 12,264/μl; ASAQ arm: 12,827/μl) and highest in the Central site (AL arm: 35,153/μl; ASAQ arm: 30,864/μl). Geometric mean of initial parasitaemia of patients in the Northern site was 21,616/μl and 22,614/μl in AL and ASAQ arm, respectively. In total, 594 patients completed 28-day follow-up. The overall PCR-corrected cure rates of AL and ASAQ for all three sites 100-2.5=97.5% and 100-0=100%, respectively. When looking at genetic polymorphisms, the authors found that most of the samples carried *pfcrt*-76T (>75%), *pfmdr1*-N86 (>90%), and *pfmdr1*-184F (>60%) at baseline (we consider mixed alleles as mutants). However, there was no information on genotype (e.g. *pfcrt*-K76 + *pfmdr1*-N86-Y184, *pfcrt*-K76 + *pfmdr1*-N86-184F, *pfcrt*-K76 + *pfmdr1*-86Y-Y184, etc.) distribution in the study.

We calibrated our model so that day-28 cure rates of AQ monotherapy reflect the selection pressure imposed by this antimalarial, results from the above therapeutic efficacy studies and that day-28 efficacies of ASAQ combination on artemisinin-sensitive genotypes (i.e. genotypes carrying *kelch13*-C580) should be at least 90% as observed in a meta-analysis of 43 ASAQ trials from 1992 to 2012 (WWARN AS-AQ Study Group 2015). By setting EC50 of AQ on KNF-- (*pfcrt*-K76, *pfmdr1*-N86, *pfmdr1*-184F, single-copy *pfmdr1*) to 0.5 and on KYF-- to 0.775, we were able to obtain day-28 efficacies of AQ monotherapy on KNF-- at 92.1% and on KYF-- at 76.1%, respectively. An EC50 of 0.65 gives us an estimate for efficacy of AQ monotherapy on TNF-- at 84.7% and an EC50 of 0.82 gave us 71.4% efficacy of AQ monotherapy on TYF--. Since we expected the effect of *pfmdr1*-Y184 on susceptibility to AQ to be weaker than the effect of either *pfcrt*-76T or *pfmdr1*-86Y, we set EC50 of AQ on KNY-- at 0.62 to get an estimate of 86.2% for efficacy of AQ on KNY-- and set EC50 on TNY-- at 0.7 to get an estimate of 81.2% efficacy on TNY--. EC50 of AQ at 0.85 yielded 68.2% efficacy of AQ monotherapy on KYY-- and EC50 at 0.9 yielded 63.3% efficacy of AQ on TYY-- (Table on page 32).

| Genotype | AS | LM | AQ | PPQ | MQ | CQ | AL | ASAQ | DHAPPQ | ASMQ |
|---|---|---|---|---|---|---|---|---|---|---|
| KNY--C1 | 0.689 | 0.719 | 0.862 | 0.899 | 0.945 | 0.810 | 0.915 | 0.962 | 0.972 | 0.983 |
| KNY--C2 | 0.689 | 0.719 | 0.862 | 0.213 | 0.945 | 0.810 | 0.915 | 0.962 | 0.768 | 0.983 |
| KNY--Y1 | 0.241 | 0.719 | 0.862 | 0.899 | 0.945 | 0.810 | 0.795 | 0.896 | 0.929 | 0.962 |
| KNY--Y2 | 0.241 | 0.719 | 0.862 | 0.213 | 0.945 | 0.810 | 0.795 | 0.896 | 0.415 | 0.962 |
| KYY--C1 | 0.689 | 0.828 | 0.682 | 0.899 | 0.945 | 0.639 | 0.953 | 0.905 | 0.972 | 0.983 |
| KYY--C2 | 0.689 | 0.828 | 0.682 | 0.213 | 0.945 | 0.639 | 0.953 | 0.905 | 0.768 | 0.983 |
| KYY--Y1 | 0.241 | 0.828 | 0.682 | 0.899 | 0.945 | 0.639 | 0.878 | 0.772 | 0.929 | 0.962 |
| KYY--Y2 | 0.241 | 0.828 | 0.682 | 0.213 | 0.945 | 0.639 | 0.878 | 0.772 | 0.415 | 0.962 |
| KNF--C1 | 0.689 | 0.627 | 0.921 | 0.899 | 0.945 | 0.810 | 0.890 | 0.977 | 0.972 | 0.983 |
| KNF--C2 | 0.689 | 0.627 | 0.921 | 0.213 | 0.945 | 0.810 | 0.890 | 0.977 | 0.768 | 0.983 |
| KNF--Y1 | 0.241 | 0.627 | 0.921 | 0.899 | 0.945 | 0.810 | 0.723 | 0.948 | 0.929 | 0.962 |
| KNF--Y2 | 0.241 | 0.627 | 0.921 | 0.213 | 0.945 | 0.810 | 0.723 | 0.948 | 0.415 | 0.962 |
| KYF--C1 | 0.689 | 0.719 | 0.761 | 0.899 | 0.945 | 0.639 | 0.915 | 0.930 | 0.972 | 0.983 |
| KYF--C2 | 0.689 | 0.719 | 0.761 | 0.213 | 0.945 | 0.639 | 0.915 | 0.930 | 0.768 | 0.983 |
| KYF--Y1 | 0.241 | 0.719 | 0.761 | 0.899 | 0.945 | 0.639 | 0.795 | 0.825 | 0.929 | 0.962 |
| KYF--Y2 | 0.241 | 0.719 | 0.761 | 0.213 | 0.945 | 0.639 | 0.795 | 0.825 | 0.415 | 0.962 |
| KNYNYC1 | 0.689 | 0.523 | | 0.899 | 0.461 | 0.810 | 0.859 | | 0.972 | 0.846 |
| KNYNYC2 | 0.689 | 0.523 | | 0.213 | 0.461 | 0.810 | 0.859 | | 0.768 | 0.846 |
| KNYNYY1 | 0.241 | 0.523 | | 0.899 | 0.461 | 0.810 | 0.646 | | 0.929 | 0.607 |
| KNYNYY2 | 0.241 | 0.523 | | 0.213 | 0.461 | 0.810 | 0.646 | | 0.415 | 0.607 |
| KYYYYC1 | 0.689 | 0.662 | | 0.899 | 0.461 | 0.639 | 0.897 | | 0.972 | 0.846 |
| KYYYYC2 | 0.689 | 0.662 | | 0.213 | 0.461 | 0.639 | 0.897 | | 0.768 | 0.846 |
| KYYYYY1 | 0.241 | 0.662 | | 0.899 | 0.461 | 0.639 | 0.752 | | 0.929 | 0.607 |
| KYYYYY2 | 0.241 | 0.662 | | 0.213 | 0.461 | 0.639 | 0.752 | | 0.415 | 0.607 |
| KNFNFC1 | 0.689 | 0.422 | | 0.899 | 0.461 | 0.810 | 0.830 | | 0.972 | 0.846 |
| KNFNFC2 | 0.689 | 0.422 | | 0.213 | 0.461 | 0.810 | 0.830 | | 0.768 | 0.846 |
| KNFNFY1 | 0.241 | 0.422 | | 0.899 | 0.461 | 0.810 | 0.570 | | 0.929 | 0.607 |
| KNFNFY2 | 0.241 | 0.422 | | 0.213 | 0.461 | 0.810 | 0.570 | | 0.415 | 0.607 |
| KYFYFC1 | 0.689 | 0.523 | | 0.899 | 0.461 | 0.639 | 0.859 | | 0.972 | 0.846 |
| KYFYFC2 | 0.689 | 0.523 | | 0.213 | 0.461 | 0.639 | 0.859 | | 0.768 | 0.846 |
| KYFYFY1 | 0.241 | 0.523 | | 0.899 | 0.461 | 0.639 | 0.646 | | 0.929 | 0.607 |
| KYFYFY2 | 0.241 | 0.523 | | 0.213 | 0.461 | 0.639 | 0.646 | | 0.415 | 0.607 |
| TNY--C1 | 0.689 | 0.770 | 0.811 | 0.899 | 0.945 | 0.337 | 0.929 | 0.947 | 0.972 | 0.983 |
| TNY--C2 | 0.689 | 0.770 | 0.811 | 0.213 | 0.945 | 0.337 | 0.929 | 0.947 | 0.768 | 0.983 |
| TNY--Y1 | 0.241 | 0.770 | 0.811 | 0.899 | 0.945 | 0.337 | 0.829 | 0.864 | 0.929 | 0.962 |
| TNY--Y2 | 0.241 | 0.770 | 0.811 | 0.213 | 0.945 | 0.337 | 0.829 | 0.864 | 0.415 | 0.962 |
| TYY--C1 | 0.689 | 0.870 | 0.633 | 0.899 | 0.945 | 0.195 | 0.965 | 0.891 | 0.972 | 0.983 |
| TYY--C2 | 0.689 | 0.870 | 0.633 | 0.213 | 0.945 | 0.195 | 0.965 | 0.891 | 0.768 | 0.983 |
| TYY--Y1 | 0.241 | 0.870 | 0.633 | 0.899 | 0.945 | 0.195 | 0.908 | 0.735 | 0.929 | 0.962 |
| TYY--Y2 | 0.241 | 0.870 | 0.633 | 0.213 | 0.945 | 0.195 | 0.908 | 0.735 | 0.415 | 0.962 |
| TNF--C1 | 0.689 | 0.676 | 0.847 | 0.899 | 0.945 | 0.337 | 0.908 | 0.959 | 0.972 | 0.983 |
| TNF--C2 | 0.689 | 0.676 | 0.847 | 0.213 | 0.945 | 0.337 | 0.908 | 0.959 | 0.768 | 0.983 |
| TNF--Y1 | 0.241 | 0.676 | 0.847 | 0.899 | 0.945 | 0.337 | 0.753 | 0.893 | 0.929 | 0.962 |
| TNF--Y2 | 0.241 | 0.676 | 0.847 | 0.213 | 0.945 | 0.337 | 0.753 | 0.893 | 0.415 | 0.962 |
| TYF--C1 | 0.689 | 0.770 | 0.714 | 0.899 | 0.945 | 0.195 | 0.929 | 0.917 | 0.972 | 0.983 |
| TYF--C2 | 0.689 | 0.770 | 0.714 | 0.213 | 0.945 | 0.195 | 0.929 | 0.917 | 0.768 | 0.983 |
| TYF--Y1 | 0.241 | 0.770 | 0.714 | 0.899 | 0.945 | 0.195 | 0.829 | 0.794 | 0.929 | 0.962 |
| TYF--Y2 | 0.241 | 0.770 | 0.714 | 0.213 | 0.945 | 0.195 | 0.829 | 0.794 | 0.415 | 0.962 |
| TNYNYC1 | 0.689 | 0.575 | | 0.899 | 0.461 | 0.337 | 0.869 | | 0.972 | 0.846 |
| TNYNYC2 | 0.689 | 0.575 | | 0.213 | 0.461 | 0.337 | 0.869 | | 0.768 | 0.846 |
| TNYNYY1 | 0.241 | 0.575 | | 0.899 | 0.461 | 0.337 | 0.684 | | 0.929 | 0.607 |
| TNYNYY2 | 0.241 | 0.575 | | 0.213 | 0.461 | 0.337 | 0.684 | | 0.415 | 0.607 |
| TYYYYC1 | 0.689 | 0.719 | | 0.899 | 0.461 | 0.195 | 0.915 | | 0.972 | 0.846 |
| TYYYYC2 | 0.689 | 0.719 | | 0.213 | 0.461 | 0.195 | 0.915 | | 0.768 | 0.846 |
| TYYYYY1 | 0.241 | 0.719 | | 0.899 | 0.461 | 0.195 | 0.795 | | 0.929 | 0.607 |
| TYYYYY2 | 0.241 | 0.719 | | 0.213 | 0.461 | 0.195 | 0.795 | | 0.415 | 0.607 |
| TNFNFC1 | 0.689 | 0.474 | | 0.899 | 0.461 | 0.337 | 0.844 | | 0.972 | 0.846 |
| TNFNFC2 | 0.689 | 0.474 | | 0.213 | 0.461 | 0.337 | 0.844 | | 0.768 | 0.846 |
| TNFNFY1 | 0.241 | 0.474 | | 0.899 | 0.461 | 0.337 | 0.604 | | 0.929 | 0.607 |
| TNFNFY2 | 0.241 | 0.474 | | 0.213 | 0.461 | 0.337 | 0.604 | | 0.415 | 0.607 |
| TYFYFC1 | 0.689 | 0.575 | | 0.899 | 0.461 | 0.195 | 0.869 | | 0.972 | 0.846 |
| TYFYFC2 | 0.689 | 0.575 | | 0.213 | 0.461 | 0.195 | 0.869 | | 0.768 | 0.846 |
| TYFYFY1 | 0.241 | 0.575 | | 0.899 | 0.461 | 0.195 | 0.684 | | 0.929 | 0.607 |
| TYFYFY2 | 0.241 | 0.575 | | 0.213 | 0.461 | 0.195 | 0.684 | | 0.415 | 0.607 |

Since the copy number of *pfmdr1* does not affect the efficacy of AQ monotherapy, EC50 values of AQ on genotypes carrying single and multiple copies of *pfmdr1* are identical.

| Genotype | AS | LM | AQ | PPQ | MQ | CQ | AL | ASAQ | DHAPPQ | ASMQ |
|---|---|---|---|---|---|---|---|---|---|---|
| KNY--C1 | 0.689 | 0.719 | 0.862 | 0.899 | 0.945 | 0.810 | 0.915 | 0.962 | 0.972 | 0.983 |
| KNY--C2 | 0.689 | 0.719 | 0.862 | 0.213 | 0.945 | 0.810 | 0.915 | 0.962 | 0.768 | 0.983 |
| KNY--Y1 | 0.241 | 0.719 | 0.862 | 0.899 | 0.945 | 0.810 | 0.795 | 0.896 | 0.929 | 0.962 |
| KNY--Y2 | 0.241 | 0.719 | 0.862 | 0.213 | 0.945 | 0.810 | 0.795 | 0.896 | 0.415 | 0.962 |
| KYY--C1 | 0.689 | 0.828 | 0.682 | 0.899 | 0.945 | 0.639 | 0.953 | 0.905 | 0.972 | 0.983 |
| KYY--C2 | 0.689 | 0.828 | 0.682 | 0.213 | 0.945 | 0.639 | 0.953 | 0.905 | 0.768 | 0.983 |
| KYY--Y1 | 0.241 | 0.828 | 0.682 | 0.899 | 0.945 | 0.639 | 0.878 | 0.772 | 0.929 | 0.962 |
| KYY--Y2 | 0.241 | 0.828 | 0.682 | 0.213 | 0.945 | 0.639 | 0.878 | 0.772 | 0.415 | 0.962 |
| KNF--C1 | 0.689 | 0.627 | 0.921 | 0.899 | 0.945 | 0.810 | 0.890 | 0.977 | 0.972 | 0.983 |
| KNF--C2 | 0.689 | 0.627 | 0.921 | 0.213 | 0.945 | 0.810 | 0.890 | 0.977 | 0.768 | 0.983 |
| KNF--Y1 | 0.241 | 0.627 | 0.921 | 0.899 | 0.945 | 0.810 | 0.723 | 0.948 | 0.929 | 0.962 |
| KNF--Y2 | 0.241 | 0.627 | 0.921 | 0.213 | 0.945 | 0.810 | 0.723 | 0.948 | 0.415 | 0.962 |
| KYF--C1 | 0.689 | 0.719 | 0.761 | 0.899 | 0.945 | 0.639 | 0.915 | 0.930 | 0.972 | 0.983 |
| KYF--C2 | 0.689 | 0.719 | 0.761 | 0.213 | 0.945 | 0.639 | 0.915 | 0.930 | 0.768 | 0.983 |
| KYF--Y1 | 0.241 | 0.719 | 0.761 | 0.899 | 0.945 | 0.639 | 0.795 | 0.825 | 0.929 | 0.962 |
| KYF--Y2 | 0.241 | 0.719 | 0.761 | 0.213 | 0.945 | 0.639 | 0.795 | 0.825 | 0.415 | 0.962 |
| KNYNYC1 | 0.689 | 0.523 | 0.862 | 0.899 | 0.461 | 0.810 | 0.859 | 0.962 | 0.972 | 0.846 |
| KNYNYC2 | 0.689 | 0.523 | 0.862 | 0.213 | 0.461 | 0.810 | 0.859 | 0.962 | 0.768 | 0.846 |
| KNYNYY1 | 0.241 | 0.523 | 0.862 | 0.899 | 0.461 | 0.810 | 0.646 | 0.896 | 0.929 | 0.607 |
| KNYNYY2 | 0.241 | 0.523 | 0.862 | 0.213 | 0.461 | 0.810 | 0.646 | 0.896 | 0.415 | 0.607 |
| KYYYYC1 | 0.689 | 0.662 | 0.682 | 0.899 | 0.461 | 0.639 | 0.897 | 0.905 | 0.972 | 0.846 |
| KYYYYC2 | 0.689 | 0.662 | 0.682 | 0.213 | 0.461 | 0.639 | 0.897 | 0.905 | 0.768 | 0.846 |
| KYYYYY1 | 0.241 | 0.662 | 0.682 | 0.899 | 0.461 | 0.639 | 0.752 | 0.772 | 0.929 | 0.607 |
| KYYYYY2 | 0.241 | 0.662 | 0.682 | 0.213 | 0.461 | 0.639 | 0.752 | 0.772 | 0.415 | 0.607 |
| KNFNFC1 | 0.689 | 0.422 | 0.921 | 0.899 | 0.461 | 0.810 | 0.830 | 0.977 | 0.972 | 0.846 |
| KNFNFC2 | 0.689 | 0.422 | 0.921 | 0.213 | 0.461 | 0.810 | 0.830 | 0.977 | 0.768 | 0.846 |
| KNFNFY1 | 0.241 | 0.422 | 0.921 | 0.899 | 0.461 | 0.810 | 0.570 | 0.948 | 0.929 | 0.607 |
| KNFNFY2 | 0.241 | 0.422 | 0.921 | 0.213 | 0.461 | 0.810 | 0.570 | 0.948 | 0.415 | 0.607 |
| KYFYFC1 | 0.689 | 0.523 | 0.761 | 0.899 | 0.461 | 0.639 | 0.859 | 0.930 | 0.972 | 0.846 |
| KYFYFC2 | 0.689 | 0.523 | 0.761 | 0.213 | 0.461 | 0.639 | 0.859 | 0.930 | 0.768 | 0.846 |
| KYFYFY1 | 0.241 | 0.523 | 0.761 | 0.899 | 0.461 | 0.639 | 0.646 | 0.825 | 0.929 | 0.607 |
| KYFYFY2 | 0.241 | 0.523 | 0.761 | 0.213 | 0.461 | 0.639 | 0.646 | 0.825 | 0.415 | 0.607 |
| TNY--C1 | 0.689 | 0.770 | 0.811 | 0.899 | 0.945 | 0.337 | 0.929 | 0.947 | 0.972 | 0.983 |
| TNY--C2 | 0.689 | 0.770 | 0.811 | 0.213 | 0.945 | 0.337 | 0.929 | 0.947 | 0.768 | 0.983 |
| TNY--Y1 | 0.241 | 0.770 | 0.811 | 0.899 | 0.945 | 0.337 | 0.829 | 0.864 | 0.929 | 0.962 |
| TNY--Y2 | 0.241 | 0.770 | 0.811 | 0.213 | 0.945 | 0.337 | 0.829 | 0.864 | 0.415 | 0.962 |
| TYY--C1 | 0.689 | 0.870 | 0.633 | 0.899 | 0.945 | 0.195 | 0.965 | 0.891 | 0.972 | 0.983 |
| TYY--C2 | 0.689 | 0.870 | 0.633 | 0.213 | 0.945 | 0.195 | 0.965 | 0.891 | 0.768 | 0.983 |
| TYY--Y1 | 0.241 | 0.870 | 0.633 | 0.899 | 0.945 | 0.195 | 0.908 | 0.735 | 0.929 | 0.962 |
| TYY--Y2 | 0.241 | 0.870 | 0.633 | 0.213 | 0.945 | 0.195 | 0.908 | 0.735 | 0.415 | 0.962 |
| TNF--C1 | 0.689 | 0.676 | 0.847 | 0.899 | 0.945 | 0.337 | 0.908 | 0.959 | 0.972 | 0.983 |
| TNF--C2 | 0.689 | 0.676 | 0.847 | 0.213 | 0.945 | 0.337 | 0.908 | 0.959 | 0.768 | 0.983 |
| TNF--Y1 | 0.241 | 0.676 | 0.847 | 0.899 | 0.945 | 0.337 | 0.753 | 0.893 | 0.929 | 0.962 |
| TNF--Y2 | 0.241 | 0.676 | 0.847 | 0.213 | 0.945 | 0.337 | 0.753 | 0.893 | 0.415 | 0.962 |
| TYF--C1 | 0.689 | 0.770 | 0.714 | 0.899 | 0.945 | 0.195 | 0.929 | 0.917 | 0.972 | 0.983 |
| TYF--C2 | 0.689 | 0.770 | 0.714 | 0.213 | 0.945 | 0.195 | 0.929 | 0.917 | 0.768 | 0.983 |
| TYF--Y1 | 0.241 | 0.770 | 0.714 | 0.899 | 0.945 | 0.195 | 0.829 | 0.794 | 0.929 | 0.962 |
| TYF--Y2 | 0.241 | 0.770 | 0.714 | 0.213 | 0.945 | 0.195 | 0.829 | 0.794 | 0.415 | 0.962 |
| TNYNYC1 | 0.689 | 0.575 | 0.811 | 0.899 | 0.461 | 0.337 | 0.869 | 0.947 | 0.972 | 0.846 |
| TNYNYC2 | 0.689 | 0.575 | 0.811 | 0.213 | 0.461 | 0.337 | 0.869 | 0.947 | 0.768 | 0.846 |
| TNYNYY1 | 0.241 | 0.575 | 0.811 | 0.899 | 0.461 | 0.337 | 0.684 | 0.864 | 0.929 | 0.607 |
| TNYNYY2 | 0.241 | 0.575 | 0.811 | 0.213 | 0.461 | 0.337 | 0.684 | 0.864 | 0.415 | 0.607 |
| TYYYYC1 | 0.689 | 0.719 | 0.633 | 0.899 | 0.461 | 0.195 | 0.915 | 0.891 | 0.972 | 0.846 |
| TYYYYC2 | 0.689 | 0.719 | 0.633 | 0.213 | 0.461 | 0.195 | 0.915 | 0.891 | 0.768 | 0.846 |
| TYYYYY1 | 0.241 | 0.719 | 0.633 | 0.899 | 0.461 | 0.195 | 0.795 | 0.735 | 0.929 | 0.607 |
| TYYYYY2 | 0.241 | 0.719 | 0.633 | 0.213 | 0.461 | 0.195 | 0.795 | 0.735 | 0.415 | 0.607 |
| TNFNFC1 | 0.689 | 0.474 | 0.847 | 0.899 | 0.461 | 0.337 | 0.844 | 0.959 | 0.972 | 0.846 |
| TNFNFC2 | 0.689 | 0.474 | 0.847 | 0.213 | 0.461 | 0.337 | 0.844 | 0.959 | 0.768 | 0.846 |
| TNFNFY1 | 0.241 | 0.474 | 0.847 | 0.899 | 0.461 | 0.337 | 0.604 | 0.893 | 0.929 | 0.607 |
| TNFNFY2 | 0.241 | 0.474 | 0.847 | 0.213 | 0.461 | 0.337 | 0.604 | 0.893 | 0.415 | 0.607 |
| TYFYFC1 | 0.689 | 0.575 | 0.714 | 0.899 | 0.461 | 0.195 | 0.869 | 0.917 | 0.972 | 0.846 |
| TYFYFC2 | 0.689 | 0.575 | 0.714 | 0.213 | 0.461 | 0.195 | 0.869 | 0.917 | 0.768 | 0.846 |
| TYFYFY1 | 0.241 | 0.575 | 0.714 | 0.899 | 0.461 | 0.195 | 0.684 | 0.794 | 0.929 | 0.607 |
| TYFYFY2 | 0.241 | 0.575 | 0.714 | 0.213 | 0.461 | 0.195 | 0.684 | 0.794 | 0.415 | 0.607 |

Wilson, Craig M., Sarah K. Volkman, Sodsri Thaithong, Roger K. Martin, Dennis E. Kyle, Wilbur K. Milhous, and Dyann F. Wirth. 1993. “Amplification of Pfmdr1 Associated with Mefloquine and Halofantrine Resistance in

Plasmodium Falciparum from Thailand." *Molecular and Biochemical Parasitology* 57 (1): 151–60. https://doi.org/10.1016/0166-6851(93)90252-S.

Witkowski, Benoit, Valentine Duru, Nimol Khim, Leila S Ross, Benjamin Saintpierre, Johann Beghain, Sophy Chy, et al. 2016. "A Surrogate Marker of Piperaquine-Resistant *Plasmodium Falciparum* Malaria: A Phenotype–Genotype Association Study." *The Lancet Infectious Diseases*, November. https://doi.org/10.1016/S1473-3099(16)30415-7.

World Health Organization. 1973. "Chemotherapy of Malaria and Resistance to Antimalarials. Report of a WHO Scientific Group [meeting Held in Geneva from 17 to 24 October 1972]." *World Health Organization Technical Report Series* 529: 1–121.

———. 1996. "Assessment of Therapeutic Efficacy of Antimalarial Drugs: For Uncomplicated Falciparum Malaria in Areas with Intense Transmission." Geneva: World Health Organization.

———. 2003. *Assessment and Monitoring of Antimalarial Drug Efficacy for the Treatment of Uncomplicated Falciparum Malaria*. Geneva: World Health Organization.

WWARN AL Dose Impact Study Group. 2015. "The Effect of Dose on the Antimalarial Efficacy of Artemether–Lumefantrine: A Systematic Review and Pooled Analysis of Individual Patient Data." *The Lancet Infectious Diseases* 15 (6): 692–702. https://doi.org/10.1016/S1473-3099(15)70024-1.

WWARN AS-AQ Study Group. 2015. "The Effect of Dosing Strategies on the Therapeutic Efficacy of Artesunate-Amodiaquine for Uncomplicated Malaria: A Meta-Analysis of Individual Patient Data." *BMC Medicine* 13 (1). https://doi.org/10.1186/s12916-015-0301-z.

Yeka, Adoke, Ruth Kigozi, Melissa D. Conrad, Myers Lugemwa, Peter Okui, Charles Katureebe, Kassahun Belay, et al. 2015. "Artesunate/Amodiaquine Versus Artemether/Lumefantrine for the Treatment of Uncomplicated Malaria in Uganda: A Randomized Trial." *Journal of Infectious Diseases*, November, jiv551. https://doi.org/10.1093/infdis/jiv551.

Zani, Babalwa, Michael Gathu, Sarah Donegan, Piero L Olliaro, and David Sinclair. 2014. "Dihydroartemisinin-Piperaquine for Treating Uncomplicated *Plasmodium Falciparum* Malaria." In *Cochrane Database of Systematic Reviews*, edited by The Cochrane Collaboration. Chichester, UK: John Wiley & Sons, Ltd. http://doi.wiley.com/10.1002/14651858.CD010927.

Zongo, Issaka, Grant Dorsey, Noel Rouamba, Christian Dokomajilar, Moise Lankoande, Jean-bosco Ouedraogo, and Philip J. Rosenthal. 2005. "Amodiaquine, Sulfadoxine-Pyrimethamine, and Combination Therapy for Uncomplicated Falciparum Malaria: A Randomized Controlled Trial from Burkina Faso." *The American Journal of Tropical Medicine and Hygiene* 73 (5): 826–32.
